# A 4-bp natural deletion of maize Na^+^/H^+^ exchanger gene alters maize salt stress tolerance

**DOI:** 10.1101/2022.08.14.503892

**Authors:** Meijie Luo, Yanxin Zhao, Yunxia Zhang, Ruyang Zhang, Manjun Cai, Panpan Zhang, Dengxiang Du, Jingna Li, Jinfeng Xing, Xuan Sun, Minxiao Duan, Xiaoduo Lu, Yadong Xue, Ya Liu, Fengge Wang, Baishan Lu, Yuandong Wang, Ronghuan Wang, Wei Song, Jiuran Zhao

## Abstract

Soil salinity is a major environmental constraint severely reducing plant growth and crop productivity worldwide. Knowledge of salt tolerance-related genes can facilitate improving crop salt tolerance and alleviating the threat of increasing saline area to world food security. Here, we identified a major locus *SALT TOLERANCE 1* (*qST1*) conferring maize salt tolerance via bulked segregant RNA-Seq (BSR-Seq). *qST1* encodes a plasma membrane Na^+^/H^+^ exchanger *ZmSOS1* which is the ortholog of *Arabidopsis thaliana SOS1* gene. In salt-sensitive variety D9H, the natural variation of 4-bp deletion in the coding sequence of *ZmSOS1* gene was the causal allele for salt sensitivity. We identified two ethyl methanesulfonate-induced mutants, *zmsos1-1* and *zmsos1-2*, which were sensitive to salt stress and can’t complement salt-sensitive variety under salt stress within an allelism test. Overexpression of *ZmSOS1* enhanced maize seedling salt tolerance. *ZmSOS1* can increase the salt tolerance of *Arabidopsis sos1-1* mutant and can be activated by *AtSOS2* and *AtSOS*3 in yeast cells, suggesting that *ZmSOS1* confers salt tolerance through the conserved SOS signaling pathway in maize. The detrimental allele harboring the 4-bp deletion was rarely found in the natural population but appeared in an important heterotic group composed of x1132x-derived inbred lines which have been used widely for breeding dozens of hybrid varieties in China. The 4-bp deletion-based molecular marker has been successfully used to improve salt-sensitive varieties in a backcross and marker-assisted breeding program by screening and purging this deleterious allele.

## Introduction

Saline soils containing excess Na^+^ are widespread globally and severely affect agricultural production on 800 million ha of arable land on earth (Munns and Tester, 2008; Hasegawa, 2013). Soil salinity in agricultural areas is increasing due to improper agricultural practice, industrial pollution and global climate changes (Ouhibi *et al*., 2014; Ismail and Horie, 2017), that is seriously threatening food security of human beings. Saline soil is characterized by high concentration of soluble Na^+^ and Cl^-^ (Ismail *et al*., 2014), and excess Na^+^ accumulation in plants under saline stress can lead to ionic toxicity, osmotic damage and secondary stresses, particularly oxidative stress (Zhu, 2016; Yang and Guo, 2018a). To survive salt stress, plants have evolved a variety of mechanisms for adaptation to saline habitats, including maintenance of ion homeostasis by removing Na^+^ from the plant cell or sequestering Na+ into the vacuole (Kronzucker and Britto, 2011), and osmotic tolerance through generation and accumulation of osmoprotectants (Maathuis and Amtmann 1999; Zhu, 2003; Pardo *et al*. 2006; Roy *et al*., 2014; Yang and Guo, 2018a). Therefore, exploring molecular mechanisms of plant salt tolerance and developing salt tolerant crops have become important research aims for agricultural scientists and breeders to feed the world (Chinnusamy *et al*., 2005; Munns *et al*., 2006; Munns and Tester, 2008; Roy *et al*., 2014; Ismail and Horie, 2017).

Currently, several Na^+^ selective transporters have been identified in diverse plants based on their important roles in maintenance of Na^+^ ion homeostasis in plant cells (Munns and Tester, 2008; Zhu, 2016; Yang and Guo, 2018a), such as HKT1 Na^+^ transporters (Horie *et al*., 2009; Hamamoto *et al*., 2015), HAK4 Na^+^ transporter (Zhang *et al*., 2019), NHX1 and SOS1 Na^+^/H^+^ antiporters (Kronzucker and Britto, 2011; Ji *et al*., 2013). HKT1 improves salt tolerance by distributing Na^+^ content between root and shoot and reducing Na^+^ accumulation in shoot tissue (Rus *et al*., 2004; Ren *et al*., 2005; Sunarpi *et al*., 2005; Davenport *et al*., 2007; Horie *et al*., 2009; Moller *et al*., 2009). Most of the genes associated with salt tolerance mapped and identified through genome-wide association studies (GWASs) and quantitative trait locus (QTL) analyses are members of the HKT family, including *AtHKT1* in *Arabidopsis thaliana* (Rus *et al*., 2006), *OsSKC1* in *Oryza sativa* (Ren *et al*., 2005), *TmNax1* and *TmNax2* in *Triticum monococcum* (Lindsay *et al*., 2004; Huang *et al*., 2006; James *et al*., 2006a; Byrt *et al*., 2007; Munns *et al*., 2012), *TaKna1* and *TaHKT2;1* in *Triticum aestivum* (Byrt *et al*., 2007; Ariyarathna *et al*., 2016), *SlHKT1;1* and *SlHKT1;2* in *Solanum lycopersicum* (Asins *et al*., 2013), and *ZmHKT1* in maize (Jiang *et al*., 2018; Li *et al*., 2019; Zhang *et al*., 2018). *AtNHX1* encodes a tonoplast membrane Na^+^/H^+^ antiporter which is able to sequester Na^+^ into the vacuole and reduces Na^+^ accumulation in the cytosol. Overexpression of *AtNHX1* in *Arabidopsis* and tomato enhances plant salt tolerance (Apse *et al*., 1999; Zhang and Blumwald 2001). *SOS1*, encoding a Na^+^/H^+^ antiporter at the plasma membrane, is known to be important for ion homeostasis and salt tolerance in plants and it confers salt tolerance under salt stress through the salt overly sensitive (SOS) pathway, which is well documented in *Arabidopsis* and rice and play a key role in modulating the Na^+^ homeostasis in plants (Yang and Guo, 2018a; Zhu, 2016; Martínez-Atienza *et al*., 2007). In *Arabidopsis* under salt stress condition, SOS3 and SCaBP8/CBL10 sense the cytosolic calcium signal induced by salt stress and then, interacts with and activates SOS2, which in turn phosphorylates and activates SOS1, resulting in Na^+^ exclusion from the cytosol to the apoplast and from shoot to root (Shi *et al*., 2002; Zhu, 2016; Yang and Guo, 2018b). *SOS3* and *SCaBP8/CBL10* are nonredundant paralogues and encode EF-hand Ca^2+^ binding proteins. *SOS2* encodes a aserine/threonine protein kinase (Qiu *et al*. 2002; Quan *et al*., 2007). Overexpression of SOS1 or multiple SOS pathway genes can enhance salt tolerance in *Arabidopsis* (Shi *et al*., 2003; Yang *et al*., 2009). Nevertheless, the QTL and genes conferring salt tolerance by protecting plants from Na^+^ toxicity, such as most of the SOS pathway members, investigated in crops are still lacking.

Maize (*Zea mays* L.) is a critically important source of food, feed, energy and forage. Maize also is a glycophyte which is vulnerable to saline stress. Identification and cloning of maize salt tolerance-conferring QTL and genes will benefit molecular improvement of salt-sensitive maize. In this study, we cloned a new salt tolerance-related locus *SALT TOLERANCE 1* (*qST1*) in maize using bulked segregant RNA-Seq (BSR-Seq) and fine mapping, which is identical to the major QTL on maize chromosome 1 for maize salt tolerance mapped previously (Luo *et al*., 2017a; 2019). *qST1* encodes a plasma membrane Na^+^/H^+^ exchanger ZmSOS1. The ethyl methanesulfonate (EMS) mutation analysis and overexpression analysis of *ZmSOS1* confirmed *ZmSOS1* gene is responsible for the *qST1* and essential for coping with salt stress in maize. *ZmSOS1* can suppressed the salt sensitivity of *Arabidopsis sos1-1* mutant and can be activated by *AtSOS2* and *AtSOS3* in yeast cells, suggesting that *ZmSOS1* confers salt tolerance through the conserved SOS pathway existing in maize. The 4-bp deletion in the coding sequence of *ZmST1* gene resulting in frame shift mutation was causal allele for salt sensitivity in salt-sensitive variety and the 4-bp insertion/deletion (InDel) -based molecular marker has been successfully used to improve salt-sensitive varieties by marker assisted selection (MAS) breeding.

## Results

### Salt tolerance of maize Jing724 is controlled by a single dominant locus

Jing724 and D9H were maize sister inbred lines derived from the DuPont-Pioneer hybrid X1132x, namely Xianyu335. X1132x-derived lines (X lines) have formed a novel important heterotic group used for breeding elite hybrid varieties in China, and are particularly used in the heterosis model of X line and Huangzaosi-improved line (HIL) in China, such as Jingke968, Jingnongke728 and Jingke665 (**Table S1**) (Wang *et al*., 2015). A previous maize salt tolerance study indicates that Jing724 was a salt-tolerant line while D9H was a salt-sensitive line (Luo *et al*., 2018b). Here, we re-evaluated salt tolerance differences between Jing724 and D9H under salt stress in field trials and at the germination stage. In field trials, the plant height, biomass and yield of both Jing724 and D9H grown in Tongzhou (TZ, saline soil) were severely reduced compared with those grown in Changping (CP, normal soil) (**Figure S1a-h**). To compare the salt tolerance ability of two different lines, the salt tolerance index (STI) was calculated with the formula, STI = P_salt_ / P_control_×100%, where P_salt_ was the phenotypic value of one trait under salt stress and P_control_ was the phenotypic value under normal condition. The STIs of the traits of D9H examined here were significantly less than those of Jing724, suggesting D9H was sensitive to salt stress (**Figure S1i-l**). For instance, the single ear grain yield STI of D9H was 7.93% ± 2.93% (mean ± standard deviation) while that of Jing724 was 74.17% ± 8.69% (**Figure S1l**). As shown in **Figure 1**, the salt-treatment experiment at the germination stage further confirmed that D9H was more sensitive to salt stress than Jing724 (**Figure 1a-g**). Germinated under 100 mM NaCl for 10 days, D9H had a shorter root length of 3.64 ± 0.23 cm than Jing724 with a root length of 11.74 ± 1.14 cm (**Figure 1c**). We analyzed the root length values of a Jing724×D9H F_2_ population containing 198 individuals under salt stress and found that, based on the root length data, the F_2_ individuals can form two distinct groups, the salt-sensitive group (n=43) with the root length of 0.7–4.8 cm and the salt-tolerant group (n=155) with the root length of 6.6–14.2 cm (**Figure 1h; Table S2**). The ratio of salt-sensitive F_2_ individuals to salt-sensitive individuals agreed very well with the ratio of 1:3 (χ^2^_43:155_ = 1.138, *P* = 0.286), implying that the salt tolerance of Jing724 was controlled by a major dominant locus, designated as *SALT TOLERANCE 1* (*qST1*).

**Figure 1.**
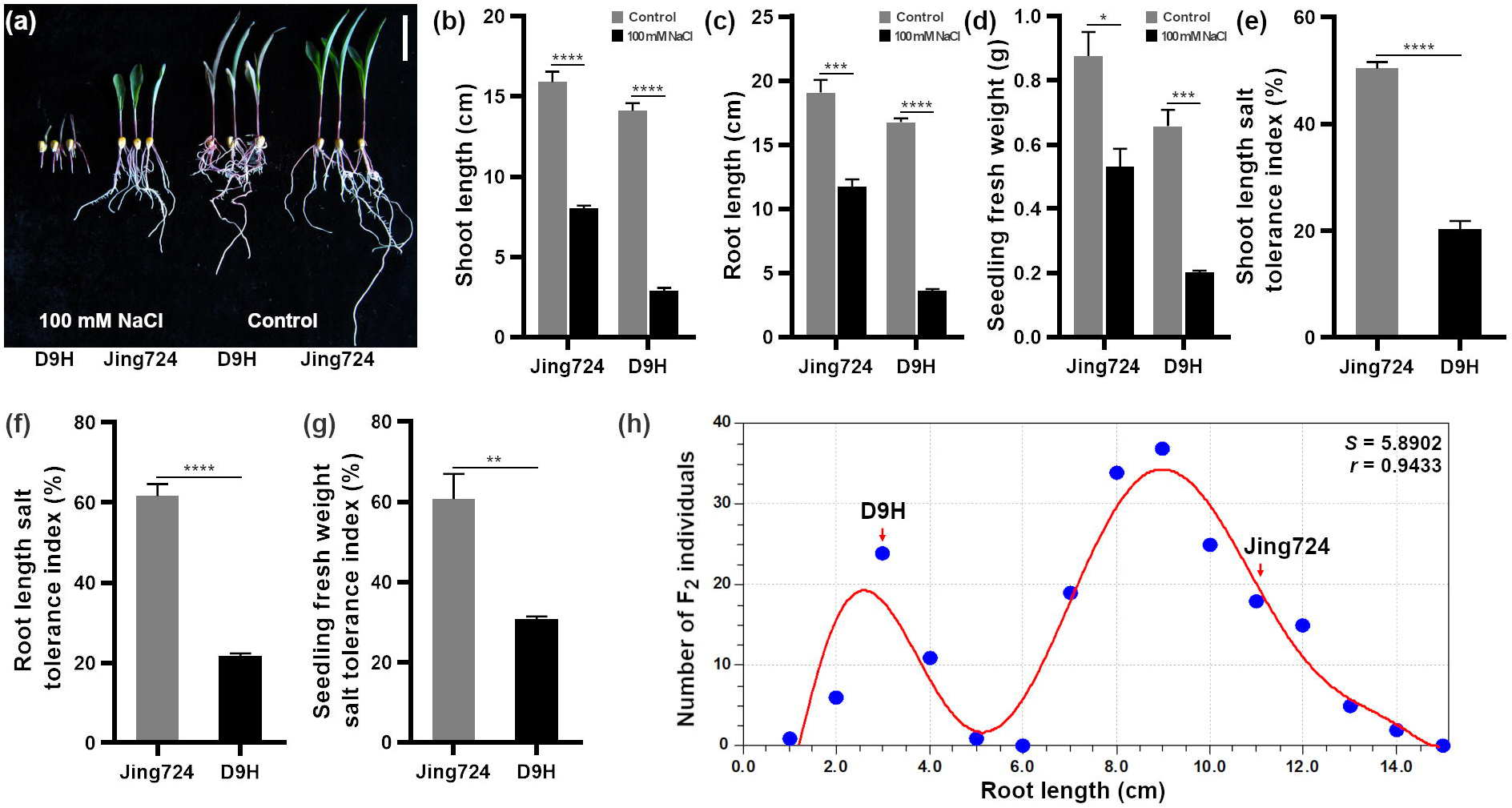
Phenotypes of maize Jing724 and D9H under salt stress. (a) Seedling phenotypes of Jing724 and D9H under salt stress. Jing724 and D9H seeds were germinated under 0 mM (control) and 100 mM NaCl conditions for 10 d and then imaged. Bar = 5 cm. (b-d) Statistical analysis of shoot length (b), root length (c), and seedling fresh weight per plant (d) of Jing724 and D9H as shown in (a). The mean values of four independent experiments were used and five seedlings were measured in each independent experiment. (e-g) Salt tolerance indexes of shoot length (e), root length (f), and seedling fresh weight per plant (g) are calculated as the ratio of phenotypes under 100 mM NaCl treatment to those under the control condition. Data were analyzed using a two-tailed Student’s *t*-test. Error bar = standard error. *, *P* < 0.05. **, *P* < 0.01. ***, *P* < 0.001. ****, *P* < 0.0001. (h) Root length frequency distribution of 198 Jing724 × D9H F_2_ individuals. The Jing724 × D9H F_2_ individuals were germinated under 100 mM NaCl condition for 10 d followed by measuring root length of them. The curve was calculated and displayed by CurveExpert (version 1.4) (http://www.curveexpert.net/), and it revealed two conspicuous groups with 5 cm of root length as the threshold: the salt-sensitive group (n = 43) and the salt-tolerant group (n = 155).

### BSR-seq analysis and fine mapping of the *qST1* locus responsible for maize salt tolerance

To map the *qST1* locus, the BSR-seq was conducted with two pool RNAs from 30 extreme salt-sensitive and 30 extreme salt-tolerant individuals of the Jing724×D9F_2_ population (**Table S2**). A total of 78,463 SNPs were identified by aligning the trimmed reads to the B73 RefGen_v3 reference genome with the GSNAP (Wu *et al*., 2016) and a GATK v4.1.4.0 pipeline (McKenna *et al*., 2010). An empirical Bayesian approach was used to estimate the probability of the linkage between the SNP and the causal locus for maize salt tolerance (Liu *et al*., 2012) and identified 145 associated SNPs clustered on chromosome 1 (**Table S3**). Then a sliding window scan was performed for the associated SNPs (window, 20 SNPs; step, 5 SNPs) to narrow the *qST1* to a reliable interval of 172–183 Mb on chromosome 1 (**Figure 2a; Table S3**), which overlapped the major QTL conferring maize salt tolerance identified previously using Xianyu335 doubled haploid (DH) population (Luo *et al*., 2017; 2019a), implying that they were identical because of the origination of Jing724 and D9H from Xianyu335 (Luo *et al*., 2017; 2019a).

**Figure 2.**
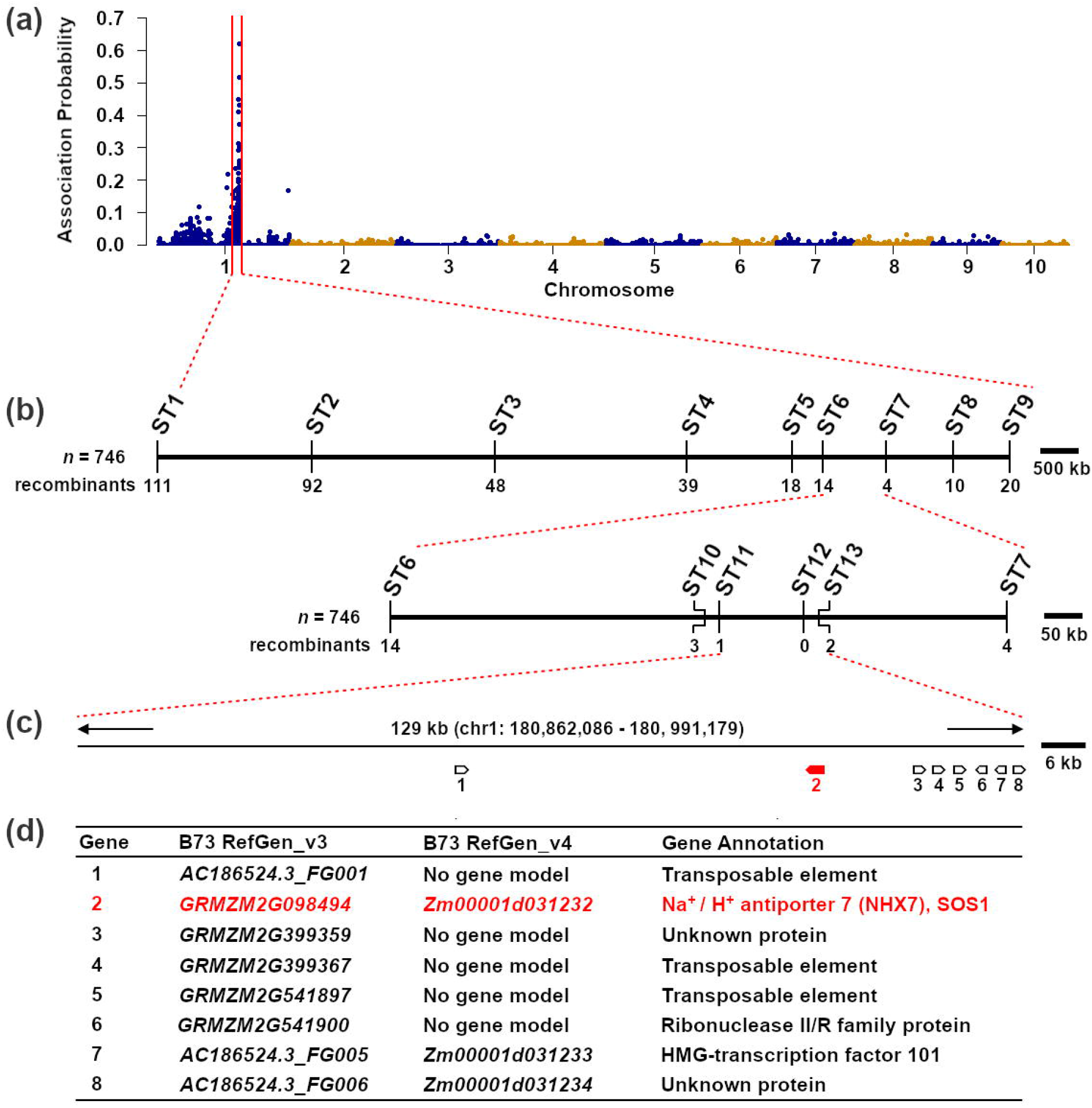
BSR-seq analysis and fine mapping of the *qST1* locus controlling salt tolerance in maize. (a) BSR-seq result. A major QTL *qST1* was mapped on chromosome 1. (b) Fine mapping of the *qST1*. The *qST1* was delimited to a 129-kb interval between KASP markers ST11 and ST13. The genotypes of the 13 KASP markers of all recombinants were shown in **Supplementary Table S4**. (c) Distribution of the candidate genes in the *qST1* QTL interval. The positions of the KASP markers and candidate genes were displayed in (a-c) based on the B73 RefGen_v3 reference genome. (d) Gene functional annotation of the candidate genes in the *qST1* QTL interval. The probable candidate gene *GRMZM2G098494*/*ZmSOS1* is highlighted in red (c, d).

To fine map the *qST1*, we designed 13 SNP-based Kompetitive Allele Specific PCR (KASP) markers in the *qST1* interval (**Table S4**) and used them to genotype 746 extremely salt-sensitive Jing724×D9H F_2_ individuals with a root length of < 4.0 cm. The *qST1* was delimited into a 129-kb region between the markers ST11 and ST13 (**Figure 2b; Table S5**). This region harbored 8 genes annotated in the B73 RefGen_v3 genome, of which only three genes were annotated in the B73 RefGen_v4 genome (**Figure 2d**). Except for *GRMZM2G541897* which putatively was a 196-bp transposon fragment, gene expression of these candidate genes was analyzed using a reverse transcriptase-polymerase chain reaction (RT-PCR), and only *GRMZM2G098494/Zm00001d031232* was expressed in Jing724 and D9H seedlings (**Figure S2**). Gene functional annotation analysis revealed that *GRMZM2G098494/Zm00001d031232* encodes a Na^+^/H^+^ antiporter ZmNHX7 (Kong *et al*., 2021), an ortholog of the *Arabidopsis SOS1* gene in maize, and it is thus also named *ZmSOS1* (Figure 2d). These results indicated that *ZmSOS1* was probably the candidate gene of the *qST1*.

### A 4-bp deletion within *ZmSOS1* gene causes maize D9H sensitive to salt stress

Genomic DNA and complementary DNA (cDNA) sequences of *ZmSOS1* in D9H and Jing724 were obtained by PCR and RT-PCR (**Figure 3a**). The wild-type *ZmSOS1*^*Jing724*^ encodes a 1136-amino acid (aa) protein which shared 59.8% and 81.7% identity with AtSOS1 and OsSOS1, respectively (**Figure S3**). By sequence comparison of *ZmSOS1*^*Jing724*^ and *ZmSOS1*^*D9H*^, we identified six sequence variations in *ZmSOS1* of Jing724 and D9H, including five SNPs in the promoter or introns and one 4-bp InDel in the coding sequence of last exon (**Figure 3a-c**). The 4-bp deletion in *ZmSOS1*^*D9H*^ caused frameshift mutation, resulting in a truncated ZmSOS^D9H^ protein of 1079 aa (**Figure 3c**). It is speculated that the 68 conserved residues at C-terminal end lost in ZmSOS1^D9H^ protein, particular the two residues S1118 and S1120, would be important for ZmSOS1’s function in salt tolerance because their orthologous residues in AtSOS1 were essential for phosphorylation regulation by AtSOS2 (**Figure S3**) (Quintero *et al*., 2011). 5’-and 3’-rapid amplification of cDNA ends (RACE) were used to identify 5’-transcription start site (TSS) and 3’-polyadenylation site of *ZmSOS1* transcripts in D9H and Jing724, respectively. The result indicated that *ZmSOS1*^*D9H*^ and *ZmSOS1*^*Jing724*^ mRNAs had the same 5’-TSS at 121 bp upstream of start codon and the same 3’-polyadenylation site at 141 bp downstream of stop codon. Ten-day-old seedlings of Jing724 and D9H were subjected to 100 mM NaCl treatment and then were used for *ZmSOS1* gene expression analysis and Na^+^ content measurement. The quantitative reverse transcription PCR (qRT-PCR) results indicated that expression of *ZmSOS1* in the root and shoot of Jing724 and D9H seedlings were upregulated after salt treatment (**Figure 3e, f**). In either root or shoot, however, *ZmSOS1*^*D9H*^ and *ZmSOS1*^*Jing724*^ did not show significantly differential expression at each time points after salt treatment (**Figure 3e, f**). Shoot and root Na^+^ contents of D9H and Jing724 seedlings at 0 d, 1 d, 3 d, 5d, and 7 d after salt treatment were examined. At three time points (0 d, 1 d, and 5 d) after salt stress, root Na^+^ content of D9H was significantly higher than that of Jing724, while at each tested time point, D9H had a significantly higher Na^+^ accumulation in shoot than Jing724 (**Figure 3g, h**). After 24-h 100 NaCl treatment, the net Na^+^ and H^+^ flux at root apex of D9H and Jing724 seedlings were measured using the NMT method. The NMT results showed that the net Na^+^ efflux of D9H was significantly lower than that of Jing724 while the net H^+^ efflux of D9H was significantly higher than that of Jing724 (**Figure 3i-l**), suggesting that D9H root had decreased Na^+^ efflux compared with Jing724 under salt stress, leading to higher Na^+^ accumulation in D9H seedlings (**Figure 3g-l**). These findings suggested that the salt sensitivity of D9H was probably due to higher Na^+^ accumulation in shoot caused by the 4-bp deletion in *ZmSOS1*^*D9H*^ which resulted in dysfunction of the truncated ZmSOS1 protein in Na^+^ transport.

**Figure 3.**
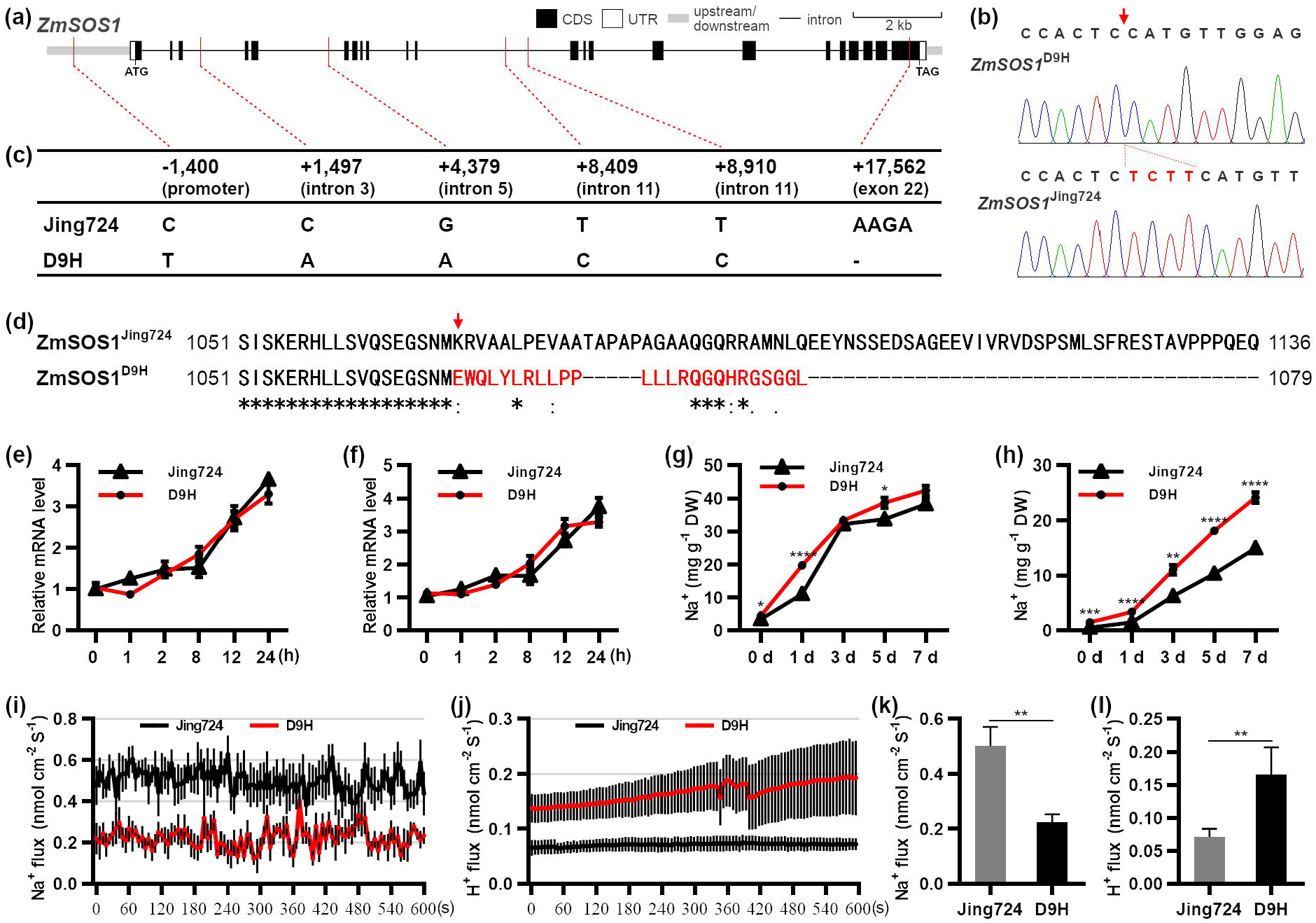
Sequence variation and functional analysis of *ZmSOS1* in Jing724 and D9H. (a) Gene structure of *ZmSOS1*. The genomic DNA and cDNA sequences of *ZmSOS1*^*Jing724*^ confirmed by PCR sequencing were aligned using GSDS2.0 (http://gsds.gao-lab.org/) to build and display the gene structure. The positions of sequence variations were marked by the red line. (b) Validation of the 4-bp deletion variation in D9H. (c) Sequence variations of *ZmSOS1* gene between Jing724 and D9H. The positions of the SNP and InDel variations were shown on the basis of the start codon (NATG, N = -1 and A = +1). (d) Alignment of C-terminal sequences of ZmSOS1^Jing724^ and ZmSOS1^D9H^ proteins. (e, f) Relative expression levels of *ZmSOS1* gene in leaf (e) and root (f) under salt stress. Ten-day-old seedlings grown in 1 × Hoagland buffer were transferred to the 100 mM NaCl-containing 1 × Hoagland buffer and grew for 7 days. The leaf and root RNAs of Jing724 and D9H seedlings were extracted at 0 h, 1h, 2 h, 8 h, 12 h, and 24 h after 100 mM NaCl treatment. The 2^-ΔΔCt^ method was used to analyze qRT-PCR data (Livak and Schmittgen, 2001). Maize *Actin 1* (*GRMZM2G126010*) was used as an internal control. Error bar = standard error, n = 3. (g, h) Shoot and root Na^+^ content of Jing724 and D9H at different times after 100 mM NaCl treatment. DW, dry weight. Error bar = SEM, n = 4. (i,-l) Na^+^ flux (i,j) and H^+^ flux (k,l) in roots of Jing724 and D9H seedlings. Ten-day-old seedlings grown in 1 × Hoagland buffer were subjected to 100 mM NaCl treatment for one day, and then the Na^+^ and H^+^ flux on the root meristem zones of fresh roots was measured for ten minutes using the NMT technique as described in ***Experimental Procedures***. Six biological repeats were conducted for each analysis. Error bar = standard error. Data were analyzed using a two-tailed Student’s *t*-test. *, *P* < 0.05. **, *P* < 0.01. ***, *P* < 0.001. ****, *P* < 0.0001.

### Protein subcellular location and mRNA in situ hybridization of *ZmSOS1* gene

ZmSOS1 protein was predicted to comprise 12 transmembrane domains (**Figure S3**) and to be targeted to the plasma membrane (Kong *et al*., 2021). To determine the subcellular location of ZmSOS1, transient co-expression of ZmSOS1^Jing724^-eGFP and AtCBL1-mCherry fusion proteins in maize protoplast cells was visualized by confocal microscopy and the green and red fluorescence signals were perfectly overlapped on the cell surface. The fluorescence signals in the protoplast cells co-expressing eGFP and AtCBL1-mCherry proteins were not overlapped (**Figure 4a, b**).AtCBL1 was a known plasma membrane protein (Batistic *et al*., 2008). These findings indicated that *ZmSOS1* encodes a plasma membrane protein. In situ hybridization assays were used to detect specific expression of *ZmSOS1* in the different tissues of maize seedlings (**Figure 4c**). Analysis of the conspicuous hybridization signals observed in root tip sections revealed preferential expression of *ZmSOS1* in root epidermal cells and parenchyma cells surrounding the xylem of root tips (**Figure 4d-i**). The shoot transverse sections contained leaf sheath of the first leaf and unexpanded leaf blades of the second and third leaves. The shoot hybridization results indicated that *ZmSOS1* was preferentially expressed in leaf sheath vascular bundle cells and young unexpanded leaves (**Figure 4j, k**).

**Figure 4.**
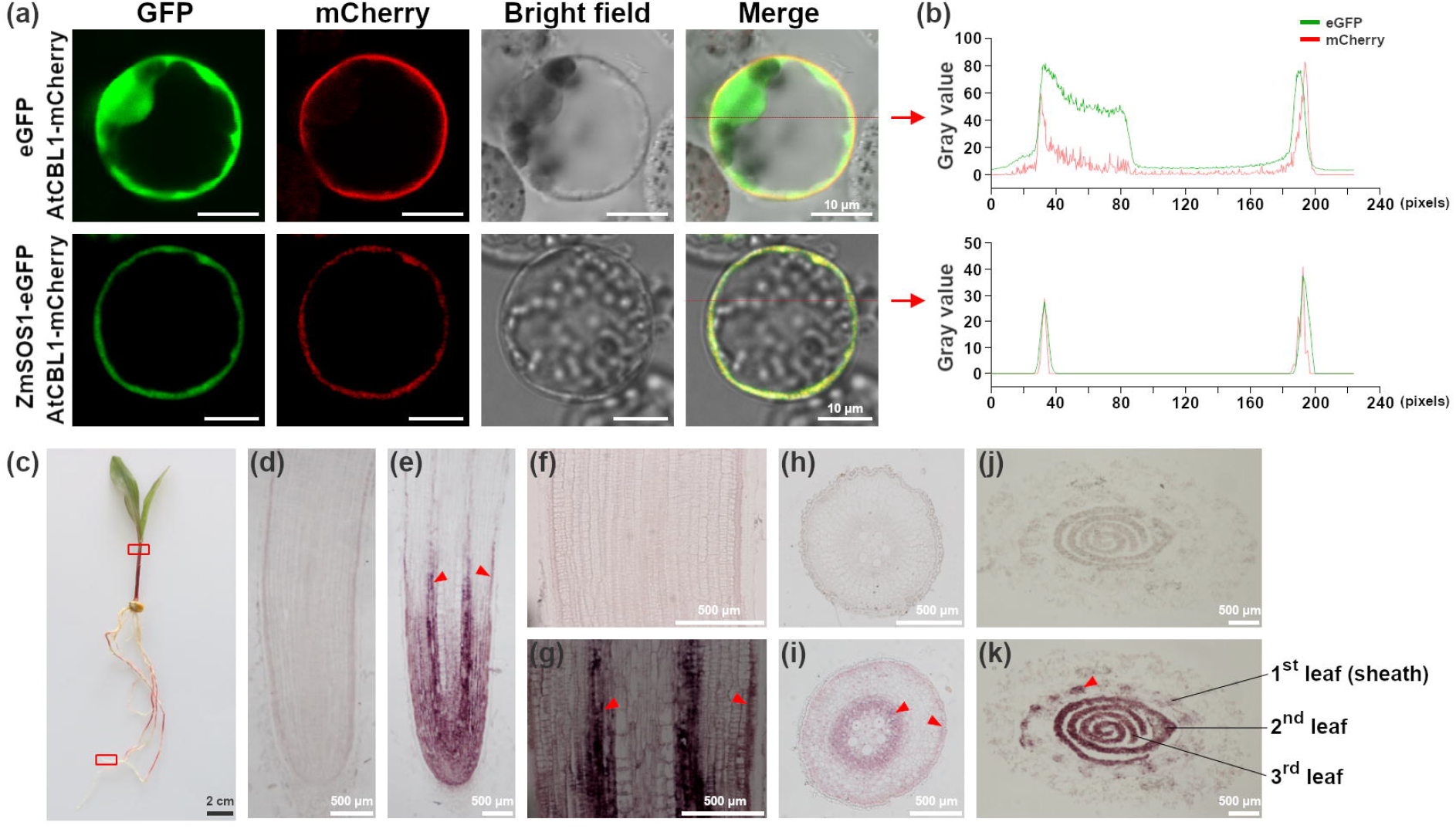
Protein subcellular location and mRNA in situ hybridization of *ZmSOS1* gene. (a) Subcellular localization assay of ZmSOS1 protein in maize protoplasts. ZmSOS1-eGFP fusion protein and AtCBL1-mCherry fusion protein were transiently co-expressed in the maize protoplast cells, and the eGFP fluorescence and the mCherry fluorescence were examined by confocal microscopy. The *Arabidopsis* CBL1 protein was a known plasma membrane protein control (Batistic *et al*., 2008). Transient co-expression of AtCBL1-mCherry fusion protein and eGFP alone protein in the maize protoplast cells was performed as a negative control. (b) Overlapped eGFP and mCherry fluorescence signals of the left-to-right scan lines in Merge images in (a). Green (GFP) and red (mCherry) fluorescence intensities as the gray values were analyzed by ImageJ (https://imagej.nih.gov/). (c-k) mRNA in situ hybridization of *ZmSOS1* in Jing724 seedling root and shoot. (c) The root tip and shoot tissues of 5-day-old Jing724 seedlings used for in situ hybridization assay were marked with red rectangle frames. (d, f, h, j) Hybridizations with a *ZmSOS1* sense probe. (e, g, i, k) Hybridizations with the *ZmSOS1* antisense probe. (d, e) Maize root tips as seen in the longitudinal section. (f, g) Enlarged root tips. (h, i) The transverse section of maize root tips. (j, k) The transverse section of maize shoot at the leaf sheath of the first leaf shown in (c). The striking hybridization signals shown in root tip epidermal cells and parenchyma cells surrounding the xylem (e, g, i) and the vein vascular bundle cells of young seedling leaves (k) were indicated by red arrows.

### Mutant allelism test and overexpression of *ZmSOS1* in maize

We screened the B73 EMS mutant database (http://www.elabcaas.cn/memd/) and obtained two mutants of *ZmSOS1, zmsos1-1* (EMS4-010467) and *zmsos1-2* (EMS4-010486). The point mutation in *zmsos1-1* mutant caused the premature termination of ZmSOS1 protein (**Figure 5a; Figure S4**). In *zmsos1-2* mutant, the point mutation at the splice donor of intron 11 caused the production of aberrant transcripts (**Figure 5a; Figure S4**). The frameshift mutations in the aberrant transcripts resulted in the truncated proteins which only had 169–322 aa identical to the wild-type ZmSOS1 protein (**Figure S4**). In field trials, the growth and development of *zmsos1-1* and *zmsos1-2* mutants grown in TZ were severely inhibited (**Figure S5a**). Plant height and shoot fresh weight of the two mutants were significantly reduced compared with the wild-type B73, and moreover, the mutants failed to produce ears (**Figure S5a-d**). Under 10-d salt stress at the germination stage, the seedling growth of *zmsos1-1* and *zmsos1-2* mutants was also severely inhibited (**Figure 5b**). The root length of the two mutants was significantly less than that of B73 under salt stress or normal conditions (**Figure 5c**). We performed the allelism test by comparing the root length of D9H, D9H×*zmsos1-1*, D9H×*zmsos1-2, zmsos1-1*, and *zmsos1-2* seedlings under salt stress. All these seedlings under salt stress had shorter roots with mean length values of 1.82–3.91 cm, indicating that D9H and the two mutants could not complement each other to the salt tolerance level of the wild-type lines, such as B73 and Jing724 (**Figure 5d**). The root length values of D9H, D9H×*zmsos1-1*, and D9H×*zmsos1-2* seedlings were not significantly different but they were significantly higher than those of *zmsos1-1* and *zmsos1-2*. These results indicated that the causal genes conferring the salt sensitivity of D9H, *zmsos1-1*, and *zmsos1-2* mutants were identical, and also suggested that the *ZmST1*^D9H^ is weak mutation compared with the two EMS mutants (**Figure 5d; Figure S5**).

**Figure 5.**
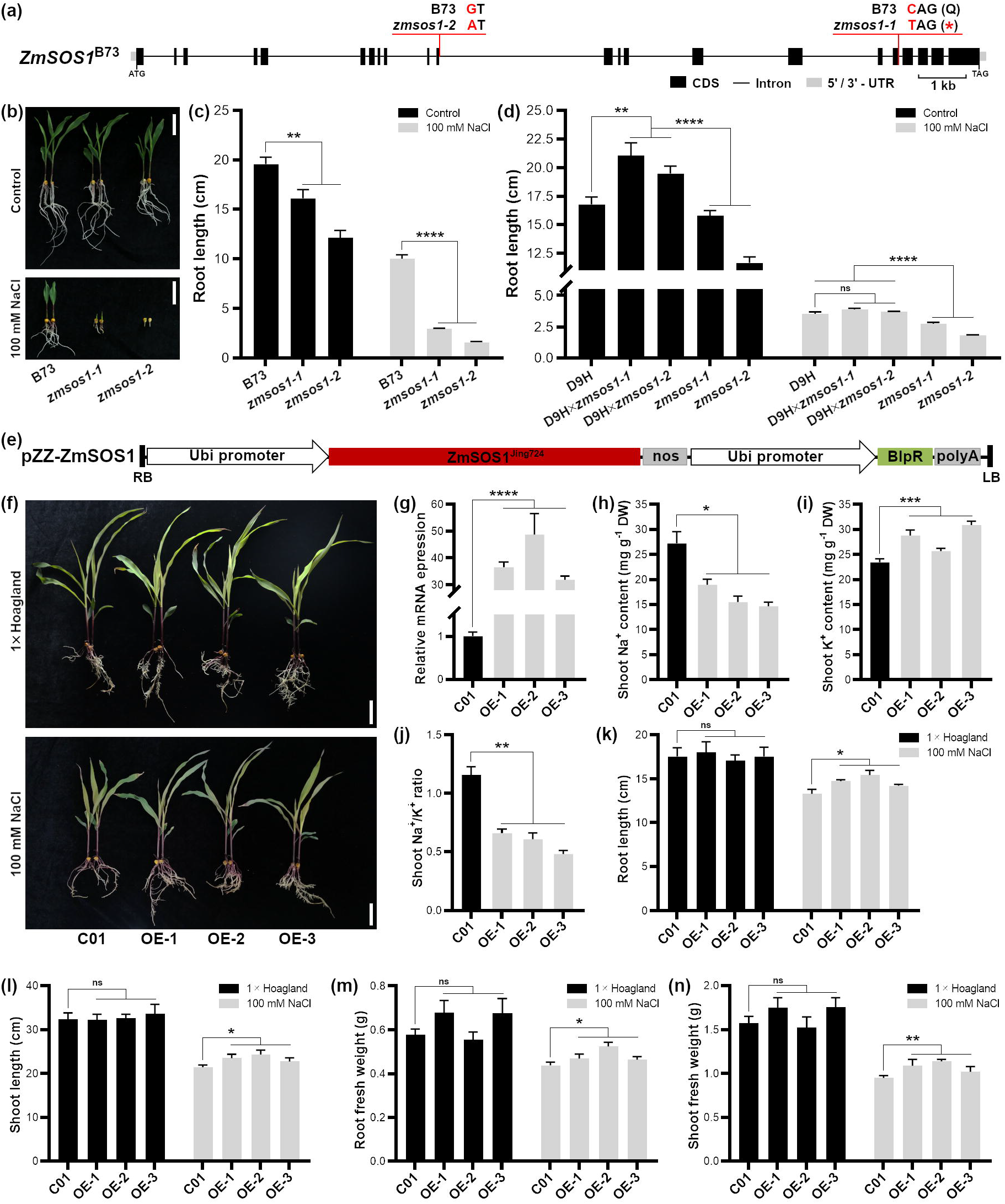
EMS mutant and overexpression analysis of *ZmSOS1*. (a) Point mutations in *zmsos1-1* and *zmsos1-2* mutants. (b) The salt-sensitive phenotype of *zmsos1-1* and *zmsos1-2* mutants. The wild-type B73 and the two *zmsos1* mutants were germinated under 0 mM (control) or 100 mM NaCl condition and imaged after 10 days. (c) Root length of B73, *zmsos1-1*, and *zmsos1-2* seedlings. (d) The allelism test of D9H and the *zmsos1* mutants. Error bar = standard error, n = 4. (e-n) Over-expression of *ZmSOS1*^*Jing724*^ in maize C01 enhanced maize salt tolerance. (e) *ZmSOS1*^*Jing724*^ transgenic vector. (f) Ten-day-old seedlings of C01 and transgenic maize hydroponically grown in 1 × Hoagland buffer were treated with 100 mM NaCl included in 1 × Hoagland buffer and imaged after 7 days. (g) qRT-PCR results of *ZmSOS1* gene expression driven by the maize Ubiquitin promoter in the three *ZmSOS1*^*Jing724*^-overexpressing transformants. Three biological repeats were performed and the 2^-ΔΔCt^ method was used to analyze qRT-PCR data (Livak and Schmittgen, 2001). Maize *Actin 1* (*GRMZM2G126010*) was used as an internal control. (h,i) Shoot Na^+^ (h) and K^+^ (i) content. DW, dry weight. (j) The ratio of shoot Na^+^ content to K^+^ content. (k) Root length. (l) Shoot length. (m) Root fresh weight. (k) Shoot fresh weight. Error bar = standard error, n = 4. Data were analyzed using a two-tailed Student’s *t*-test. *, *P* < 0.05. **, *P* < 0.01. ***, *P* < 0.001. ****, *P* < 0.0001.

To further confirm the salt tolerance ability of *ZmSOS1*, we transformed *ZmSOS1*^*Jing724*^ gene driven by maize ubiquitin promoter into the wild-type maize C01 and generated three independent transgenic lines (**Figure 5e, f**). The higher expression levels of *ZmSOS1* in these transgenic lines were validated by qRT-PCR (**Figure 5g**). Ten-day-old transgenic seedlings were treated with 100 mM NaCl for seven days to evaluate their salt tolerance ability. The *ZmSOS1* transgenic lines exhibited significantly enhanced salt tolerance in seedling growth and biomass production compared with C01. For instance, the shoot fresh weight of *ZmSOS1*-overexpressing lines was significantly higher than that of C01 (*P* = 0.0072) (**Figure 5f-n**). Furthermore, we measured shoot Na^+^ and K^+^ contents of *ZmSOS1* transgenic seedlings after 7-d salt treatment and found that *ZmSOS1* transgenic shoot Na^+^ content was lower than C01 while shoot K^+^ content of the transgenic seedlings was higher than that of C01 (**Figure 5h-j**). The results indicated that over-expression of *ZmSOS1* enhances maize salt tolerance mainly through regulating the long-distance root-to-shoot Na^+^ transport to decrease their shoot Na^+^ content as *AtSOS1* does in *Arabidopsis* (Shi *et al*., 2002).

### Conserved function of *ZmSOS1* involved in the SOS signaling pathway in *Arabidopsis* and yeast

To investigate the conserved role of *ZmSOS1* in *Arabidopsis*, we generated three independent transgenic *Arabidopsis* by transforming *ZmSOS1* gene driven by CaMV 35S promoter into *Arabidopsis sos1-1* (*atsos1-1*) mutant which has a 14-bp deletion in exon 7 (**Figure 6a, b**) (Shi *et al*., 2000). The blank transgenic control was *atsos1-1* mutant transformed with the pMRC35 empty vector and all transgenic *Arabidopsis* were validated by PCR (**Figure 6e**). Five-day-old *Arabidopsis* seedlings grown in MS plates were transferred to 100 mM NaCl-containing MS plates for 14-day salt treatment. The *atsos1-1* mutant and the blank transgenic control exhibited growth arrest and albino leaves under salt stress while all *ZmSOS1* transgenic *Arabidopsis* showed more robust growth (**Figure 6d**). The root elongation of *ZmSOS1* transgenic *atsos1-1* lines was significantly higher than that of *atsos1-1* mutant and the blank transgenic control but was lower than that of the wild-type Col-0, suggesting that *ZmSOS1* could, at least partially, complement the salt-sensitive *atsos1-1* mutant (**Figure 6c-g**).

**Figure 6.**
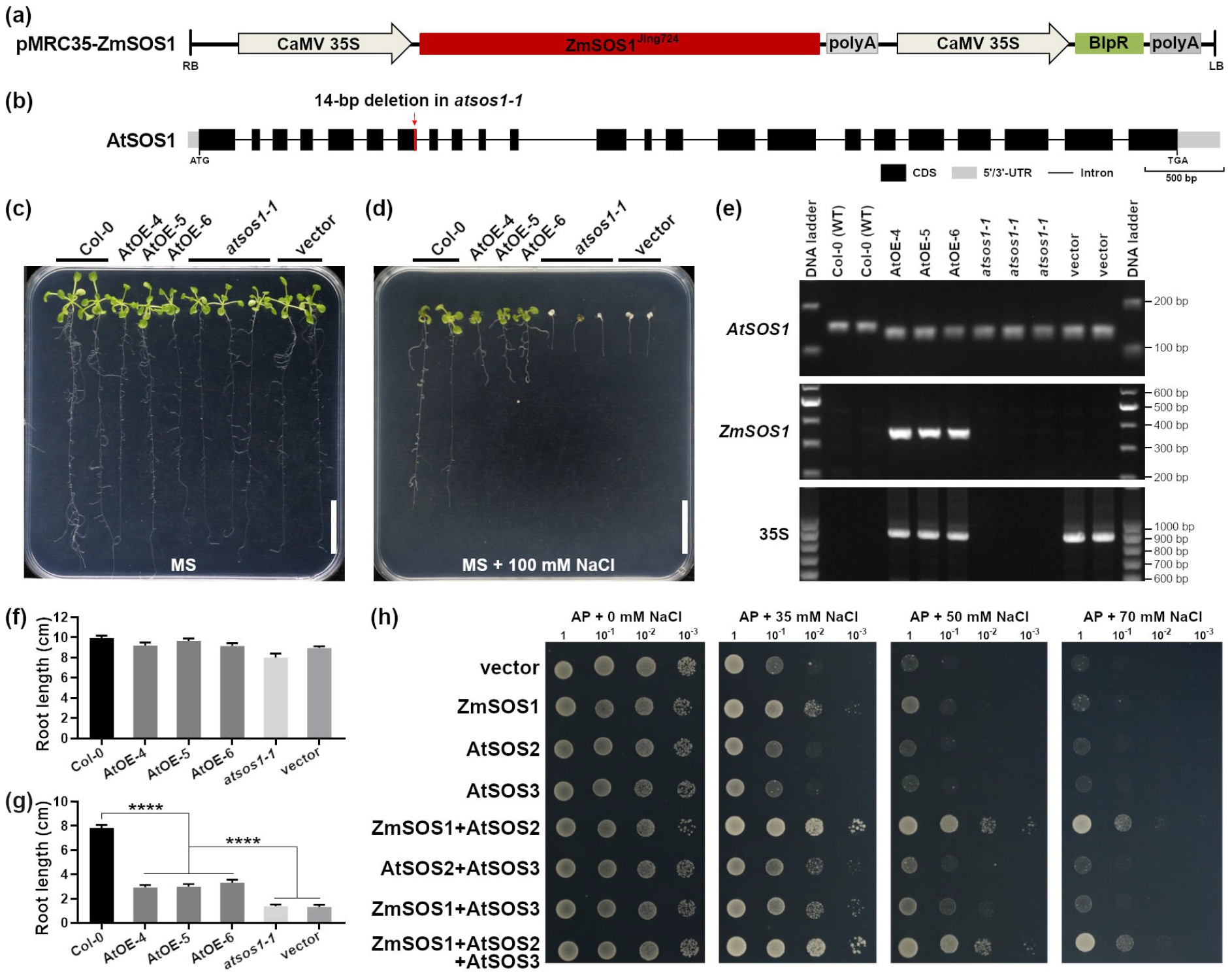
Conserved function of *ZmSOS1* in *Arabidopsis* and yeast. (a) The transgenic vector construction. (b) Gene structure of *AtSOS1*. (c, d) Salt tolerance phenotypes of *ZmSOS1*^*Jing724*^-overexpressing *Arabidopsis* transformants, AtOE-4, AtOE-5, and AtOE-6. *Arabidopsis* seedlings were grown on MS plates for 5 days and then were transferred to MS mediums containing 0 mM (control) or 100 mM NaCl for 14 days in a culture chamber under the conditions of 16h light (22°C) /8h dark (20°C), and light intensity 150 μEm^-2^s^-1^. Bar = 2 cm. (e) PCR detection of *ZmSOS1*^*Jing724*^ transgenic *Arabidopsis*. Primers were listed in **Supplementary Table S11**, including atsos1-1F/R (147/133 bp) for *atsos1-1* mutant, 837F/R (340 bp) for *ZmSOS1*^*Jing724*^ CDS, and 35S-F/R (951 bp) for CaMV 35S promoter of the transgenic vectors. 100-bp DNA Ladder (TransGen Biotech., Beijing, China) was used for defining the PCR product size by electrophoresis on 2% agarose gel. (f,g) Root length of *Arabidopsis* seedlings grown on MS (f) and MS containing 100 mM NaCl (g). Significant differences between means were statistically analyzed by a two-tailed Student’s *t*-test. Error bar = standard error, n = 4. (f) Reconstruction of the SOS pathway in yeast strain AXT3K. The AXT3K strains transformed with *ZmSOS1, AtSOS2*, and *AtSOS3* yeast expression vectors or their combinations were spotted on AP solid medium plates which contain 1 mM KCl and different concentrations of NaCl (0 mM as the control, 35 mM, 50 mM, or 70 mM). From left to right in each panel, the spot concentrations were serially diluted in a 1/10 gradient (from 1 to 10^−3^). The plates were cultivated for 4 days in the dark at 30°C followed by photographing.

In *Arabidopsis*, AtSOS1 was directly activated by AtSOS2 in the SOS signaling pathway under salt stress (Zhu, 2016). To determine the activation of ZmSOS1 protein by the SOS pathway, we reconstructed the SOS system including ZmSOS1, AtSOS2, and AtSOS3 proteins in yeast strain AXT3K deficient in endogenous Na^+^ transporters (Quintero *et al*., 2002). The AXT3K strains transformed with *ZmSOS1, AtSOS2*, and *AtSOS3* and their gene combinations, as well as the empty vector pDR196, were cultivated on an AP solid medium supplemented with 0–70 mM NaCl for salt tolerance assessment. Compared with the stains only harboring pDR196-ZmSOS1 or other vectors, the salt tolerance of the transformant co-expressing *ZmSOS1* and *AtSOS2* and the transformant co-expressing *ZmSOS1, AtSOS2*, and *AtSOS3* were dramatically increased under 50 and 70 mM NaCl treatment (**Figure 6h**), suggesting that the salt-tolerant function of ZmSOS1 was activated by AtSOS2 in these salt-tolerant strains, and implying that ZmSOS1 was probably also activated by ZmSOS2 in an maize SOS pathway as its orthologs do in rice (Martínez-Atienza *et al*., 2007) and *Arabidopsis* (Quintero *et al*., 2002).

### The 4-bp InDel KASP marker is successfully used to improve maize salt tolerance

A KASP marker was designed based on the 4-bp InDel variation between Jing724 and D9H to screen X lines. The 4-bp InDel KASP marker could be used to accurately distinguish different *ZmSOS1* genotypes of 95 Jing724 × F_2_ individuals (**Table S2; Figure S6a**). We used the KASP marker to screen 14 X lines and four HILs and found that seven X lines were homozygous of the 4-bp deletion and the rest were wild-type *ZmSOS1*^*Jing724*^ (**Figure S6e; Table S6**). The salt treatment experiment was performed on nine X lines and the result indicated that five lines harboring *ZmSOS1*^*D9H*^ were more sensitive to salt than those having wild-type *ZmSOS1*^*Jing724*^ (**Figure S7**). Genotype analysis of 45 modern hybrid varieties widely planted in China using the 4-bp InDel KASP marker revealed that 11 hybrid varieties have heterozygous 4-bp deletion allele (**Figure S6f**), of which nine hybrids each have an X-line parent, besides Xianyu335 and Dafeng30 which have the same parental line PH4CV (**Table S7**). Since X lines were derived from the DuPont-Pioneer hybrid Xianyu335, we speculated that the 4-bp deletion of *ZmSOS1* in these hybrid varieties detected here all probably originated from the DuPont-Pioneer maize germplasm.

Using a marker-assisted backcrossing approach (**Figure S8**), the favorable *ZmSOS1* allele was introduced from Jing724 into the salt-sensitive X lines (D9B, D9H, DH382 and MC01) to improve their salt tolerance. Under salt stress in the field trials, the improved lines, D9BST, D9HST, DH382ST and MC01ST, showed dramatically enhanced salt tolerance in plant growth and yield compared with their salt-sensitive versions (**Figure 7**). For example, the single ear grain yield of the improved lines grown in TZ was significantly increased by 1.9 to 25.6 folds compared with their salt-sensitive lines (**Figure 7**). These results suggested that the 4-bp InDel KASP marker can be used to accurately screen salt-sensitive maize and effectively improve its salt tolerance through MAS breeding.

**Figure 7.**
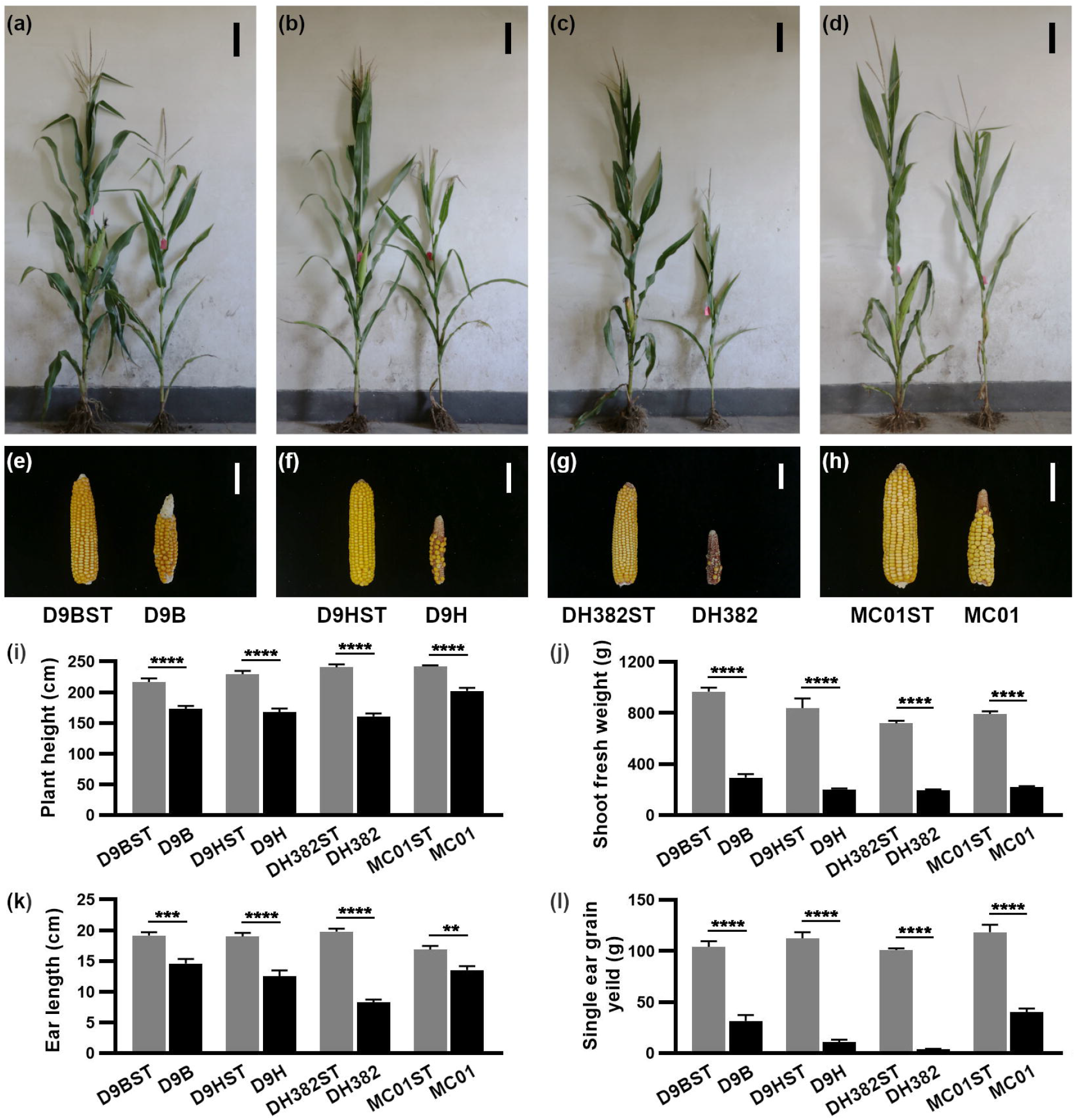
Application of 4-bp InDel-based KASP molecular marker in the improvement of salt-sensitive maize. The salt-sensitive lines (D9B, D9H, DH382, and MC01) and their improved salt-tolerant lines (D9BST, D9HST, DH382ST, and MC01ST) were planted in Tongzhou saline soil on April 25, 2020. (a-d) Plant phenotypes of the salt-sensitive lines and the improved salt-tolerant lines were imaged 10 days after pollination (DAP). Bar = 20 cm. (e-h) Ear phenotypes of the salt-sensitive lines and the improved salt-tolerant lines were imaged after harvesting. Bar = 5 cm. (i) 10-DAP plant height. (j) Shoot fresh weight of 10-DAP plants. (k) Ear length. (l) Single ear grain weight per plant. Data were statistically analyzed using a two-tailed Student’s *t*-test. Error bar = standard error, n = 4. **, *P* < 0.01. ***, *P* < 0.001. ****, *P* < 0.0001.

### The 4-bp deletion in *ZmSOS1*^*D9H*^ is a rare allele in maize natural population

We surveyed the natural population of 489 maize inbred lines, 133 maize landraces and 52 nearest wild relatives of maize (teosintes) with the 4-bp InDel KASP marker to further investigate the source of the 4-bp deletion of *ZmSOS1*. There was no 4-bp deletion allele found in maize landraces and teosintes (**Table S8**), and only two natural population inbred lines, CIMBL50 and CIMBL84, having the 4-bp deletion in *ZmSOS1* were detected (**Figure S6b-d**), suggesting that the 4-bp deletion in *ZmSOS1*^*D9H*^ is a rare allele with the minor allele frequency (MAF) of 0.41% in maize natural population. A comparison of the STI values of CIMBL50, CIMBL84 and Jing724 validated that CIMBL50 and CIMBL84 were salt-sensitive lines (**Figure S9**). The pedigree analysis of CIMBL50 and CIMBL84 showed that their 4-bp deletion allele might originate from the CIMMYT line DTPYC9 (**Figure S9a**).

To mine putative natural variations within *ZmSOS1* conferring different salt tolerance, we performed an association study with the 55 unfiltered SNPs in *ZmSOS1* (**Table S9**) and the phenotypic data of shoot Na^+^ contents of 304 lines (Cao *et al*., 2020) and seedling root lengths of 347 lines under salt stress (Luo *et al*., 2021) (**Table S10**). With the significant cutoff of *P* = 0.018, three SNPs (chr1.S_180935613, chr1.S_180936537, and chr1.S_180941748) significantly associated with shoot Na^+^ content under salt stress and only one SNP (chr1.S_180950792) significantly associated with root length under salt stress were identified (**Figure S10**). Nevertheless, the MAFs of these SNPs were lower than 0.02, ranging from 0.0086 (chr1.S_180950792) to 0.0164 (chr1.S_180935613), and these SNPs were not linked with the 4-bp deletion of CIMBL50 and CIMBL84. These significant SNPs were probably the false-positive results although the phenotype values of two alleles of chr1.S_180950792 showed significant difference (**Figure S10**). These findings could explain why no significant SNPs associated with salt tolerance within the *ZmSOS1*-harboring region on chromosome 1 were identified in the three GWAS analyses performed previously using the similar natural population to this study (Cao *et al*., 2020; Luo *et al*., 2019b; 2021).

## Discussion

The majority of crop species are glycophytes (Horie *et al*., 2009). High Na^+^ concentration in fields seriously impedes crop plant growth, development and production, and threatens future food security (Munns and Tester, 2008). Identification and characterization of the determinants and molecular basis of salt tolerance is essential for breeding and improving salt-tolerant crops (Asins, 2002; Chinnusamy *et al*., 2005; Salvi and Tuberosa, 2005; Ismail and Horie, 2017).

Plant salt tolerance is a complex trait controlled by many QTLs (Ashraf and Foolad, 2012). Most of QTLs for salt tolerance have been cloned in terms of plant Na^+^ content under salt stress as salt tolerance-related trait (Ren *et al*., 2005; Munns *et al*., 2012; Zhang *et al*., 2018; Zhang <u>et al</u>., 2019) since maintaining Na^+^ homeostasis is critical for plant salt tolerance (Zhu *et al*., 2016; Yang and Guo, 2018ab). Here, we cloned a major locus, *qST1*, controlling salt tolerance in maize with root length of seedling under stress as salt tolerance-related trait using the BSR-seq and fine mapping methods (**Figure 2**). The candidate gene *ZmSOS1* encodes a plasma membrane Na^+^/H^+^ antiporter and is orthologous to *AtSOS1* gene. The 4-bp deletion at 3’-end of *ZmSOS1* coding sequence, which caused frameshift mutation and truncated protein, was causal variation for salt sensitivity in D9H maize under salt stress (**Figure 3**). *ZmSOS1* confers maize salt tolerance through reducing shoot Na^+^ content under salt stress. Overexpression of *ZmSOS1* enhanced maize seedling salt tolerance (**Figure 5**). *ZmSOS1* can suppress the salt sensitivity of *Arabidopsis sos1-1* mutant and can be activated by AtSOS2 and AtSOS3 in yeast (**Figure 6**). It is speculated that *ZmSOS1* plays a conserved role in Na^+^ efflux, as *AtSOS1* does in *Arabidopsis*, within the SOS signaling pathway existing in maize. In *Arabidopsis*, the SOS pathway is a complicated regulatory machinery contributing to different responses to salt stress besides ion homeostasis (Ji *et al*., 2013). Besides the SOS core members *SOS1, SOS2, SOS3* and *SCaBP8/CBL10*, many regulatory factors are involved in SOS pathway through interacting with SOS members, such as ABI2 (Ohta *et al*., 2003), 14-3-3 (Zhou *et al*., 2015; Yang *et al*., 2019; Tan *et al*., 2016), PKS5 (Yang *et al*., 2019), AtANN4 (Ma *et al*., 2019), GRIK1 (Barajas-Lopez *et al*., 2018), MPK6 (Yu *et al*., 2010) and GIGANTEA (Kim *et al*., 2013). Identification and functional analysis of *ZmSOS* in this study will facilitate cloning and characterizing SOS pathway genes and their regulatory genes in maize thanks to more and more sequenced maize genomes (Jiao et al., 2017; Hufford *et al*., 2021), although *ZmSOS1* was also characterized by another research team several days ago (Zhou *et al*., 2022).

One important aim of mapping and characterizing the QTLs for salt tolerance is to mine favorable natural variations conferring salt tolerance for crop breeding (Yang and Guo, 2018a). The natural variation of 4-bp deletion at 3’-end of *ZmSOS1*^*D9H*^ was rarely found in natural population, of which only two CIMMYT lines harbor this variation site. Previous GWAS analysis with shoot Na^+^ content, seedling root length, and the seedling survival rate after salt treatment as salt tolerance traits using the same natural population didn’t detect any locus significantly associated with salt tolerance in the chromosomal region harboring *ZmSOS1* (Cao *et al*., 2020; Luo *et al*., 2019b; 2021), probably due to exclusion of rare SNPs (MAF < 0.05) in these GWAS analysis (**Figure S10**).

The 4-bp deletion was found in X lines and hybrid varieties planted widely in China (**Figure S6**). X lines has formed a famous heterotic group in China, which has been used to produce over a dozen of elite hybrid varieties approved in China, particularly by crossing X lines and Huangzaosi-derived lines, such as Jingke968, Denghai605 and Jingnongke728 (**Table S1**). Therefore, the 4-bp deletion detected in 9 of 45 hybrid varieties was from X lines which were involved in breeding these hybrids except Xianyu335 and Dafeng30. The deleterious natural variation was rare in natural population and was not found in landrace and teosinte populations investigated here (**Figure S6**), implying that the 4-bp deletion presumably occurs in an unknown maize in North America and is reserved in these materials which could not undergo rigorous evaluation and selection under salt stress.

Although the salt sensitivity phenotype is recessive and the hybrids, generated by crossing one salt tolerant line and one salt sensitive line, show salt tolerance, the hybrid seed production will be reduced under salt stress when one parent of the hybrid is a salt sensitive line, especially as the maternal parent, since the majority of hybrid seeds are produced in northwest China where saline soils are widespread (Li *et al*., 2014; Xu, 2004). Hence, the deleterious allele should be excluded during hybrid variety breeding and hybrid seed production, in particular, to avoid the situation that two parents of one hybrid variety both possess the deleterious alleles, which will cause significant losses of crop yield under salt stress. Four X lines D9H, D9B, MC01 and DH382 were successfully improved for salt tolerance using the 4-bp InDel KASP marker to remove the deleterious salt sensitive allele from them within a backcrossing MAS breeding program (**Figure 7; Figure S8**), suggesting that the 4-bp InDel KASP marker can be used to screen maize inbred lines in future to remove the salt-sensitive allele, particularly in CIMMYT, DuPont-Pioneer and X line germplasms.

In conclusion, we cloned and characterized a major locus for salt tolerance with BSR-seq analysis in maize, which encodes a plasma membrane Na^+^/H^+^ antiporter and plays a critical role in maintaining Na^+^ homeostasis in maize under salt stress probably via the SOS signal pathway. The deleterious natural variation of 4-bp deletion causing salt sensitivity can be removed from salt sensitive lines to improve salt tolerance by the MAS breeding. This study lays the foundation for deciphering the SOS pathway in maize and provides an insight into molecular breeding applications of natural variations of salt tolerant genes.

## Experimental procedures

### Plant Materials and Phenotyping

X lines including Jing724 and D9H were selectively bred and provided by Maize Research Center, Beijing Academy of Agriculture and Forestry Sciences (BAAFS). The Jing724×D9H F_2_ population was generated by crossing Jing724 and D9F followed by self-crossing and was used for BSR-seq analysis and fine mapping of *qST1*. In field trials, each line of 20 plants was grown in a 60 × 500-cm plot at Changping (CP, N40°10′50.38″, E116°27′15.40″ in Google Earth™) and Tongzhou (TZ, N39°41′49.70″, E116°40′50.75′) experimental stations of Maize Research Center of BAAFS with four replicates in a randomized block design. TZ field soil was moderately salinized with the Na^+^ content (0–40 cm layer) of 123.5 ± 16.1 mg/kg dry weight (DW) and CP field soil was the control with the Na^+^ content of 26.5 ± 1.1 mg/kg DW (**Luo et al**., **2017**). It should be noted that the planting times in fields ranging from April 20 to May 20 were used in this study since the severe salt stress on maize planted in TZ later or after May would be alleviated due to massive rainfall in summer (data not shown). The salt stress experiments at germination and seedling stages of maize were performed using a specific maize seedling identifying apparatus (**Luo *et al***., **2018a; 2018b; 2019; 2021**). Briefly, for germination stage, maize seeds were germinated in 100 mM NaCl solution and sterile water (control) and seedling phenotypes were examined after 10 days. For seedling stage, ten-day-old seedlings grown in 1 × Hoagland’s solution were transferred to 1×Hoagland’s solution supplemented with 0 mM NaCl (Control) and 100 mM NaCl, and were cultivated for 7 days followed by phenotyping. The fresh weight of root and seedling excluded the seed weight.

### Na^+^ and K^+^ content measurement

Soil sampling and measurement of the Na^+^ content in the 0–40 cm soil layer of CP and TZ fields were performed in May, 2017as described previously (**Luo et al**., **2017**). Ten-day-old seedlings of Jing724, D9H, C01, and transgenic maize grown in 1 × Hoagland’s solution were treated with 100 mM NaCl for 7 days. Shoot and root tissues of Jing724 and D9H were collected after different durations (0–7 d) of salt stress treatment to measure their Na^+^ contents. After 7-d salt treatment, shoot Na^+^ and K^+^ contents of C01 and transgenic maize were determined. Dried seedling root and shoot were grounded and filtered by the 40-mesh sieve and 0.2-g powder was digested at 180°C for 4 h in a 50-ml tube with 10 ml nitric acid and 2 ml hydrogen peroxide. Sterile water was added to a final weight of 50 g. The Na^+^ and K^+^ contents were determined by a Dionex ICS-600 ion chromatographic system (Thermo Fisher Scientific, MA, USA) with the Dionex IonPac AS14 column (Thermo Fisher Scientific, MA, USA).

### DNA and RNA isolation

Genomic DNAs of maize and *Arabidopsis* plants were extracted using the CTAB method (Abdel-Latif and Osman, 2017). Total RNAs of different tissues were isolated using Trizol^®^ reagent (Lifescience, CA, USA) following the manufacturer’s instructions.

### BSR-seq analysis and fine mapping of *qST1*

The Jing724 × D9H F_2_ population was used for the BSR-seq analysis. The F_2_ individuals were germinated in 100 mM NaCl solution and after 10 days, their root lengths were measured. Based on the values of root length, the F_2_ population was divided into two distinct groups, the salt-sensitive group including short-root individuals and the salt-tolerant group comprising long-root individuals (**Figure 1h**). The total RNAs of 30 extremely salt-sensitive and 30 extremely salt-tolerant F_2_ individuals were equally mixed to form the salt-sensitive and salt-tolerant RNA pools, respectively. The RNA-seq data of the two RNA pools were produced by the Illumina HiSeq X Ten platform (**Illumina, CA, USA**). SNP calling and identification of significant SNPs associated with the root length of seedlings under salt treatment were performed as described previously (**Liu et al**., **2012; Tang et al**., **2021**). The probability cutoff of 0.05 was used for identification of significantly associated SNPs (**Liu *et al***., **2012**).

To fine map the *qST1* locus, the KASP markers in the *qST1* interval were designed based on the SNPs identified in the above RNA-seq data and the genomic DNA re-sequencing data of Jing724 and D9H reported previously (**Luo *et al***., **2018b**). A large Jing724×D9H F_2_ population of over 3200 individuals was generated and of them, 746 salt-sensitive F_2_ individuals (root length < 4.0 cm under salt stress) were used to identify the recombination events of the KASP markers flanking the *qST1* locus, to narrow the *qST1* region. In the Jing724×D9H F_2_ population, theoretically, the *qST1* genotype of the salt-sensitive individuals should be the same as that of D9H. All primers used in the fine mapping of *qST1* were designed and synthesized by LGC Biosearch Technologies (UK) and were included in **Supplementary Table S4**. The genotypes of the F_2_ individuals were determined using the KASP method as described below.

### Kompetitive allele specific PCR (KASP) assay

The KASP method was used to fine map the *qST1* and screen the 4-bp deletion of *ZmSOS1* in teosintes, maize landraces, inbred lines, hybrid varieties and Jing724 × D9H F_2_ individuals. The KASP primers were designed and synthesized by LGC Genomics (London, UK) and were shown in **Supplementary Table S4**. The high-throughput SNP genotyping was performed using the LGC SNPline PCR Genotyping System following standard experimental procedures of the Laboratory of the Government Chemistry (LGC Genomics) as described previously (Tang *et al*., 2021). A total of 1.5 μl DNA of each sample was pipetted into 96-, 384-or 1536-well PCR plates and then dried at 55°C. The KASP genotyping reactions were carried out in a total volume of 1 μl, including 0.014 μl KASP Assay mix (72×), 0.5 μl KASP Master Mix (2×) and 0.486 μl DNase/RNase-free deionized water. The PCR reactions were performed in a water bath as the following procedure: 94°C for 15min, ten touchdown cycles (94°C for 20s; 61—55°C for 1 min, 0.6°C decrease in each cycle), and then further 26 cycles (94°C for 20 s, 55°C for 1 min). The fluorescence signal in each KASP reaction was detected by the Pherastar SNP genotyping detector (BMG Labtech, Ortenberg, Germany). The genotyping data were analyzed and displayed by the Kraken software (LGC Genomics).

### RT-PCR and qRT-PCR

Total RNAs from shoot and root of maize and *Arabidopsis* were isolated using the TRIzol reagent (Thermo Fisher Scientific) according to the manufacturer instructions (Zhang *et al*., 2017). cDNA synthesis and qRT-PCR were performed described previously (Tang *et al*., 2021). Primers used for RT-PCR and qRT-PCR analysis were listed in **Supplementary Table S11**. qRT-PCR data were analyzed using the 2^−ΔΔCt^ method (Livak and Schmittgen, 2001) with three biological replicates and maize *Actin1* (*GRMZM2G126010*) gene was used as an internal control.

### 5’ and 3’ RACE

5’-and 3’-ends of *ZmSOS1*^*Jing724*^ and *ZmSOS1*^*D9H*^ mRNAs were identified by 5’ and 3’ RACE experiments, respectively, using the the FirstChoice™ RLM-RACE Kit (Thermo Fisher Scientific) as described previously (Tang *et al*., 2021). The RACE PCR products were inserted into the pEASY Blunt Simple vector (TransGen Biotech, Beijing, China) and then ten clones of each experiment were sequenced by Tianyihuiyuan BioTech (Beijing, China).

### Subcellular localization

The *ZmSOS1*^*Jing724*^ coding sequence was amplified and integrated into pM999-eGFP vector at *Xba*I site to express the fused protein ZmSOS1-eGFP driven by the cauliflower mosaic virus (CaMV) 35S promoter. RT-PCR product of *Arabidopsis CBL1* coding sequence was inserted into pM999-mCherry vector at *Xba*I site to construct the pM999-AtCBL1-mCherry vector. Primes used for vector construction were listed in **Supplementary Table S11**. The pM999-ZmSOS1-eGFP and pM999-AtCBL1-mCherry vectors were co-transformed into maize Jing724 leaf protoplasts using the plant protoplast preparation and transformation kit (ZhongkeRuitai Biotech., Beijing, China). Additionally, pM999-eGFP and pM999-AtCBL1-mCherry vectors were co-transformed into Jing724 leaf protoplasts as a control. The subcellular localization of GFP and mCherry fluorescence signals were investigated with the SP8 confocal microscope (Leica, NC, USA). The subcellular localization assay was performed with three repeats and similar results were observed, and the representative images were presented in **Figure 4a**.

### In situ hybridization

Root tip and shoot of five-old-day Jing724 seedlings grown in 1 × Hoagland’s solution were collected for in situ hybridization assay as shown in **Figure 4c**. Sample preparation, RNA hybridization, and immunologic detection of hybridization signals were conducted as described previously (Sun *et al*., 2018). A 432-bp fragment in exon 22 of ZmSOS1 was amplified with the primer pairs, SOS1-F1/SOS1-T7-R1 and SOS1-T7-F1/SOS1-R1 (**Table S11**), where SOS1-T7-R1 and SOS1-T7-F1 have a T7 promoter sequence at their 5’-end. The PCR products were purified and used to *in vitro* synthesize digoxigenin-labeled sense and antisense probes using T7 RNA polymerases (Roche Diagnostics, Mannheim, Germany). *ZmSOS1* sense probe was used as the negative control.

### Non-invasive micro-test technology

The non-invasive micro-test technology (YoungerUSA LLC, MA, USA) was utilized to measure the net Na^+^ and H^+^ fluxes of Jing724 and D9H root tips. Ten-day-old seedlings grown in 1 × Hoagland’s solution were transferred to 1×Hoagland’s solution supplemented with 100 mM NaCl and incubated for 24 h. These roots after 100 mM NaCl treatment were incubated in the measuring solution (0.1 mM KCl, 0.1 mM CaCl_2_, 0.1 mM MgCl_2_, 0.5 mM NaCl, 0.3 mM MES, 0.2 mM Na_2_SO_4_, pH 6.0) to balance for 10 min, then the Na^+^ and H^+^ net fluxes were measured at fresh root meristem zone (∼500 μm from the root tip) using the NMT Physiolyzer^®^ system (YoungerUSA LLC) with the Na^+^-and H^+^-selective microsensors, respectively, which were prepared as described previously (Fan *et al*., 2019). The Na^+^ or H^+^ net fluxes were calculated using imFluxes v2.0 (YoungerUSA LLC). Six seedlings were measured for each sample and each seedling served as one replicate for statistical analysis.

### EMS mutant analysis and allelism test

To obtain null mutations of *ZmSOS1* gene, we used the gene model *Zm00001d031232* to query the B73 EMS mutant database (http://www.elabcaas.cn/memd/). Two EMS mutants *zmsos1-1* (EMS4-010467) and *zmsos1-2* (EMS4-010486) were identified and their point mutation sites were validated by PCR sequencing with the primers listed in **Supplementary Table S11**. The point mutation in *zmsos1-1* mutant caused a premature stop codon in exon 18. In *zmsos1-2* mutant, an aberrant splice donor site (GT-to-AT mutation) of intron 11 caused incorrect intron splicing events. Salt-sensitive phenotypes of the two mutants were examined in the field trial and the salt stress experiment at the germination stage as described above. The D9H×*zmsos1-1* and D9H×*zmsos1-2* F_1_ seeds were produced by crossing D9H with *zmsos1-1* and *zmsos1-2* in CP normal field. Seedling root length of D9H, D9H×*zmsos1-1*, D9H×*zmsos1-2, zmsos1-1* and *zmsos1-2* germinated and grown under 100 mM NaCl condition for 10 days was compared to prove that the salt-sensitive causal gene of D9H was allelic to *ZmSOS1* gene mutated in *zmsos1-1* and *zmsos1-2*.

### Overexpression of *ZmSOS1* gene in maize

The coding sequence of *ZmSOS1*^*Jing724*^ was integrated into the binary vector pZZ (Luo *et al*., 2022) to create the overexpression construct pZZ-ZmSOS1^Jing724^ under the control of the Ubiquitin promoter (Figure 5e). pZZ-ZmSOS1^Jing724^ was introduced into maize inbred line C01 with an *Agrobacterium*-mediated immature embryo transformation method as described previously (Luo *et al*., 2022). Three independent T_0_ transgenic events (OE-1, -2 and -3) were obtained and the homozygous T_2_ transgenic plants were selected by glufosinate ammonium and PCR amplification of transgenic *ZmSOS1*^*Jing724*^ coding sequence which was intronless. High-level expression of *ZmSOS1* mRNA in transgenic plants was validated by qRT-PCR assay with the primer pair SOS1qF1/SOS1qR1 (**Table S11**). Ten-day-old seedlings of C01 and the T_2_ transformants grown in 1×Hoagland’s solution were treated with 100 mM NaCl and after seven days their salt tolerance-related phenotypes were examined. The salt tolerance tests were performed four times and the representative images were presented in **Figure 5f**.

### *Arabidopsis sos1-1* mutant complementation assay

Full-length coding sequence of *ZmSOS1*^*Jing724*^ was amplified from Jing724 seedling cDNAs with the primers B-11F/B-17905R (**Table S11**), and the PCR product was inserted into the *Bam*HI cloning site of the binary vector pMRC35 with *BlpR* as the selection marker gene (**Figure 6a**), resulting in the vector pMRC35-ZmSOS1^Jing724^, where expression of *ZmSOS1*^*Jing724*^ was driven by CaMV 35S promoter. pMRC35-*ZmSOS1*^*Jing724*^ and empty vector pMRC35 were introduced into *Agrobacterium* GV3101 strain. Then, the resulting *Agrobacterium* clones were used to transform *atsos1-1* mutant with the floral dip method (Zhang *et al*., 2006). The homozygous T_2_ transgenic plants were screened by glufosinate ammonium and PCR with *ZmSOS1*^*Jing724*^-specific primers. Three independent transformants (AtOE-4, -5 and -6) were subjected to complementation tests. *Arabidopsis* seedlings were grown on MS plates for 5 days and then were transferred to MS mediums supplemented with 0 mM and 100 mM NaCl for 14 days in a culture chamber under the conditions of 16h light (22°C) /8h dark (20°C), and light intensity 150 μEm^-2^s^-1^. Similar results were observed in four independent experiments and the representative pictures were presented in **Figure 6**.

### Construction of the SOS pathway in yeast

The coding sequences of *ZmSOS1*^*Jing724*^, *AtSOS2* (*At5g35410*) and *AtSOS3* (*At5g24270*) were cloned into the *Spe*I and *Xho*I cloning sites of the yeast expression vector pDR196 using the ClonExpress^®^ II One Step Cloning Kit (Vazyme Biotech, Nanjing, China) to construct pDR196-ZmSOS1, pDR196-AtSOS2 and pDR196-AtSOS3, respectively. The *AtSOS2* and *AtSOS3* coding sequences were cloned in p404MET25 at the *Spe*I and *Xho*I sites to create the expression cassettes MEF17promoter-AtSOS2-CYC1terminator and MEF17promoter-AtSOS3-CYC1terminator, respectively. Then, the MEF17promoter-AtSOS2-CYC1terminator cassette was amplified and integrated into the *Kas*I site of pDR196-ZmSOS1 to generate pDR196-ZmSOS1-AtSOS2. At the same time, the MEF17promoter-AtSOS3-CYC1terminator cassette was amplified and cloned in pDR196-ZmSOS1 and pDR196-AtSOS2 at their *Kas*I site, resulting in pDR196-ZmSOS1-AtSOS2 and pDR196-AtSOS2-AtSOS3, respectively. The methods and primers used for yeast vector construction are shown in **Supplementary Table S12**. The vector sequences were shown in **Supplementary Dataset S1**. The pDR196 empty vector (as a blank control), pDR196-ZmSOS1, pDR196-AtSOS2, pDR196-AtSOS3, pDR196-ZmSOS1-AtSOS2, pDR196-ZmSOS1-AtSOS2 and pDR196-AtSOS2-AtSOS3 were transformed into the yeast strain AXT3K (Δ*ena1*::*HIS3*::*ena4*, Δ*nha1*::*LEU2*, Δ*nhx1*::*KanMX4*) (Quintero *et al*., 2002) using the polyethylene glycol-lithium acetate method (Elble, 1992). pDR196-ZmSOS1-AtSOS2 and pDR196-AtSOS3 were co-transformed into AXT3K stain to obtain the transformant carrying the three *SOS* genes. The positive transformants were selected by the synthetic dextrose drop-out media (SD/-Ura) and PCR with specific primers (**Table S12**). The salt tolerance tests of these transformants were performed on the solid AP medium (Rodriguez-Navarro and Ramos, 1984) supplemented with 1 mM KCl and with different concentrations of NaCl as required in each experiment. Similar results of three independent experiments were obtained and the representative images are shown in **Figure 6h**.

### Association analysis of *ZmSOS1*

To identify putative genetic variations of *ZmSOS1* gene associated with salt tolerance, the association mapping tests for *ZmSOS1* were performed with the known phenotypic data of shoot Na^+^ content (of 304 lines) and root length (of 347 lines) under salt stress reported previously (Cao *et al*., 2020; Luo *et al*., 2021), and the genotypic data of 55 SNPs in *ZmSOS1* gene retrieved from the 1.06-million SNP data of 368 lines (Fu *et al*., 2013). A mixed linear model was used to perform association analysis in the TASSEL5.2.44 software (Glaubitz *et al*., 2014). The significance threshold of *P* value was 1/n, where n is the number of SNPs.

### MAS breeding for improving salt-tolerant maize varieties

The 4-bp InDel-based KASP marker was used to screen the elite inbred lines which have the *ZmSOS1*^*D9H*^ allele. To improve the salt-sensitive lines (e.g. D9B, D9H, DH382 and MC01), the favorable *ZmSOS1*^*Jing724*^ allele was introgressed from Jing724 into the four salt-sensitive lines using a backcrossing strategy with the salt-sensitive lines as the recurrent parents (**Figure S8**). From generations BC_1_ to BC_5_F_2_, the 4-bp InDel KASP marker was used to screen and select the individuals carrying the *ZmSOS1*^*Jing724*^ allele. Finally, the salt-tolerant individuals harboring homozygous *ZmSOS1*^*Jing724*^ gene were subjected to background selection with 40 pairs of SSR marker primers (Wang *et al*., 2011). D9B, D9H, DH382 and MC01 and their salt-tolerant counterparts (D9BST, D9HST, DH382ST and MC01ST) were planted in TZ saline field on April 25, 2020 for salt tolerance assessment of them.

### Statistical analysis

Two-tailed student’s test was conducted with Office Excel 2007 (www.microsoft.com) and the data charts in this study were generated and displayed using GraphPad Prism 8.3.0 (www.graphpad.com/).

## Data availability

RNA-seq data have been deposited in the NCBI SRA database under the accession number PRJNA596065. *ZmSOS1* gene sequences of D9H, Jing724 and B73 have been deposited in the NCBI GenBank database under the following accession numbers MN864203 (*ZmSOS1*^*D9H*^), MN864209 (*ZmSOS1*^*Jing724*^), and MN864201 (*ZmSOS1*^*B73*^). The sequence information of all vectors used in this study was shown in **Supplementary Dataset S1**.

## Acknowledgements

This work was supported by the Construction of Collaborative Innovation Center of Beijing Academy of Agricultural and Forestry Sciences (Grant No. KJCX201917), the National Natural Science Foundation of China (Grant No. 31860381), the Open Project from State Key Laboratory of Wheat and Maize Crop Science (Grant No. SKL2021KF07), the Public Institution Project of Maize Research Center (Grant No. YMZX202001), the Beijing Scholars Program (Grant No. BSP041). The authors would like to thank Prof. Yi Wang (China Agricultural University) for kindly providing yeast strain AXT3K and helping with the NMT experiment, Prof. Jiankang Zhu (Southern University of Science and Technology) for kindly providing *Arabidopsis sos1-1* mutant, and Profs. Jianbing Yan (Huazhong Agricultural University) and Xiaohong Yang (China Agricultural University) for kindly providing the DNAs of teosintes and maize landraces. This work was dedicated to the 25^th^ anniversary of Maize Research Center, Beijing Academy of Agriculture and Forestry Sciences.

## Author contributions

ZY (Yanxin Zhao), SW, WR and ZJ designed the experiment, conceived the project, and supervised the study; LM and ZY (Yanxin Zhao) conducted all the data analysis and wrote the manuscript; LJ, XJ, SX, DM and LB performed the field plant phenotyping; LM, ZY (Yunxia Zhang), ZP, LX and XY performed the gene mapping; CM, DD and LY performed *Arabidopsis* and maize transformation; ZY (Yanxin Zhao), ZR, WF and WY performed MAS breeding. All the authors read and approved the manuscript.

## Supporting Information

**Figure S1. High-salinity soils inhibit maize growth and yield**.

**Figure S2. RT-PCR analysis of the *qST1* candidate genes**.

**Figure S3. Sequence alignment of maize, *Arabidopsis* and rice SOS1 proteins**.

**Figure S4. Aberrant transcripts of *ZmSOS1* gene produced in *zmsos1-1* and *zmsos1-2* mutants**.

**Figure S5. Phenotypes of B73, *zmsos1-1*, and *zmsos1-2* plants under salt stress in the field**.

**Figure S6. The genotypes of *ZmSOS1* gene in diverse maize materials and teosintes**.

**Figure S7. The phenotype of X1132x-derived lines under salt stress**.

**Figure S8. A molecular marker-assisted selection strategy for improvement of maize salt tolerance with D9H as an example**.

**Figure S9. Growth inhibition of CIMBL50 and CIMBL84 seedlings under salt stress**.

**Figure S10. Association analysis of *ZmSOS1* gene and maize salt tolerance**.

**Table S1. The hybrid varieties certificated in China with x1132x-derived lines as parental plants (2009-2018)**.

**Table S2. Performance of the Jing724 and D9H F**_**2**_ **population seedlings under salt stress**.

**Table S3. The SNPs on chromosome 1 significantly associated with the salt tolerance of maize**.

**Table S4. KASP primers used for fine mapping of *qST1* in this study**.

**Table S5. Recombinant events identified within the salt-sensitive plants of Jing724 and D9H F**_**2**_ **population for fine mapping *qST1***.

**Table S6. X1132x-derived lines and Huangzaosi-improved lines used to screen the 4-bp InDel alleles**.

**Table S7. The current hybrid varieties certificated and cultivated in China used in this study**.

**Table S8. Maize landraces, natural population and teosintes used for genotyping *ZmSOS1* gene in this study**

**Table S9. List of the 55 SNPs used for association analysis of *ZmSOS1* gene**.

**Table S10. Phenotypes of shoot Na**^**+**^ **content and root length of GWAS lines under salt stress used for association analysis of *ZmSOS1* gene**

**Table S11. The PCR primers used in this study**.

**Table S12. Construction of yeast expression vectors for *ZmSOS1, AtSOS2*, and *AtSOS3* genes**

**Dataset S1. The detailed information of all vectors used in this study**.

## References

Abdel-Latif A, Osman G (2017) Comparison of three genomic DNA extraction methods to obtain high DNA quality from maize. Plant Methods 13: 1. doi: 10.1186/s13007-016-0152-4.

Apse MP, Aharon GS, Snedden WA, Blumwald E (1999) Salt tolerance conferred by overexpression of a vacuolar Na^+^ antiport in Arabidopsis. Science 285(5431): 1256–1258. doi: 10.1126/science.285.5431.1256.

Ariyarathna HA, Oldach KH, Francki MG (2016) A comparative gene analysis with rice identified orthologous group II HKT genes and their association with Na^+^ concentration in bread wheat. BMC Plant Biology 16:21. doi: 10.1186/s12870-016-0714-7.

Ashraf M, Foolad MR (2013) Crop breeding for salt tolerance in the era of molecular markers and marker-assisted selection. Plant Breeding 132(1): 10–20. doi: 10.1111/pbr.12000.

Asíns MJ (2002) Present and future of quantitative trait locus analysis in plant breeding. Plant Breeding 121(4): 281–291. doi: 10.1046/j.1439-0523.2002.730285.x

Asins MJ, Villalta I, Aly MM, Olías R, Alvarez DE Morales P, Huertas R, Li J, Jaime-Pérez N, Haro R, Raga V, Carbonell EA, Belver A (2013) Two closely linked tomato HKT coding genes are positional candidates for the major tomato QTL involved in Na^+^ homeostasis. Plant, Cell and Environment 36(6): 1171–1191. doi: 10.1111/pce.12051.

Barajas-Lopez JD, Moreno JR, Gamez-Arjona FM, Pardo JM, Punkkinen M, Zhu JK, Quintero FJ, Fujii H (2018) Upstream kinases of plant SnRKs are involved in salt stress tolerance. Plant Journal 93(1): 107–118. doi: 10.1111/tpj.13761.

Batistic O, Sorek N, Schültke S, Yalovsky S, Kudla J (2008) Dual fatty acyl modification determines the localization and plasma membrane targeting of CBL/CIPK Ca^+^ signaling complexes in Arabidopsis. Plant Cell 20(5): 1346–1362. doi: 10.1105/tpc.108.058123.

Byrt CS, Platten JD, Spielmeyer W, James RA, Lagudah ES, Dennis ES, Tester M, Munns R (2007) HKT1;5-like cation transporters linked to Na^+^ exclusion loci in wheat, Nax2 and Kna1. Plant Physiology 143(4): 1918–28. doi: 10.1104/pp.106.093476.

Cao YB, Zhang M, Liang XY, Li FR, Shi YL, Yang XH, Jiang CF (2020) Natural variation of an EF-hand Ca^2+^-binding-protein coding gene confers saline-alkaline tolerance in maize. Nature Communications 11(1):186. doi: 10.1038/s41467-019-14027-y.

Chinnusamy V, Jagendorf A, Zhu JK (2005) Understanding and improving salt tolerance in plants. Crop Science 45(2): 437–448. doi: 10.2135/cropsci2005.0437.

Davenport RJ, Muñoz-Mayor A, Jha D, Essah PA, Rus A, Tester M (2007) The Na^+^ transporter AtHKT1;1 controls retrieval of Na^+^ from the xylem in Arabidopsis. Plant, Cell and Environment 30(4): 497–507. doi: 10.1111/j.1365-3040.2007.01637.x.

Elble R (1992) A simple and efficient procedure for transformation of yeasts. Biotechniques 13(1): 18–20.

Fan YF, Yin XC, Xie Q, Xia YQ, Wang ZY, Song J, Zhou Y, Jiang XY (2019) Co-expression of SpSOS1 and SpAHA1 in transgenic Arabidopsis plants improves salinity tolerance. BMC Plant Biology 19(1): 74. doi: 10.1186/s12870-019-1680-7.

Fu JJ, Cheng YB, Linghu JJ, Yang XH, Kang L, Zhang ZX, Zhang J, He C, Du XM, Peng ZY, Wang B, Zhai LH, Dai CM, Xu JB, Wang WD, Li XR, Zheng J, Chen L, Luo LH, Liu JJ, Qian XJ, Yan JB, Wang J, Wang GY (2013) RNA sequencing reveals the complex regulatory network in the maize kernel. Nature Communications 4: 2832. doi: 10.1038/ncomms3832.

Glaubitz JC, Casstevens TM, Lu F, Harriman J, Elshire RJ, Sun Q, Buckler ES (2014) TASSEL-GBS: a high capacity genotyping by sequencing analysis pipeline. PLoS One 9(2): e90346. doi: 10.1371/journal.pone.0090346.

Hamamoto S, Horie T, Hauser F, Deinlein U, Schroeder JI, Uozumi N (2015) HKT transporters mediate salt stress resistance in plants: from structure and function to the field. Current Opinion in Biotechnology 32: 113–120. doi: 10.1016/j.copbio.2014.11.025.

Hasegawa PM (2013) Sodium (Na^+^) homeostasis and salt tolerance of plants. Environmental and Experimental Botany 92: 19–31. doi: 10.1016/j.envexpbot.2013.03.001.

Horie T, Hauser F, Schroeder JI (2009) HKT transporter-mediated salinity resistance mechanisms in Arabidopsis and monocot crop plants. Trends in Plant Science 14(12): 660–668. doi: 10.1016/j.tplants.2009.08.009.

Huang SB, Spielmeyer W, Lagudah ES, James RA, Platten JD, Dennis ES, Munns R (2006) A sodium transporter (HKT7) is a candidate for Nax1, a gene for salt tolerance in durum wheat. Plant Physiology 142(4): 1718–1727. doi: 10.1104/pp.106.088864.

Hufford MB, Seetharam AS, Woodhouse MR, Chougule KM, Ou SJ, Liu JN, Ricci WA, Guo TT, Olson A, Qiu YJ, Coletta RD, Tittes S, Hudson AI, Marand AP, Wei S, Lu ZY, Wang B, Tello-Ruiz MK, Piri RD, Wang N, Kim DW, Zeng YB, O’Connor CH, Li XR, Gilbert AM, Baggs E, Krasileva KV, Portwood JL, Cannon EKS, Andorf CM, Manchanda N, Snodgrass SJ, Hufnagel DE, Jiang QH, Pedersen S, Syring ML, Kudrna DA, Llaca V, Fengler K, Schmitz RJ, Ross-Ibarra J, Yu JM, Gent JI, Hirsch CN, Ware D, Dawe RK (2021) De novo assembly, annotation, and comparative analysis of 26 diverse maize genomes. Science 373(6555): 655–662. doi: 10.1126/science.abg5289.

Ismail A, Takeda S, Nick P (2014) Life and death under salt stress: same players, different timing? Journal of Experimental Botany 65(12): 2963–2979. doi: 10.1093/jxb/eru159.

Ismail AM, Horie T (2017) Genomics, physiology, and molecular breeding approaches for improving salt tolerance. Annual Review of Plant Biology 68:405–434. doi: 10.1146/annurev-arplant-042916-040936.

James RA, Davenport RJ, Munns R (2006) Physiological characterization of two genes for Na^+^ exclusion in durum wheat, Nax1 and Nax2. Plant Physiology 142(4): 1537–1547. doi: 10.1104/pp.106.086538.

Ji HT, Pardo JM, Batelli G, Van Oosten MJ, Bressan RA, Li X (2013) The Salt Overly Sensitive (SOS) pathway: established and emerging roles. Molecular Plant 6(2): 275–286. doi: 10.1093/mp/sst017.

Jiang ZL, Song GS, Shan XH, Wei ZY, Liu YZ, Jiang C, Jiang Y, Jin FX, Li YD (2018) Association analysis and identification of ZmHKT1;5 variation with salt-stress tolerance. Frontiers in Plant Science 9: 1485. doi: 10.3389/fpls.2018.01485.

Jiao YP, Peluso P, Shi JH, Liang T, Stitzer MC, Wang B, Campbell MS, Stein JC, Wei XH, Chin CS, Guill K, Regulski M, Kumari S, Olson A, Gent J, Schneider KL, Wolfgruber TK, May MR,, Springer NM, Antoniou E, McCombie WR, Presting GG, McMullen M, Ross-Ibarra J, Dawe RK, Hastie A, Rank DR, Ware D (2017) Improved maize reference genome with single-molecule technologies. Nature 546(7659): 524–527. doi: 10.1038/nature22971.

Kim WY, Ali Z, Park HJ, Park SJ, Cha JY, Perez-Hormaeche J, Quintero FJ, Shin G, Kim MR, Qiang Z, Ning L, Park HC, Lee SY, Bressan RA, Pardo JM, Bohnert HJ, Yun DJ (2013) Release of SOS2 kinase from sequestration with GIGANTEA determines salt tolerance in Arabidopsis. Nature Communications 4: 1352. doi: 10.1038/ncomms2357.

Kong MS, Luo MJ, Li JN, Feng Z, Zhang YX, Song W, Zhang RY, Wang RH, Wang YD, Zhao JR, Tao YS, Zhao YX (2021) Genome-wide identification, characterization, and expression analysis of the monovalent cation-proton antiporter superfamily in maize, and functional analysis of its role in salt tolerance. Genomics 113(4): 1940–1951. doi: 10.1016/j.ygeno.2021.04.032.

Kronzucker HJ, Britto DT (2011) Sodium transport in plants: a critical review. New Phytologist 189(1): 54–81. doi: 10.1111/j.1469-8137.2010.03540.x.

Li JG, Pu LJ, Han MF, Zhu M, Zhang RS, Xiang YZ (2014) Soil salinization research in China:advances and prospects. Journal of Geographical Sciences 24(5): 943–960. doi: 10.1007/s11442-014-1130-2.

Li PC, Pan T, Wang HM, Wei J, Chen MJ, Hu XH, Zhao Y, Yang XY, Yin SY, Xu Y, Fang HM, Liu J, Xu CW, Yang ZF (2019) Natural variation of ZmHKT1 affects root morphology in maize at the seedling stage. Planta 249(3): 879–889. doi: 10.1007/s00425-018-3043-2.

Lindsay MP, Lagudah ES, Hare RA, Munns R (2004) A locus for sodium exclusion (Nax1), a trait for salt tolerance, mapped in durum wheat. Functional Plant Biology 31(11): 1105–1114. doi: 10.1071/FP04111.

Liu SZ, Yeh CT, Tang HM, Nettleton D, Schnable PS (2012) Gene mapping via bulked segregant RNA-Seq (BSR-Seq). PLoS One 7(5): e36406. doi: 10.1371/journal.pone.0036406.

Livak KJ, Schmittgen TD (2001) Analysis of relative gene expression data using real-time quantitative PCR and the 2^-ΔΔCT^. Methods 25(4): 402–408. doi: 10.1006/meth.2001.1262.

Luo MJ, Lu BS, Shi YX, Zhao YX, Wei ZY, Zhang CY, Wang YD, Liu H, Shi YM, Yang JX, Song W, Lu XD, Fan YL, Xu L, Wang RH, Zhao JR (2022) A newly characterized allele of ZmR1 increases anthocyanin content in whole maize plant and the regulation mechanism of different ZmR1 alleles. Theoretical and Applied Genetics. doi: 10.1007/s00122-022-04166-0. [In press]

Luo MJ, Zhang YX, Chen K, Kong MS, Song W, Lu BS, Shi YX, Zhao YX, Zhao JR (2019a) Mapping of quantitative trait loci for seedling salt tolerance in maize. Molecular Breeding 39: 64. doi: 10.1007/s11032-019-0974-7.

Luo MJ, Zhang YX, Li JN, Zhang PP, Chen K, Song W, Wang XQ, Yang JX, Lu XD, Lu BS, Zhao YX, Zhao JR (2021) Molecular dissection of maize seedling salt tolerance using a genome-wide association analysis method. Plant Biotechnology Journal 19(10): 1937–1951. doi: 10.1111/pbi.13607.

Luo MJ, Zhao YX, Song W, Zhang RY, Su AG, Li CH, Wang XP, Xing JF, Shi Z, Zhao JR (2018a) Effect of saline stress on the physiology and growth of maize hybrids and their related inbred lines. Maydica 62(2): 1–8.

Luo MJ, Zhao YX, Wang YD, Shi Z, Zhang PP, Zhang YX, Song W, Zhao JR (2018b) Comparative proteomics of contrasting maize genotypes provides insights into salt-stress tolerance mechanisms. Journal of Proteome Research 17(1): 141–153. doi: 10.1021/acs.jproteome.7b00455.

Luo MJ, Zhao YX, Zhang RY, Xing JF, Duan MX, Li JN, Wang NS, Wang WG, Zhang SS, Chen ZH, Zhang HS, Shi Z, Song W, Zhao JR (2017) Mapping of a major QTL for salt tolerance of mature field-grown maize plants based on SNP markers. BMC Plant Biology 17(1): 140. doi: 10.1186/s12870-017-1090-7.

Luo X, Wang BC, Gao S, Zhang F, Terzaghi W, Dai M (2019b) Genome-wide association study dissects the genetic bases of salt tolerance in maize seedlings. Journal of Integrative Plant Biology 61(6): 658–674. doi: 10.1111/jipb.12797.

Ma L, Ye JM, Yang YQ, Lin HX, Yue LL, Luo J, Long Y, Fu HQ, Liu XN, Zhang YL, Wang Y, Chen LY, Kudla J, Wang YJ, Han SC, Song CP, Guo Y (2019) The SOS2-SCaBP8 complex generates and fine-tunes an AtANN4-dependent calcium signature under salt stress. Developmental Cell 48(5): 697-709.e5. doi: 10.1016/j.devcel.2019.02.010.

Maathuis FJM, Amtmann A (1999) K^+^ nutrition and Na^+^ toxicity: the basis of cellular K^+^ ratios. Annals of Botany 84(2): 123-133. doi.org/10.1006/anbo.1999.0912.

Martínez-Atienza J, Jiang XY, Garciadeblas B, Mendoza I, Zhu JK, Pardo JM, Quintero FJ (2007) Conservation of the salt overly sensitive pathway in rice. Plant Physiology 143(2): 1001–1012. doi: 10.1104/pp.106.092635.

McKenna A, Hanna M, Banks E, Sivachenko A, Cibulskis K, Kernytsky A, Garimella K, Altshuler D, Gabriel S, Daly M, DePristo MA (2010) The Genome Analysis Toolkit: a MapReduce framework for analyzing next-generation DNA sequencing data. Genome Research 20(9): 1297–1303. doi: 10.1101/gr.107524.110.

Møller IS, Gilliham M, Jha D, Mayo GM, Roy SJ, Coates JC, Haseloff J, Tester M (2009) Shoot Na^+^ exclusion and increased salinity tolerance engineered by cell type-specific alteration of Na^+^ transport in Arabidopsis. Plant Cell 21(7): 2163–2178. doi: 10.1105/tpc.108.064568.

Munns R, James RA, Läuchli A (2006) Approaches to increasing the salt tolerance of wheat and other cereals. Journal of Experimental Botany 57(5): 1025–1043. doi: 10.1093/jxb/erj100.

Munns R, James RA, Xu B, Athman A, Conn SJ, Jordans C, Byrt CS, Hare RA, Tyerman SD, Tester M, Plett D, Gilliham M (2012) Wheat grain yield on saline soils is improved by an ancestral Na^+^ transporter gene. Nature Biotechnology 30(4):360–364. doi: 10.1038/nbt.2120.

Munns R, Tester M (2008) Mechanisms of salinity tolerance. Annual Review of Plant Biology 59: 651–681. doi: 10.1146/annurev.arplant.59.032607.092911.

Ohta M, Guo Y, Halfter U, Zhu JK (2003) A novel domain in the protein kinase SOS2 mediates interaction with the protein phosphatase 2C ABI2. Proceedings of the National Academy of Sciences of the United States of America 100(20):11771–11776. doi: 10.1073/pnas.2034853100.

Ouhibi C, Attia H, Rebah F, Msilini N, Chebbi M, Aarrouf J, Urban L, Lachaal M (2014) Salt stress mitigation by seed priming with UV-C in lettuce plants: growth, antioxidant activity and phenolic compounds. Plant Physiology and Biochemistry 83:126–133. doi: 10.1016/j.plaphy.2014.07.019.

Pardo JM, Cubero B, Leidi EO, Quintero FJ (2006) Alkali cation exchangers: roles in cellular homeostasis and stress tolerance. Journal of Experimental Botany 57(5): 1181–1199. doi: 10.1093/jxb/erj114.

Qiu QS, Yan Guo, Dietrich MA, Schumaker KS, Zhu JK (2002) Regulation of SOS1, a plasma membrane Na+/H+ exchanger in Arabidopsis thaliana, by SOS2 and SOS3. Proceedings of the National Academy of Sciences of the United States of America 99(12): 8436–8441. doi: 10.1073/pnas.122224699.

Quan RD, Lin HX, Mendoza I, Zhang YG, Cao WH, Yang YQ, Shang M, Chen SY, Pardo JM, Guo Y (2007) SCABP8/CBL10, a putative calcium sensor, interacts with the protein kinase SOS2 to protect Arabidopsis shoots from salt stress. Plant Cell 19(4):1415–1431. doi: 10.1105/tpc.106.042291.

Quintero FJ, Martinez-Atienza J, Villalta I, Jiang XY, Kim WY, Ali Z, Fujii H, Mendoza I, Yun DJ, Zhu JK, Pardo JM (2011) Activation of the plasma membrane Na/H antiporter Salt-Overly-Sensitive 1 (SOS1) by phosphorylation of an auto-inhibitory C-terminal domain. Proceedings of the National Academy of Sciences of the United States of America 108(6): 2611–2616. doi: 10.1073/pnas.1018921108.

Quintero FJ, Ohta M, Shi HZ, J., Pardo JM (2002) Reconstitution in yeast of the Arabidopsis SOS signaling pathway for Na^+^ homeostasis. Proceedings of the National Academy of Sciences of the United States of America 99(13): 9061–9066. doi: 10.1073/pnas.132092099.

Ren ZH, Gao JP, Li LG, Cai XL, Huang W, Chao DY, Zhu MZ, Wang ZY, Luan S, Lin HX (2005) A rice quantitative trait locus for salt tolerance encodes a sodium transporter. Nature Genetics 37(10): 1141–1146. doi: 10.1038/ng1643.

Rodríguez-Navarro A, Ramos J (1984) Dual system for potassium transport in Saccharomyces cerevisiae. Journal of Bacteriology 159(3): 940–945. doi: 10.1128/jb.159.3.940-945.1984.

Roy SJ, Negrão S, Tester M (2014) Salt resistant crop plants. Current Opinion in Biotechnology 26: 115–124. doi: 10.1016/j.copbio.2013.12.004.

Rus A, Baxter I, Muthukumar B, Gustin J, Lahner B, Yakubova E, Salt DE (2006) Natural variants of AtHKT1 enhance Na^+^ accumulation in two wild populations of Arabidopsis. PLoS Genetics 2(12): e210. doi: 10.1371/journal.pgen.0020210.

Rus A, Lee BH, Muñoz-Mayor A, Sharkhuu A, Miura K, Zhu JK, Bressan RA, Hasegawa PM (2004) AtHKT1 facilitates Na^+^ homeostasis and K^+^ nutrition in planta. Plant Physiology 136(1): 2500–2511. doi: 10.1104/pp.104.042234.

Salvi S, Tuberosa R (2005) To clone or not to clone plant QTLs: present and future challenges. Trends in Plant Science 10(6): 297–304. doi: 10.1016/j.tplants.2005.04.008.

Shi HZ, Ishitani M, Kim C, Zhu JK (2000) The Arabidopsis thaliana salt tolerance gene SOS1 encodes a putative Na^+^ antiporter. Proceedings of the National Academy of Sciences of the United States of America 97(12): 6896–6901. doi: 10.1073/pnas.120170197.

Shi HZ, Lee BH, Wu SJ, Zhu JK (2003) Overexpression of a plasma membrane Na^+^ antiporter gene improves salt tolerance in Arabidopsis thaliana. Nature Biotechnology 21(1): 81–85. doi: 10.1038/nbt766.

Shi HZ, Quintero FJ, Pardo JM, Zhu JK (2002) The putative plasma membrane Na^+^ antiporter SOS1 controls long-distance Na^+^ transport in plants. Plant Cell 14(2): 465–477. doi: 10.1105/tpc.010371.

Sun W, Xiang XL, Zhai LH, Zhang D, Cao Z, Liu L, Zhang ZX (2018) AGO18b negatively regulates determinacy of spikelet meristems on the tassel central spike in maize. Journal of Integrative Plant Biology 60(1): 65–78. doi: 10.1111/jipb.12596.

Sunarpi H, Horie T, Motoda J, Kubo M, Yang H, Yoda K, Horie R, Chan WY, Leung HY, Hattori K, Konomi M, Osumi M, Yamagami M, Schroeder JI, Uozumi N (2005) Enhanced salt tolerance mediated by AtHKT1 transporter-induced Na unloading from xylem vessels to xylem parenchyma cells. Plant Journal 44(6): 928–938. doi: 10.1111/j.1365-313X.2005.02595.x.

Tan TH, Cai JQ, Zhan EB, Yang YQ, Zhao JF, Guo Y, Zhou HP (2016) Stability and localization of 14-3-3 proteins are involved in salt tolerance in Arabidopsis. Plant Molecular Biology 92(3): 391–400. doi: 10.1007/s11103-016-0520-5.

Tang B, Luo MJ, Zhang YX, Guo HL, Li JN, Song W, Zhang RY, Feng Z, Kong MS, Li H, Cao ZY, Lu XD, Li DL, Zhang JH, Wang RH, Wang YD, Chen ZH, Zhao YX, Zhao JR (2021) Natural variations in the P-type ATPase heavy metal transporter gene ZmHMA3 control cadmium accumulation in maize grains. Journal of Experimental Botany 72(18): 6230–6246. doi: 10.1093/jxb/erab254.

Wang FG, Tian HL, Zhao JR, Yi HM, Wang L, Song W (2011) Development and characterization of a core set of SSR markers for fingerprinting analysis of Chinese maize varieties. Maydica 56(1): 7–18.

Wang YD, Zhang HS, Duan MX, Zhang XY, Zhang CY, Chen CY, Zhao JR (2015) Study on the elite inbred lines from exotic hybrid X1132x. China Seed Industry (2): 41–44. doi: 10.19462/j.cnki.1671-895x.2015.02.017 [in Chinese]

Wu TD, Reeder J, Lawrence M, Becker G, Brauer MJ (2016) GMAP and GSNAP for genomic sequence alignment: enhancements to speed, accuracy, and functionality. Methods in Molecular Biology 1418: 283–334. doi: 10.1007/978-1-4939-3578-9_15.

Xu, HG (2004) Halophyte and ecological management of salinization in China. Chinese Agricultural Science and Technology Press: Beijing, China.

Yang Q, Chen ZZ, Zhou XF, Yin HB, Li X, Xin XF, Hong XH, Zhu JK, Gong ZZ (2009) Overexpression of SOS (Salt Overly Sensitive) genes increases salt tolerance in transgenic Arabidopsis. Molecular Plant 2(1): 22–31. doi: 10.1093/mp/ssn058.

Yang YQ, Guo Y (2018a) Elucidating the molecular mechanisms mediating plant salt-stress responses. New Phytologist 217(2): 523–539. doi: 10.1111/nph.14920.

Yang YQ, Qin YX, Xie CG, Zhao FY, Zhao JF, Liu DF, Chen SY, Fuglsang AT, Palmgren MG, Schumaker KS, Deng XW, Guo Y (2010) The Arabidopsis chaperone J3 regulates the plasma membrane H^+^-ATPase through interaction with the PKS5 kinase. Plant Cell 22(4): 1313–1332. doi: 10.1105/tpc.109.069609.

Yang YQ, Yan Guo (2018b) Unraveling salt stress signaling in plants. Journal of Integrative Plant Biology 60(9): 796–804. doi: 10.1111/jipb.12689.

Yu LJ, Nie JN, Cao CY, Jin YK, Yan M, Wang FZ, Liu J, Xiao Y, Liang YH, Zhang WH (2010) Phosphatidic acid mediates salt stress response by regulation of MPK6 in Arabidopsis thaliana. New Phytologist 188(3): 762–773. doi: 10.1111/j.1469-8137.2010.03422.x.

Zhang DF, Wu SW, An XL, Xie K, Dong ZY, Zhou Y, Xu LW, Fang W, Liu SS, Liu SS, Zhu TT, Li JP, Rao LQ, Zhao JR, Wan XY (2017) Construction of a multicontrol sterility system for a maize male-sterile line and hybrid seed production based on the ZmMs7 gene encoding a PHD-finger transcription factor. Plant Biotechnology Journal 16(2): 459–471. doi: 10.1111/pbi.12786.

Zhang HX, Blumwald E (2001) Transgenic salt-tolerant tomato plants accumulate salt in foliage but not in fruit. Nature Biotechnology 19(8):765–768. doi: 10.1038/90824.

Zhang M, Cao YB, Wang ZP, Wang ZQ, Shi JP, Liang XY, Song WB, Chen QJ, Lai JS, Jiang CF (2018) A retrotransposon in an HKT1 family sodium transporter causes variation of leaf Na ^+^ exclusion and salt tolerance in maize. New Phytologist 217(3): 1161–1176. doi: 10.1111/nph.14882.

Zhang M, Liang XY, Wang LM, Cao YB, Song WB, Shi JP, Lai JS, Jiang CF (2019) A HAK family Na^+^ transporter confers natural variation of salt tolerance in maize. Nature Plants 5(12): 1297–1308. doi: 10.1038/s41477-019-0565-y.

Zhang XR, Henriques R, Lin SS, Niu QW, Chua NH (2006) Agrobacterium-mediated transformation of Arabidopsis thaliana using the floral dip method. Nature Protocols 1(2): 641–646. doi: 10.1038/nprot.2006.97.

Zhou Y, Yin XC, Duan RJ, Hao GP 4, Guo JC, Jiang XY (2015) SpAHA1 and SpSOS1 coordinate in transgenic yeast to improve salt tolerance. PLoS One 10(9): e0137447. doi: 10.1371/journal.pone.0137447.

Zhu JK (2003) Regulation of ion homeostasis under salt stress. Current Opinion in Plant Biology 6(5):441–445. doi: 10.1016/s1369-5266(03)00085-2.

hu JK (2016) Abiotic stress signaling and responses in plants. Cell 167(2): 313–324. doi: 10.1016/j.cell.2016.08.029.

