## Supplementary Figures S1-S10 for "A 4-bp natural deletion of maize Na^+^/H^+^ exchanger gene alters maize salt stress tolerance"

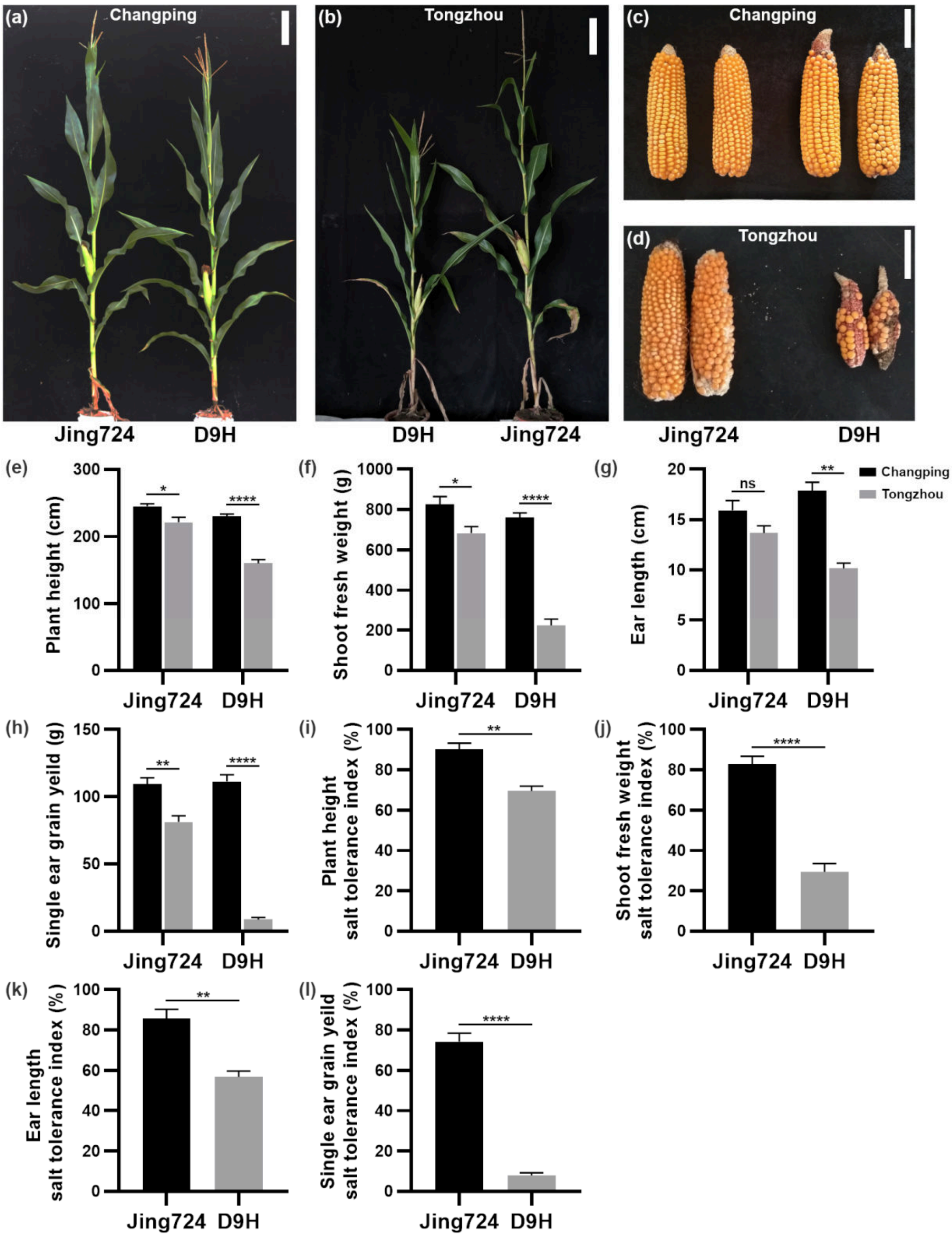


**Figure S1. High-salinity soils inhibit maize growth and yield.**

Jing724 and D9H were grown in Changping (CP, normal soil) and Tongzhou (TZ, high-salinity soil) with the sowing date of April 25, 2017. (a, b) Jing724 and D9H plant performance in CP (a) and TZ (b) 20 days after pollination (DAP). Bar = 20 cm. (c, d) Ears of Jing724 and D9H in CP (c) and TZ (d). Bar = 5cm. (e) 20-DAP plant height. (f) 20-DAP shoot fresh weight. (g) Ear length. (h) Single ear grain yield. The legends for charts (e-h) were shown on the right of figure (g). (i-l) Salt tolerance indexes of plant height (i), shoot fresh weight (j), ear length (k), and single ear grain yield (l) are calculated as the ratio of phenotypes in TZ to those in CP. Four replicates were performed for each data and five plants were used for each replicate. Data were analyzed using a two-tailed Student's *t*-test. Error bar = standard error. ns, not statistically significant. *, $P < 0.05$. **, $P < 0.01$. ***, $P < 0.001$. ****, $P < 0.0001$.

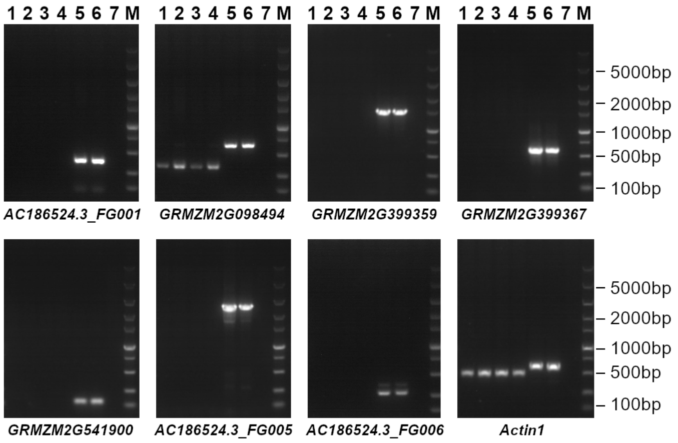


**Figure S2. RT-PCR analysis of the *qST1* candidate genes.**

RT-PCRs were performed to analyze the expression of seven candidate genes with cDNA of Jing724 and D9H seedlings as templates and gene-specific primers shown in **Supplementary Table S11**. *Actin1* (*GRMZM2G126010*) gene was the positive control for RT-PCRs. Lane 1, Jing724 seedling cDNA under normal condition; lane 2, Jing724 seedling cDNA under 100 mM NaCl treatment for 24 h; lane 3, D9H seedling cDNA under normal condition; lane 4, D9H seedling cDNA under 100 mM NaCl treatment for 24 h; lane 5, Jing724 seedling DNA; lane 6, D9H seedling DNA; lane 7, negative control without templates. M, DL5000 DNA marker (Vazyme Biotech., Nanjing, China).

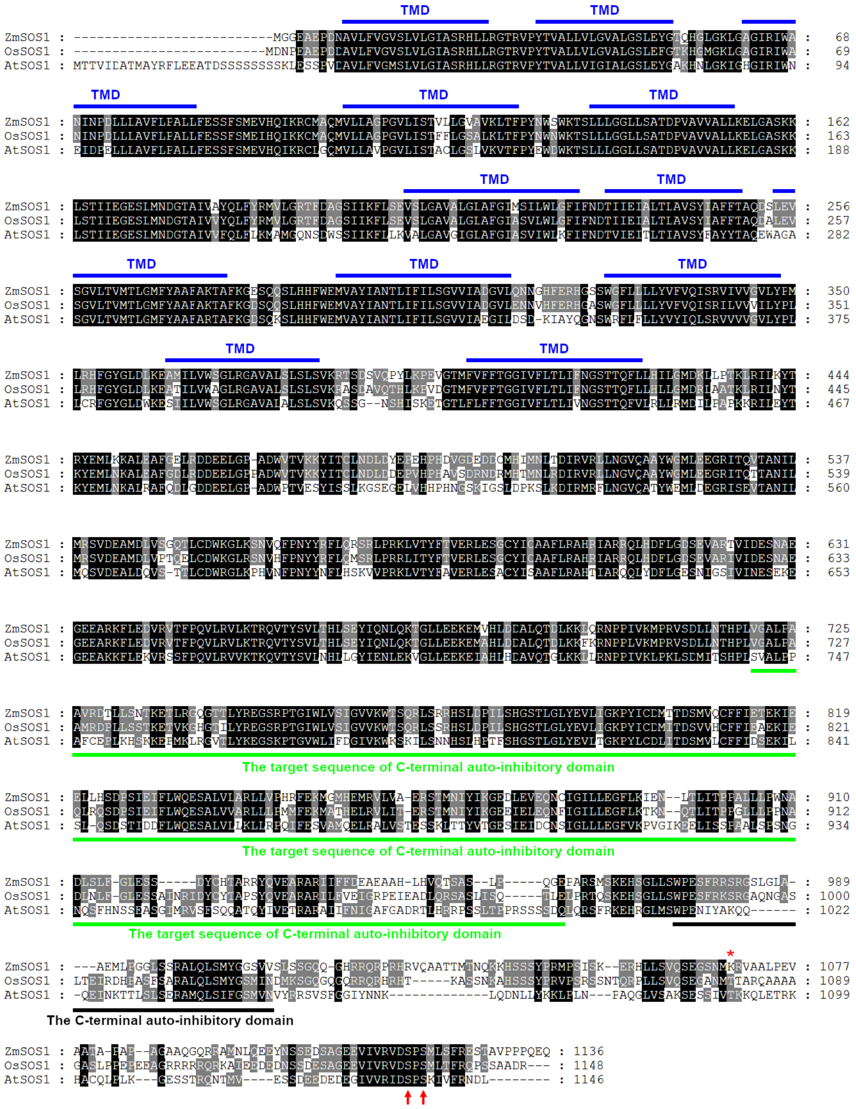


**Figure S3. Sequence alignment of maize, *Arabidopsis* and rice SOS1 proteins.**

The amino acid sequence alignment of ZmSOS1$^{Jing724}$, AtSOS1 (AT2G01980, GenBank accession number NP_178307) and OsSOS1 (GenBank accession number AAW33875) was performed with Clustal Omega (https://www.ebi.ac.uk/Tools/msa/clustalo/) and displayed with GenDoc (Nicholas *et al*., 1997). Transmembrane domains (TMDs) of ZmSOS1$^{Jing724}$ protein predicted by the TMHMM 2.0 online server (https://services.healthtech.dtu.dk/service.php?TMHMM-2.0) were overlined, and the auto-inhibitory domain and the target sequence of the auto-inhibitory domain of AtSOS1 were underlined (Quintero *et al*., 2011). The conserved two serine residues (S1136 and S1138 in AtSOS1) subjecting to phosphorylation were highlighted by red arrows (Quintero *et al*., 2011). The amino acid marked by a red asterisk and all amino acid residues following it were altered in ZmSOS$^{D9H}$ due to the 4-bp deletion.

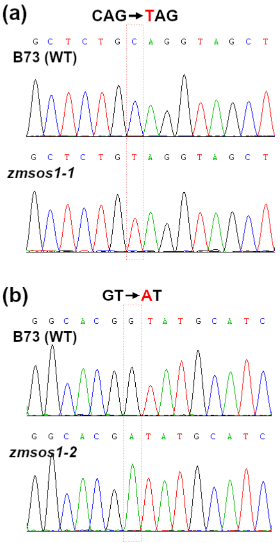


(c)

| Transcripts | Clones | Alternative splicing events |
|---|---|---|
| *zmsos1-2_T01* | 3 | 22-bp (+2,755 - +2,776 in exon 5) **exon skipping, and 11-bp** (+6,384 - +6,394 in intron 11) **intron retention** |
| *zmsos1-2_T02* | 2 | 49-bp (+5,935 - +5,983 in intron 9) **intron retention, and 11-bp** (+6,384 - +6,394 in intron 11) **intron retention** |
| *zmsos1-2_T03* | 2 | **exon 11** (+6,328 - +6,383) **skipping** |
| *zmsos1-2_T04* | 1 | **11-bp** (+6,384 - +6,394 in intron 11) **intron retention** |
| *zmsos1-2_T05* | 1 | **49-bp** (+5,935 - +5,983 in intron 9) **intron retention, 11-bp** (+6,384 - +6,394 in intron 11) **intron retention, and 4-bp** (+9,848 - +9,851 in exon 12) **exon skipping** |
| *zmsos1-2_T06* | 1 | **22-bp** (+2,755 - +2,776 in exon 5) **exon skipping, and exon 11** (+6,328 - +6,383) **skipping** |

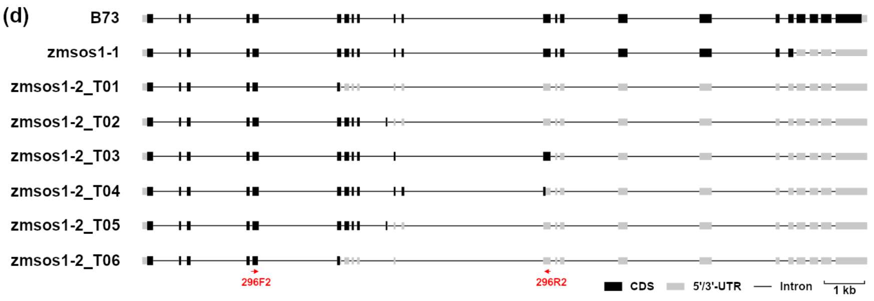


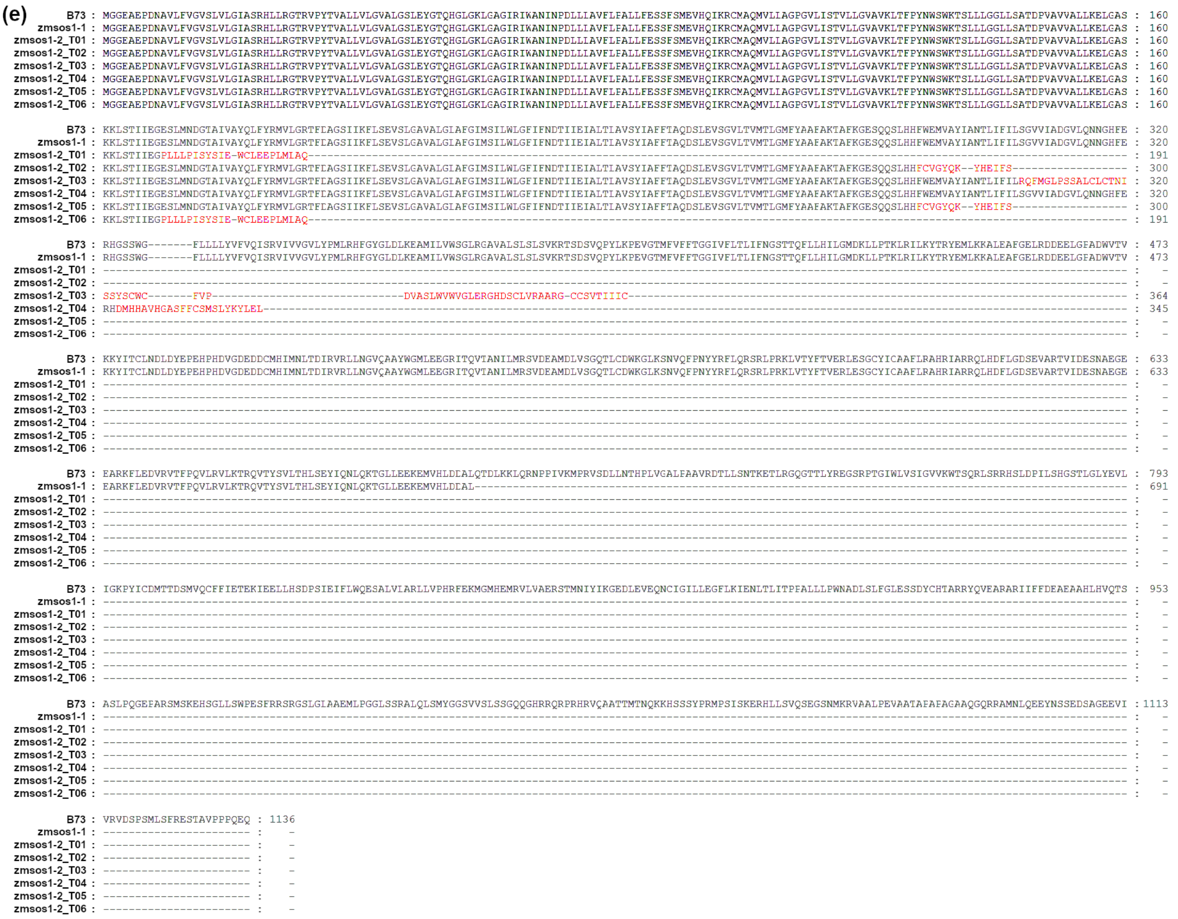


**Figure S4.** **Aberrant transcripts of *ZmSOS1* gene produced in *zmsos1-1* and *zmsos1-2* mutants.** (a, b) PCR sequencing validation of the point mutation sites in *zmsos1-1* (a) and *zmsos1-2* mutants (b). (c-e) Transcript and protein sequence changes of *ZmSOS1* in *zmsos1-1* and *zmsos1-2*. RT-PCR was performed with *zmsos1-2* seedling cDNA as template and the primer pair 296F2/296R2 (**Table S11**) to identify the aberrant splicing events caused by the G-to-A splice acceptor site mutation in *zmsos1-2*. (c) Six types of incorrectly spliced transcripts were identified from sequenced 10 cDNA clones. The nucleotide positions were based on the start codon positions (NATG, A = +1). (d) Gene structures of *ZmSOS1* genes in B73, *zmsos1-1* and *zmsos1-2*. The full-length cDNA sequences of *zmsos1-2* transcripts were speculated on the basis of the above RT-PCR results and were aligned to genomic DNA sequence to build the gene structures within GSDS 2.0 server (http://gsds.gao-lab.org/). (e) Alignment of ZmSOS1 protein sequences of B73, *zmsos1-1*, and *zmsos1-2*. The aberrant amino acid residues caused by the aberrant splicing events in *zmsos1-2* mutant were highlighted in red.

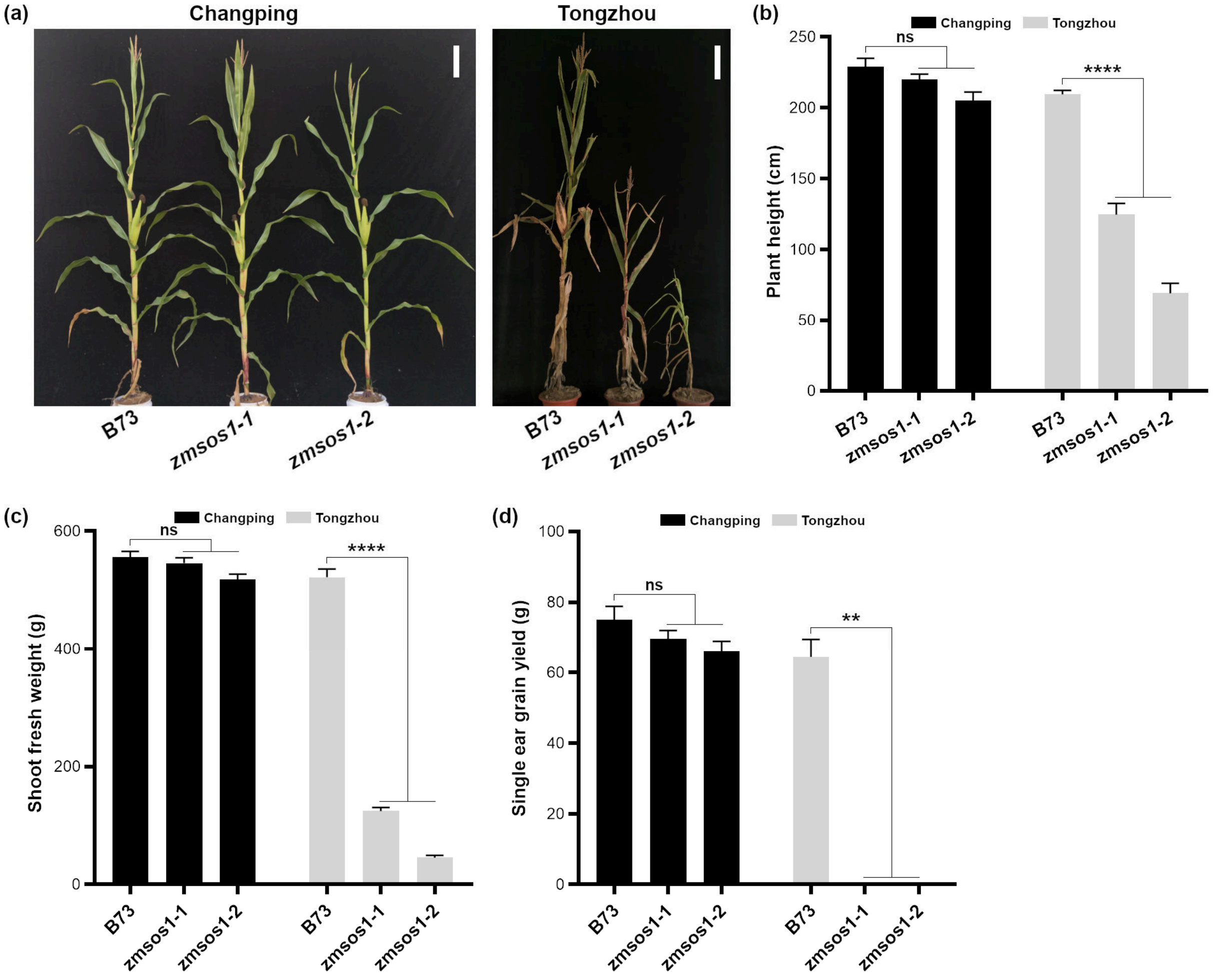


**Figure S5. Phenotypes of B73, *zmsos1-1*, and *zmsos1-2* plants under salt stress in the field.** EMS mutants *zmsos1-1*, *zmsos1-2*, and wild-type B73 were planted in Changping (CP, normal soil) and Tongzhou (TZ, high-salinity soil) on April 28, 2019. (a) Photos of B73, *zmsos1-1*, and *zmsos1-2* plants were taken at 20 days after pollination (DAP) in CP and at 45 DAP (before harvesting) in TZ. Bar = 20 cm. (b) Plant height. (c) Shoot fresh weight. (d) Single ear grain yield. No ear was produced in *zmsos1-1* and *zmsos1-2* plants grown in TZ. Four replicates were performed for each data and five plants were used for each replicate. Data were analyzed using a two-tailed Student's *t*-test. Error bar = standard error. ns, not statistically significant. **, $P < 0.01$. ****, $P < 0.0001$.

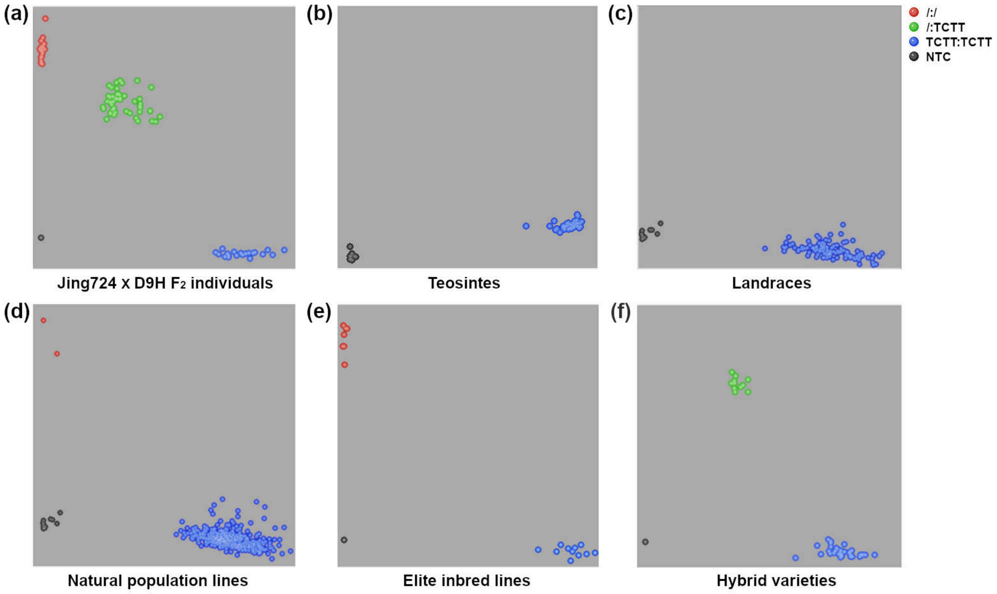


**Figure S6. The genotypes of *ZmSOS1* gene in diverse maize materials and teosintes.**

The 4-bp InDel KASP marker was used to screen the genotypes of *ZmSOS1* gene in 95 individuals of Jing724×D9H $F_2$ population (a), 52 teosintes (b), 133 landraces (c), 489 national population inbred lines (d), 14 x1132x-derived lines (e), 4 Huangzaosi-improved lines (e), and 45 hybrid varieties (f). Detailed information on the names, numbers, and genotypes of these materials were shown in **Supplementary Tables S2, S6, S7, and S8**. NTC, no template control.

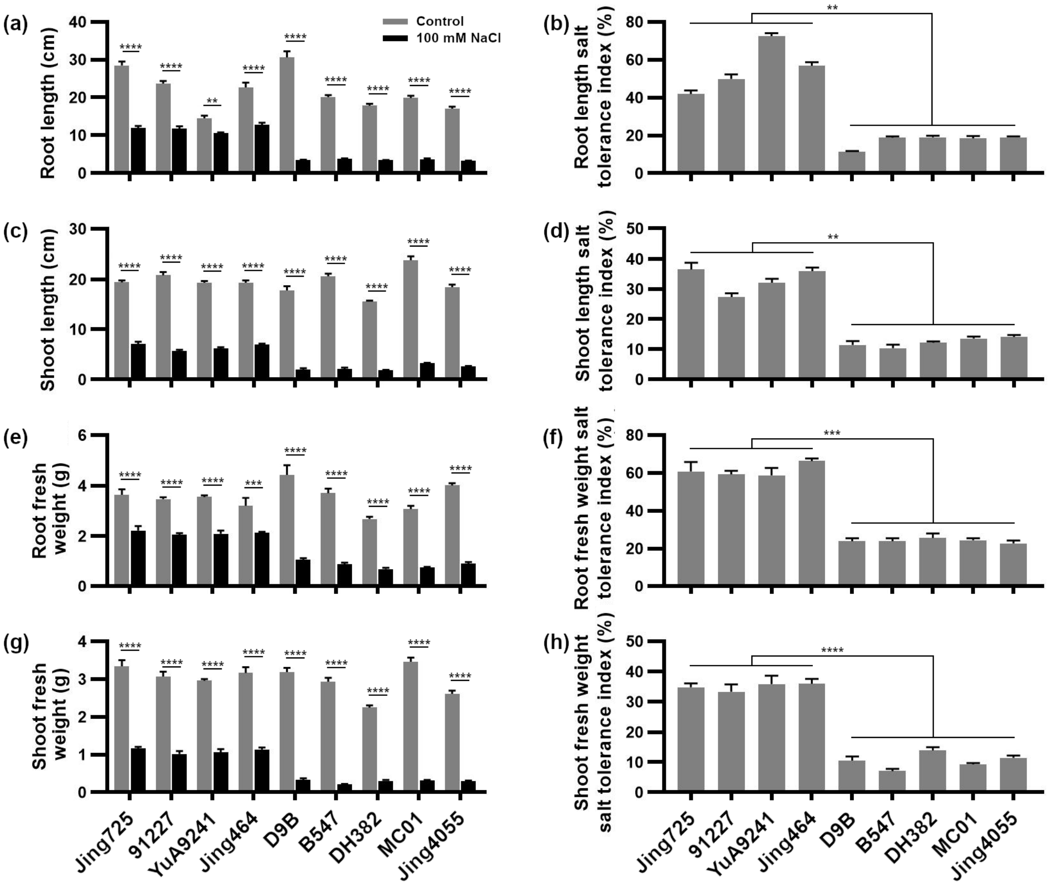


**Figure S7. The phenotype of X1132x-derived lines under salt stress.**

Salt tolerance performance of 9 X line seedlings was investigated after 10-day exposure to 100 mM NaCl treatment and the control was 10-day treatment with sterile water. Of them, 4 lines (Jing725, 91227, YuA9421 and Jing464) have the *ZmSOS1*$^{\text{Jing724}}$ allele (TCTT:TCTT) and 5 lines (D9B, B547, DH382, CM01 and Jing4055) contain the *ZmSOS1*$^{\text{D9H}}$ allele (/:/) (**Figure S6; Table S6**). (e, g) Each fresh weight value was measured with five seedlings. (b, d, f, h) The salt tolerance index was the ratio of phenotypes under salt stress conditions to those under control conditions. Four replicates were performed for each data and five plants were used for each replicate. Data were analyzed using a two-tailed Student's *t*-test. Error bar = standard error. **, $P < 0.01$. ***, $P < 0.001$. ****, $P < 0.0001$.

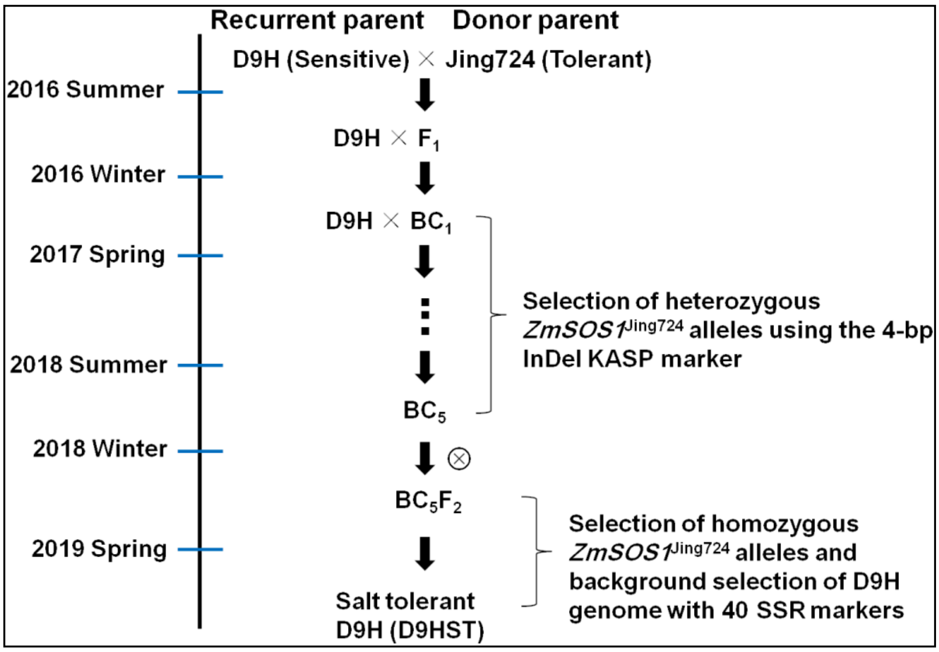


**Figure S8. A molecular marker-assisted selection strategy for improvement of maize salt tolerance with D9H as an example.**

The salt-tolerant *ZmSOS1*$^{Jing724}$ allele was transferred into the D9H genome to substitute the salt-sensitive *ZmSOS1*$^{D9H}$ allele using a backcrossing approach with D9H as the recurrent parent. From generations $BC_1$ to $BC_5F_2$, the 4-bp InDel KASP marker was used to screen and select the individuals carrying the *ZmSOS1*$^{Jing724}$ allele. Finally, the salt-tolerant D9HST individuals harboring homozygous *ZmSOS1*$^{Jing724}$ gene were subjected to background selection with 40 pairs of SSR marker primers (Wang *et al*., 2011).

(a)

| Name | Pedigree | Source |
|---|---|---|
| CIMBL50 | ([(DTPYC9-F74-1-1-1-1-B)-B X (DTPYC9-F65-2-2-1-1-B)-B]-B-3-2-B)-B0 | CIMMYT |
| CIMBL84 | (DTPYC9-F74-1-1-1-1-B)-B | CIMMYT |

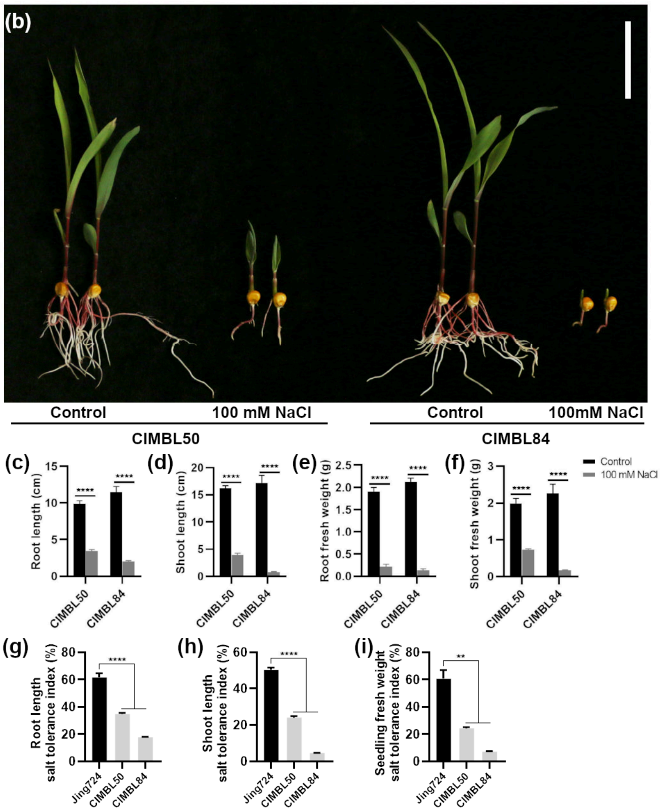


**Figure S9. Growth inhibition of CIMBL50 and CIMBL84 seedlings under salt stress.**

(a) Summary of CIMBL50 and CIMBL84. (b-g) Phenotypic comparison of CIMBL50 and CIMBL84 seedlings which had germinated and grown for 10 days under normal conditions and 100 mM NaCl treatment. (b) Picture of CIMBL50 and CIMBL84 seedlings under 100 mM NaCl treatment. Bar = 5 cm. (c) Root length. (d) Shoot length. (e) Root fresh weight of 5 seedlings. (f) Shoot fresh weight of 5 seedlings. The legends for charts (c-f) were shown on the right of figure (f). (g-i) Salt tolerance indexes of root length (g), shoot length (h), and one-seedling fresh weight (sum of root and shoot fresh weight) (i) were calculated by the ratio of phenotypes under salt stress conditions to those under normal conditions. Four replicates were performed for each data and five plants were used for each replicate. Data were analyzed using a two-tailed Student's *t*-test. Error bar = standard error. **, $P < 0.01$. ****, $P < 0.0001$.

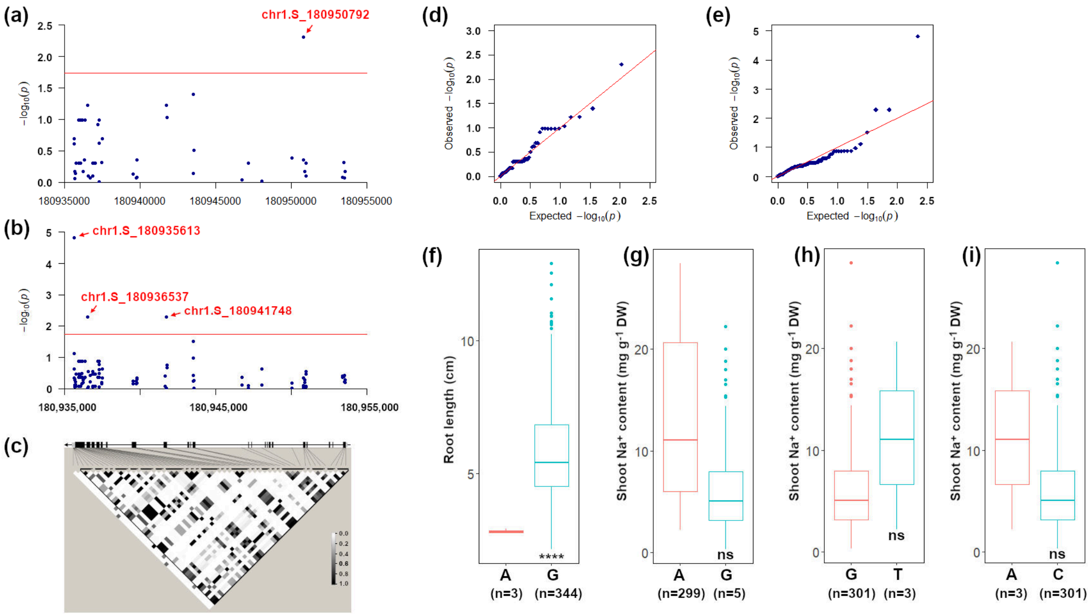


**Figure S10. Association analysis of *ZmSOS1* gene and maize salt tolerance.**

Association analysis of *ZmSOS1* gene was conducted with the phenotypic data of shoot $Na^+$ content and root length under salt stress (**Table S10**), and the genotypic data of the 55 SNPs in *ZmSOS1* gene (**Table S9**). (a) Manhattan plots calculated using root length data under 100 mM NaCl treatment obtained from Luo *et al*. (2021) (b) Manhattan plots calculated using shoot $Na^+$ content data under 100 mM NaCl treatment obtained from Cao *et al*. (2020) The cutoff value of 1.74 was indicated by the red line. The chromosome positions of the SNPs were based on the B73 RefGen_v2 genome. (c) The linkage disequilibrium structure between the 55 SNPs in *ZmSOS1*. (d, e) Quantile-Quantile plots for the association analysis of root length (d) and shoot $Na^+$ content (e). (f-i) Phenotypic differences of the two alleles of the significant SNPs identified above, including chr1.S_180950792 (f), chr1.S_180935613 (g), chr1.S_180936537 (h) and chr1.S_180941748 (i). DW, dry weight. Data were analyzed using a two-tailed Student's *t*-test. ns, not statistically significant. ****, $P < 0.0001$.

### Reference


**Nicholas KB, Nicholas HB, Deerfield DW (1997)** GeneDoc: Analysis and visualization of genetic variation. *EMBnet News* **4**:14-17.
