## Supplementary Tables S1-S12 for "A 4-bp natural deletion of maize Na^+^/H^+^ exchanger gene alters maize salt stress tolerance"

**Table S1. The hybrid varieties certificated in China with x1132x-derived lines as parental plants (2009-2018).**

| Variety Name | Maternal parent | Paternal parent | Approval Number | Bred and provided by |
|---|---|---|---|---|
| Denghai662 (登海662) | DH371 | DH382 | GUOSHENYU2009010 | Shandong Denghai Seeds Co., Ltd. |
| Denghai605 (登海605) | DH351 | DH382 | GUOSHENYU2010009 | Shandong Denghai Seeds Co., Ltd. |
| Jingke968 (京科968) | Jing724 | Jing92 | GUOSHENYU2011007; GUOSHENYU20180314 | Maize Research Center, Beijing Academy of Agriculture and Forestry Sciences |
| Jingnongke728 (京农科728) | MC01 | Jing2416 | GUOSHENYU2012003; GUOSHENYU20170007 | Maize Research Center, Beijing Academy of Agriculture and Forestry Sciences |
| Jingke665 (京科665) | Jing725 | Jing92 | GUOSHENYU2013003 | Maize Research Center, Beijing Academy of Agriculture and Forestry Sciences |
| MC220 | JingX220 | JingC632 | GUOSHENYU2013018 | Maize Research Center, Beijing Academy of Agriculture and Forestry Sciences |
| Denghai685 (登海685) | DH382 | DH357-14 | GUOSHENYU2015011 | Shandong Denghai Seeds Co., Ltd. |
| Yudan606 (豫单606) | YuA9241 | XinA3 | GUOSHENYU2015018 | Henan Agricultural University |
| Shandan609 (陕单609) | 91227 | Chang7-2 | GUOSHENYU2016001 | Northwest Agriculture and Forestry University |
| Denghai533 (登海533) | Denghai22 | DH382 | GUOSHENYU20176064 | Shandong Denghai Seeds Co., Ltd. |
| MC121 | Jing72464 | Jing2416 | GUOSHENYU20180070 | Maize Research Center, Beijing Academy of Agriculture and Forestry Sciences |
| JK9681 | Jing724 | Jing92H | GUOSHENYU20180231 | Maize Research Center, Beijing Academy of Agriculture and Forestry Sciences |
| NK718 | Jing464 | Jing2416 | GUOSHENYU20180261 | Maize Research Center, Beijing Academy of Agriculture and Forestry Sciences |
| Denghai695 (登海695) | DH382 | Denghai057 | GUOSHENYU20186120 | Shandong Denghai Seeds Co., Ltd. |
| Jingke528 (京科528) | 90110-2(D9H) | J2437(Jing2416) | JINGSHENYU2008008 | Maize Research Center, Beijing Academy of Agriculture and Forestry Sciences |
| Jingdan38 (京单38) | D9B | Jing2416 | JINGSHENYU2009005 | Maize Research Center, Beijing Academy of Agriculture and Forestry Sciences |
| MC4592 | Jing4055 | Jing92 | JINGSHENYU2014001 | Maize Research Center, Beijing Academy of Agriculture and Forestry Sciences |
| MC812 | JingB547 | Jing2416 | JINGSHENYU2015003 | Maize Research Center, Beijing Academy of Agriculture and Forestry Sciences |

Note: x1132x-derived lines were indicated in red and Huangzaosi-improved lines were denoted in green

**Table S2. Performance of the Jing724 and D9H $F_2$ population seedlings under salt stress.**

| Jing724×D9H $F_2$ individuels | Root length(cm) | Genotype by the 4-bp InDel KASP marker |
|---|---|---|
| 1 | 9.3 | /:TCTT |
| 2 | 3.4 | /:/ |
| 3 | 8.8 | /:TCTT |
| 4 | 9 | /:TCTT |
| 5 | 3.2 | /:/ |
| 6 | 11.1 | TCTT:TCTT |
| 7 | 7.4 | /:TCTT |
| 8 | 3.2 | /:/ |
| 9 | 7.2 | TCTT:TCTT |
| 10 | 10.2 | /:TCTT |
| 11 | 2.7 | /:/ |
| 12 | 11.8 | /:TCTT |
| 13 | 8.2 | /:TCTT |
| 14 | 11 | /:TCTT |
| 15 | 9.1 | TCTT:TCTT |
| 16 | 1.7 | /:/ |
| 17 | 7.1 | TCTT:TCTT |
| 18 | 9.8 | TCTT:TCTT |
| 19 | 9.7 | /:TCTT |
| 20 | 7.7 | /:TCTT |
| 21 | 11.4 | /:TCTT |
| 22 | 10.6 | /:TCTT |
| 23 | 2.5 | /:/ |
| 24 | 10.2 | /:TCTT |
| 25 | 10.5 | /:TCTT |
| 26 | 4.1 | /:/ |
| 27 | 10.7 | /:TCTT |
| 28 | 8.5 | /:TCTT |
| 29 | 11.6 | /:TCTT |
| 30 | 12.8 | /:TCTT |
| 31 | 7.9 | /:TCTT |
| 32 | 9 | /:TCTT |
| 33 | 8.7 | TCTT:TCTT |
| 34 | 9.4 | /:TCTT |
| 35 | 10.4 | /:TCTT |
| 36 | 7.6 | TCTT:TCTT |
| 37 | 9.2 | TCTT:TCTT |
| 38 | 9.5 | /:TCTT |
| 39 | 9.3 | /:TCTT |
| 40 | 9.2 | /:TCTT |
| 41 | 2.8 | /:/ |
| 42 | 7.7 | TCTT:TCTT |
| 43 | 8.9 | TCTT:TCTT |
| 44 | 11.8 | /:TCTT |
| 45 | 7.5 | TCTT:TCTT |
| 46 | 8.2 | TCTT:TCTT |
| 47 | 8.2 | TCTT:TCTT |
| 48 | 7.1 | /:TCTT |
| 49 | 7.5 | /:TCTT |
| 50 | 11.8 | /:TCTT |

| | | |
|---|---|---|
| 51 | 3.7 | /:/ |
| 52 | 10.6 | /:TCTT |
| 53 | 12.9 | TCTT:TCTT |
| 54 | 8.8 | TCTT:TCTT |
| 55 | 3.6 | /:/ |
| 56 | 9.5 | /:TCTT |
| 57 | 3.5 | /:/ |
| 58 | 7.9 | TCTT:TCTT |
| 59 | 3.5 | /:/ |
| 60 | 3.4 | /:/ |
| 61 | 9.2 | /:TCTT |
| 62 | 8.8 | TCTT:TCTT |
| 63 | 3.7 | /:/ |
| 64 | 7.3 | TCTT:TCTT |
| 65 | 9.3 | /:TCTT |
| 66 | 12.6 | /:TCTT |
| 67 | 10.8 | /:TCTT |
| 68 | 10.5 | /:TCTT |
| 69 | 10.2 | /:TCTT |
| 70 | 10.3 | /:TCTT |
| 71 | 8.1 | TCTT:TCTT |
| 72 | 3.8 | /:/ |
| 73 | 8.8 | TCTT:TCTT |
| 74 | 6.8 | TCTT:TCTT |
| 75 | 8.1 | /:TCTT |
| 76 | 10.4 | /:TCTT |
| 77 | 9.2 | /:TCTT |
| 78 | 10.6 | /:TCTT |
| 79 | 3.9 | /:/ |
| 80 | 8.7 | TCTT:TCTT |
| 81 | 3.6 | /:/ |
| 82 | 8.2 | /:TCTT |
| 83 | 7 | TCTT:TCTT |
| 84 | 0.7 | /:/ |
| 85 | 2.3 | /:/ |
| 86 | 11.5 | /:TCTT |
| 87 | 12.7 | /:TCTT |
| 88 | 12.1 | /:TCTT |
| 89 | 11.6 | /:TCTT |
| 90 | 7.5 | /:TCTT |
| 91 | 11.7 | N.T. |
| 92 | 10.1 | N.T. |
| 93 | 9.5 | N.T. |
| 94 | 6.6 | N.T. |
| 95 | 12.3 | N.T. |
| 96 | 11.6 | N.T. |
| 97 | 9.2 | N.T. |
| 98 | 9.7 | N.T. |
| 99 | 9.7 | N.T. |
| 100 | 1.5 | /:/ |
| 101 | 9.2 | N.T. |
| 102 | 10.2 | N.T. |

| | | |
|---|---|---|
| 103 | 9.1 | N.T. |
| 104 | 3.5 | /:/ |
| 105 | 7.5 | N.T. |
| 106 | 9 | N.T. |
| 107 | 2 | /:/ |
| 108 | 7.6 | N.T. |
| 109 | 10 | N.T. |
| 110 | 10.8 | N.T. |
| 111 | 11.2 | N.T. |
| 112 | 12.2 | N.T. |
| 113 | 2.2 | /:/ |
| 114 | 9.2 | N.T. |
| 115 | 8.4 | N.T. |
| 116 | 10.8 | N.T. |
| 117 | 3.2 | /:/ |
| 118 | 10.2 | N.T. |
| 119 | 4.8 | N.T. |
| 120 | 8.4 | N.T. |
| 121 | 10.5 | N.T. |
| 122 | 9.2 | N.T. |
| 123 | 11.9 | N.T. |
| 124 | 8.2 | N.T. |
| 125 | 10.1 | N.T. |
| 126 | 10.2 | N.T. |
| 127 | 9.8 | N.T. |
| 128 | 7.2 | N.T. |
| 129 | 8.1 | N.T. |
| 130 | 6.9 | N.T. |
| 131 | 9.3 | N.T. |
| 132 | 8.8 | N.T. |
| 133 | 3.8 | N.T. |
| 134 | 3.1 | N.T. |
| 135 | 7.1 | N.T. |
| 136 | 7.7 | N.T. |
| 137 | 9 | N.T. |
| 138 | 3 | N.T. |
| 139 | 9.5 | N.T. |
| 140 | 3.3 | N.T. |
| 141 | 10.6 | N.T. |
| 142 | 3.1 | N.T. |
| 143 | 2.9 | N.T. |
| 144 | 11.1 | N.T. |
| 145 | 8.1 | N.T. |
| 146 | 8.7 | N.T. |
| 147 | 3 | N.T. |
| 148 | 3.2 | N.T. |
| 149 | 9 | N.T. |
| 150 | 14.2 | N.T. |
| 151 | 8.6 | N.T. |
| 152 | 8.8 | N.T. |
| 153 | 7.2 | N.T. |
| 154 | 7.4 | N.T. |

| | | |
|---|---|---|
| 155 | 10.4 | N.T. |
| 156 | 3 | N.T. |
| 157 | 8 | N.T. |
| 158 | 9.7 | N.T. |
| 159 | 8.7 | N.T. |
| 160 | 11.4 | N.T. |
| 161 | 8 | N.T. |
| 162 | 7 | N.T. |
| 163 | 13.1 | N.T. |
| 164 | 8.5 | N.T. |
| 165 | 3.2 | N.T. |
| 166 | 13.5 | N.T. |
| 167 | 8.9 | N.T. |
| 168 | 3 | N.T. |
| 169 | 8 | N.T. |
| 170 | 7.2 | N.T. |
| 171 | 10.5 | N.T. |
| 172 | 12.2 | N.T. |
| 173 | 7.7 | N.T. |
| 174 | 11.6 | N.T. |
| 175 | 12.4 | N.T. |
| 176 | 7.9 | N.T. |
| 177 | 8.3 | N.T. |
| 178 | 7.2 | N.T. |
| 179 | 8.4 | N.T. |
| 180 | 10.3 | N.T. |
| 181 | 9.1 | N.T. |
| 182 | 8.9 | N.T. |
| 183 | 9.8 | N.T. |
| 184 | 3.3 | N.T. |
| 185 | 8 | N.T. |
| 186 | 7.2 | N.T. |
| 187 | 7.2 | N.T. |
| 188 | 8.1 | N.T. |
| 189 | 7.9 | N.T. |
| 190 | 3.4 | N.T. |
| 191 | 3 | N.T. |
| 192 | 7.7 | N.T. |
| 193 | 2.7 | N.T. |
| 194 | 7.2 | N.T. |
| 195 | 3.4 | N.T. |
| 196 | 1.5 | N.T. |
| 197 | 8.1 | N.T. |
| 198 | 3.2 | N.T. |

Note: N.T. indicates that the genotypes of these $F_2$ individuals were not tested.

**Table S3. The SNPs on chromosome 1 significantly associated with the salt tolerance of maize.**

| Chr. | Position | REF_ Allele | ALT_ Allele | Recessive Allele | Salt sensitive REF_COUNT | Salt sensitive ALT_COUNT | Salt tolerance REF_COUNT | Salt tolerance ALT_COUNT | Association Probability | Significance |
|---|---|---|---|---|---|---|---|---|---|---|
| 1 | 52151799 | C | G | C | 22 | 0 | 4 | 5 | 0.059681992 | yes |
| 1 | 64086392 | G | A | G | 26 | 0 | 7 | 7 | 0.064316284 | yes |
| 1 | 71593705 | C | T | T | 0 | 35 | 10 | 12 | 0.070295559 | yes |
| 1 | 71593761 | C | T | T | 0 | 32 | 8 | 11 | 0.050242025 | yes |
| 1 | 71593805 | G | A | A | 0 | 29 | 9 | 9 | 0.074698236 | yes |
| 1 | 72021208 | G | A | G | 16 | 0 | 4 | 18 | 0.059105307 | yes |
| 1 | 72021245 | G | T | G | 20 | 0 | 6 | 17 | 0.081314104 | yes |
| 1 | 79725383 | A | G | A | 24 | 0 | 10 | 10 | 0.058260403 | yes |
| 1 | 86595957 | C | T | C | 22 | 0 | 9 | 16 | 0.086109792 | yes |
| 1 | 91885872 | G | C | G | 26 | 0 | 5 | 16 | 0.117411663 | yes |
| 1 | 91885983 | A | G | A | 23 | 0 | 7 | 9 | 0.070035992 | yes |
| 1 | 110645805 | A | G | G | 0 | 21 | 11 | 5 | 0.080627565 | yes |
| 1 | 116097032 | A | T | T | 0 | 21 | 18 | 10 | 0.082234062 | yes |
| 1 | 116097033 | A | G | G | 0 | 21 | 18 | 10 | 0.082234062 | yes |
| 1 | 116097037 | G | C | C | 0 | 21 | 18 | 10 | 0.082234062 | yes |
| 1 | 155633412 | G | T | G | 36 | 0 | 9 | 21 | 0.1768566 | yes |
| 1 | 157069328 | T | C | T | 42 | 0 | 8 | 37 | 0.21650853 | yes |
| 1 | 161475338 | C | T | C | 18 | 0 | 9 | 19 | 0.068283023 | yes |
| 1 | 162436260 | T | C | T | 22 | 0 | 9 | 15 | 0.083344887 | yes |
| 1 | 164509417 | G | A | A | 0 | 72 | 29 | 35 | 0.116867669 | yes |
| 1 | 166685948 | C | T | C | 19 | 0 | 4 | 9 | 0.067878327 | yes |
| 1 | 166686028 | G | A | G | 32 | 0 | 4 | 21 | 0.15573444 | yes |
| 1 | 166686187 | C | G | C | 32 | 0 | 7 | 7 | 0.084147006 | yes |
| 1 | 170324145 | C | G | G | 0 | 15 | 45 | 9 | 0.053605391 | yes |
| 1 | 170324148 | G | A | A | 0 | 15 | 45 | 9 | 0.053605391 | yes |
| 1 | 170324159 | G | A | A | 0 | 18 | 41 | 9 | 0.070885171 | yes |
| 1 | 170324163 | C | A | A | 0 | 17 | 39 | 9 | 0.065043492 | yes |
| 1 | 170324172 | C | A | A | 0 | 18 | 35 | 10 | 0.070876149 | yes |
| 1 | 170596435 | A | C | C | 0 | 15 | 24 | 3 | 0.053602837 | yes |
| 1 | 172039747 | C | T | T | 0 | 33 | 24 | 21 | 0.115389321 | yes |
| 1 | 172040046 | C | T | T | 0 | 30 | 27 | 11 | 0.14273018 | yes |
| 1 | 172040820 | C | T | T | 0 | 16 | 13 | 7 | 0.052919826 | yes |
| 1 | 172838382 | G | A | A | 0 | 22 | 10 | 4 | 0.086974083 | yes |
| 1 | 173233501 | C | T | T | 0 | 15 | 15 | 6 | 0.051654083 | yes |
| 1 | 173233592 | A | G | G | 0 | 17 | 11 | 4 | 0.061319757 | yes |
| 1 | 173247642 | G | A | A | 0 | 21 | 13 | 8 | 0.075409853 | yes |
| 1 | 174660843 | G | A | A | 0 | 22 | 17 | 6 | 0.093134344 | yes |
| 1 | 174660871 | A | C | C | 0 | 34 | 27 | 9 | 0.167729454 | yes |
| 1 | 174671119 | G | T | T | 0 | 45 | 45 | 20 | 0.234218245 | yes |
| 1 | 174958614 | G | A | A | 0 | 19 | 8 | 3 | 0.06876661 | yes |
| 1 | 175250431 | G | A | A | 0 | 23 | 24 | 4 | 0.100816088 | yes |
| 1 | 177058885 | G | A | A | 0 | 15 | 16 | 4 | 0.053212925 | yes |
| 1 | 177058893 | G | T | T | 0 | 15 | 15 | 4 | 0.052995829 | yes |
| 1 | 177058926 | C | G | G | 0 | 15 | 16 | 3 | 0.053428895 | yes |
| 1 | 177058935 | T | C | C | 0 | 17 | 18 | 5 | 0.064642903 | yes |
| 1 | 178347857 | T | G | T | 32 | 0 | 7 | 11 | 0.126614123 | yes |
| 1 | 178347875 | T | C | T | 36 | 0 | 7 | 16 | 0.172812085 | yes |
| 1 | 179368000 | A | T | T | 0 | 19 | 10 | 7 | 0.058239622 | yes |
| 1 | 179368044 | C | A | A | 0 | 20 | 14 | 6 | 0.078464752 | yes |
| 1 | 180007423 | C | T | C | 21 | 0 | 7 | 9 | 0.061638988 | yes |
| 1 | 180007607 | A | C | A | 21 | 0 | 8 | 14 | 0.078607565 | yes |
| 1 | 180008313 | T | C | T | 34 | 0 | 6 | 16 | 0.163814275 | yes |
| 1 | 180008350 | C | G | C | 34 | 0 | 5 | 16 | 0.165656665 | yes |
| 1 | 180008500 | G | T | G | 31 | 0 | 4 | 9 | 0.132348103 | yes |
| 1 | 180363865 | A | G | A | 21 | 0 | 4 | 9 | 0.07844372 | yes |

| | | | | | | | | | | |
|---|---|---|---|---|---|---|---|---|---|---|
| 1 | 180364417 | T | C | T | 20 | 0 | 5 | 15 | 0.080978607 | yes |
| 1 | 180364501 | C | A | A | 0 | 20 | 12 | 6 | 0.074493418 | yes |
| 1 | 180364504 | T | G | T | 21 | 0 | 6 | 11 | 0.076756276 | yes |
| 1 | 180364552 | G | A | G | 22 | 0 | 6 | 9 | 0.072536448 | yes |
| 1 | 180364558 | C | T | T | 0 | 20 | 10 | 6 | 0.06787576 | yes |
| 1 | 180364885 | G | T | T | 0 | 28 | 9 | 3 | 0.121882725 | yes |
| 1 | 180364899 | C | T | C | 32 | 0 | 3 | 8 | 0.139559009 | yes |
| 1 | 180551216 | A | G | G | 0 | 29 | 7 | 9 | 0.052532913 | yes |
| 1 | 180636330 | G | A | A | 0 | 22 | 18 | 6 | 0.09366544 | yes |
| 1 | 180636793 | T | C | C | 0 | 28 | 18 | 5 | 0.130542045 | yes |
| 1 | 180636808 | T | C | C | 0 | 32 | 21 | 6 | 0.155316434 | yes |
| 1 | 180636809 | C | G | G | 0 | 32 | 21 | 6 | 0.155316434 | yes |
| 1 | 180715697 | G | A | G | 68 | 0 | 19 | 24 | 0.292780034 | yes |
| 1 | 180715716 | T | A | T | 42 | 0 | 12 | 13 | 0.13328929 | yes |
| 1 | 180715767 | G | T | G | 46 | 0 | 12 | 20 | 0.220526259 | yes |
| 1 | 180717953 | G | C | G | 77 | 0 | 19 | 38 | 0.411344566 | yes |
| 1 | 180718001 | G | A | A | 0 | 84 | 37 | 17 | 0.44791077 | yes |
| 1 | 180718146 | G | C | G | 60 | 0 | 18 | 32 | 0.313944005 | yes |
| 1 | 180839584 | G | A | A | 0 | 21 | 9 | 4 | 0.07844372 | yes |
| 1 | 180839627 | T | C | C | 0 | 24 | 14 | 5 | 0.103550489 | yes |
| 1 | 181332187 | G | A | G | 32 | 0 | 8 | 11 | 0.117254641 | yes |
| 1 | 181332232 | C | G | C | 43 | 0 | 13 | 16 | 0.162656097 | yes |
| 1 | 181332262 | C | T | C | 43 | 0 | 14 | 16 | 0.149968866 | yes |
| 1 | 181332275 | A | G | G | 0 | 46 | 21 | 16 | 0.1947211 | yes |
| 1 | 181332310 | A | G | G | 0 | 42 | 16 | 13 | 0.158274063 | yes |
| 1 | 181332371 | C | T | C | 36 | 0 | 14 | 12 | 0.07481839 | yes |
| 1 | 181332373 | A | G | A | 36 | 0 | 14 | 12 | 0.07481839 | yes |
| 1 | 181332377 | C | T | T | 0 | 35 | 12 | 13 | 0.08364642 | yes |
| 1 | 181332391 | G | A | G | 49 | 0 | 20 | 14 | 0.056252825 | yes |
| 1 | 181332398 | C | T | C | 49 | 0 | 20 | 14 | 0.056252825 | yes |
| 1 | 181335736 | A | G | A | 78 | 0 | 19 | 12 | 0.068555391 | yes |
| 1 | 181336512 | T | C | T | 43 | 0 | 14 | 13 | 0.107628544 | yes |
| 1 | 181337145 | T | G | T | 54 | 0 | 13 | 8 | 0.060217104 | yes |
| 1 | 181337295 | A | T | A | 66 | 0 | 23 | 23 | 0.199444041 | yes |
| 1 | 181337297 | A | T | A | 67 | 0 | 22 | 23 | 0.221314215 | yes |
| 1 | 181337524 | A | G | A | 36 | 0 | 19 | 23 | 0.137073494 | yes |
| 1 | 181337579 | T | C | T | 34 | 0 | 20 | 19 | 0.084133199 | yes |
| 1 | 181337580 | A | G | A | 35 | 0 | 20 | 19 | 0.087185457 | yes |
| 1 | 181338420 | G | T | G | 85 | 0 | 27 | 17 | 0.050131306 | yes |
| 1 | 181339202 | G | T | G | 67 | 0 | 24 | 24 | 0.202924695 | yes |
| 1 | 181339366 | C | T | C | 73 | 0 | 26 | 26 | 0.221984123 | yes |
| 1 | 181650735 | C | T | C | 31 | 0 | 7 | 9 | 0.103995619 | yes |
| 1 | 181650873 | C | T | C | 34 | 0 | 7 | 9 | 0.116731662 | yes |
| 1 | 181911078 | G | C | C | 0 | 18 | 39 | 3 | 0.070885415 | yes |
| 1 | 181911542 | G | A | A | 0 | 20 | 10 | 8 | 0.057117427 | yes |
| 1 | 181911571 | C | T | T | 0 | 17 | 7 | 4 | 0.050932459 | yes |
| 1 | 181911572 | A | G | G | 0 | 17 | 7 | 4 | 0.050932459 | yes |
| 1 | 181913076 | T | C | C | 0 | 49 | 64 | 12 | 0.258113328 | yes |
| 1 | 181913116 | T | G | G | 0 | 48 | 32 | 12 | 0.251811999 | yes |
| 1 | 181913216 | T | G | G | 0 | 35 | 45 | 10 | 0.174155504 | yes |
| 1 | 181913323 | A | G | G | 0 | 23 | 49 | 14 | 0.100830379 | yes |
| 1 | 182126400 | C | G | C | 58 | 0 | 21 | 14 | 0.055537502 | yes |
| 1 | 182127345 | G | A | G | 72 | 0 | 28 | 42 | 0.372016595 | yes |
| 1 | 182127632 | A | T | A | 83 | 0 | 32 | 48 | 0.431144012 | yes |
| 1 | 182128861 | G | C | C | 0 | 123 | 81 | 38 | 0.619337247 | yes |
| 1 | 182128929 | G | C | C | 0 | 56 | 39 | 16 | 0.298457497 | yes |
| 1 | 182128933 | A | G | G | 0 | 46 | 27 | 12 | 0.2380172 | yes |
| 1 | 182128937 | A | T | T | 0 | 41 | 13 | 11 | 0.143263273 | yes |

| | | | | | | | | | | |
|---|---|---|---|---|---|---|---|---|---|---|
| 1 | 182129206 | G | A | G | 21 | 0 | 9 | 10 | 0.054867059 | yes |
| 1 | 182129291 | G | A | G | 98 | 0 | 37 | 84 | 0.516228175 | yes |
| 1 | 182319543 | C | T | C | 23 | 0 | 3 | 5 | 0.072844063 | yes |
| 1 | 182319544 | T | G | T | 23 | 0 | 3 | 5 | 0.072844063 | yes |
| 1 | 182319545 | A | T | A | 23 | 0 | 3 | 5 | 0.072844063 | yes |
| 1 | 182319546 | T | A | T | 22 | 0 | 3 | 5 | 0.068467433 | yes |
| 1 | 182319552 | T | C | T | 22 | 0 | 3 | 5 | 0.068467433 | yes |
| 1 | 182319645 | G | A | G | 43 | 0 | 7 | 10 | 0.16875535 | yes |
| 1 | 182319749 | G | A | G | 64 | 0 | 10 | 16 | 0.300419679 | yes |
| 1 | 182649853 | T | C | T | 25 | 0 | 6 | 19 | 0.112124512 | yes |
| 1 | 182651021 | A | C | A | 18 | 0 | 8 | 18 | 0.068549142 | yes |
| 1 | 182651923 | T | C | T | 32 | 0 | 6 | 39 | 0.155840722 | yes |
| 1 | 182651945 | C | A | C | 33 | 0 | 9 | 31 | 0.161894276 | yes |
| 1 | 182651968 | C | A | A | 0 | 35 | 28 | 10 | 0.173735996 | yes |
| 1 | 182651978 | A | T | T | 0 | 34 | 30 | 11 | 0.167725228 | yes |
| 1 | 182651979 | A | T | T | 0 | 35 | 30 | 11 | 0.17381011 | yes |
| 1 | 182652065 | C | T | C | 22 | 0 | 8 | 12 | 0.076340085 | yes |
| 1 | 182652068 | T | G | T | 21 | 0 | 8 | 12 | 0.071482046 | yes |
| 1 | 182652069 | A | T | A | 20 | 0 | 8 | 12 | 0.066651305 | yes |
| 1 | 182652079 | A | G | A | 18 | 0 | 6 | 19 | 0.070332349 | yes |
| 1 | 182823100 | C | T | T | 0 | 17 | 15 | 5 | 0.063655878 | yes |
| 1 | 182823413 | G | C | G | 47 | 0 | 15 | 46 | 0.24630594 | yes |
| 1 | 182875143 | C | T | C | 25 | 0 | 9 | 19 | 0.108857456 | yes |
| 1 | 182875226 | T | C | C | 0 | 27 | 20 | 5 | 0.12491232 | yes |
| 1 | 182875292 | C | T | C | 34 | 0 | 7 | 21 | 0.166979641 | yes |
| 1 | 182875301 | C | T | C | 36 | 0 | 7 | 21 | 0.179085736 | yes |
| 1 | 182875330 | G | A | A | 0 | 30 | 20 | 7 | 0.142231334 | yes |
| 1 | 182875470 | A | G | A | 25 | 0 | 13 | 27 | 0.111268545 | yes |
| 1 | 182875660 | G | A | A | 0 | 16 | 12 | 4 | 0.056958395 | yes |
| 1 | 182875665 | G | C | C | 0 | 15 | 14 | 5 | 0.05192075 | yes |
| 1 | 182929327 | G | T | T | 0 | 17 | 20 | 10 | 0.06241709 | yes |
| 1 | 293535400 | C | T | C | 51 | 0 | 5 | 6 | 0.168967349 | yes |

Note: The positions of SNPs were based on B73 RefGen_v3 reference genome.

**Table S4. KASP primers used for fine mapping of *qST1* in this study.**

| Marker Name | SNPs in | | Position in B73 RefGen_v3 | KASP primer sequences | | |
|---|---|---|---|---|---|---|
| | Jing724 | D9H | | AlleleFAM | AlleleHEX | Common |
| ST1 | A | G | chr1: 171730547 | CCTTCACTCTACTCGCACGACA | CTTCACTCTACTCGCACGACG | CTCCTTTCCTCTCCCTTTCCCCAT |
| ST2 | A | G | chr1: 173724075 | GGTGACTTTATTTGTTCAACACCTAT | GGTGACTTTATTTGTTCAACACCTAC | ATGTTAGACTACTTGCTTGGGCATCATAT |
| ST3 | G | A | chr1: 176151909 | CCCGAGATGGAAAGGGAGGATA | CCGAGATGGAAAGGGAGGATG | CGGGTCATTAGAAACATATGGATTTGGTT |
| ST4 | A | G | chr1: 178647424 | CAAGATATAGATGGTGGGGTATCTGT | AAGATATAGATGGTGGGGTATCTGC | CGCATTGCATCCCAATTCCAGTGTA |
| ST5 | G | A | chr1: 180015649 | ACGAACACCAGCAGCAGGAGT | CGAACACCAGCAGCAGGAGC | CCACCGGAGGCACCACGGAA |
| ST6 | A | G | chr1: 180428198 | AAGATGCAGGTAGCAGAAGCACAA | GATGCAGGTAGCAGAAGCACAG | GCTGGACATTCTTAAGTTAATTCGGCAAT |
| ST10 | A | C | chr1: 180841928 | TCGTAGGCACTTTGGTCTATAGAAA | CGTAGGCACTTTGGTCTATAGAAC | TAATCTAGTACTCATTGGCTGCCCTTTTA |
| ST11 | G | T | chr1: 180862086 | GTATTCCTACTTGCGCAGGGTCA | ATTCCTACTTGCGCAGGGTCC | CTATGGACCCAAATGGGTTCAACTCTT |
| ST12 | A | G | chr1: 180972079 | GACCCACTAAGCCTTGTATCGG | GGACCCACTAAGCCTTGTATCGA | CAAGAGGCACATTGACGACTCCTTT |
| ST13 | G | A | chr1: 180991179 | GCATCTGTTCTGTCTCCATGCCA | CATCTGTTCTGTCTCCATGCCG | GTCCCGGTCCCTTCGCTTACTA |
| ST7 | G | A | chr1: 181238642 | ATAATCCGTCATCATGTGGTGTAATCT | AATCCGTCATCATGTGGTGTAATCC | AGCATCTACCTCATTCATGAAGCCAAATT |
| ST8 | G | A | chr1: 182130289 | GAAGTAACAGAGAAGCTAGCTGT | GAAGTAACAGAGAAGCTAGCTGC | GAGACCCATCTTCAGGAAGTTGCTT |
| ST9 | G | A | chr1: 182875470 | ACGAGGGTATTTACAAAGCCATTGAA | CGAGGGTATTTACAAAGCCATTGAG | TGCCGGCAAAGGATACTCATCCAT |
| 4-bp InDel | TCTT | - | chr1: 180962923 | CAGGTAGAGCTGCCACTCT | CTCAGGTAGAGCTGCCACTCC | TGCAGTCGGAGGGCTCCAACAT |

**Table S5. Recombinant events identified within the salt-sensitive plants of Jing724 and D9H $F_2$ population for fine mapping *qST1*.**

| $F_2$ plants | KASP markers | | | | | | | | | | | | | Root length (cm) |
|---|---|---|---|---|---|---|---|---|---|---|---|---|---|---|
| | ST1 | ST2 | ST3 | ST4 | ST5 | ST6 | ST10 | ST11 | ST12 | ST13 | ST7 | ST8 | ST9 | |
| Jing724 | A:A | A:A | G:G | A:A | G:G | A:A | A:A | G:G | A:A | G:G | G:G | G:G | G:G | |
| D9H | G:G | G:G | A:A | G:G | A:A | G:G | C:C | T:T | G:G | A:A | A:A | A:A | A:A | |
| rec-1 | G:G | G:G | A:A | G:G | A:A | G:G | C:C | T:T | G:G | G:A | G:A | G:A | G:A | 3 |
| rec-2 | G:G | G:G | A:A | G:G | A:A | G:G | C:C | T:T | G:G | G:A | G:A | G:A | G:A | 2.3 |
| rec-3 | G:G | G:G | A:A | G:G | A:A | G:G | C:C | T:T | G:G | A:A | G:A | G:A | G:A | 3.5 |
| rec-4 | G:G | G:G | A:A | G:G | A:A | G:G | C:C | T:T | G:G | A:A | G:A | G:A | G:A | 3.7 |
| rec-5 | G:G | G:G | A:A | G:G | A:A | G:G | C:C | T:T | G:G | A:A | A:A | G:A | G:A | 1.9 |
| rec-6 | G:G | G:G | A:A | G:G | A:A | G:G | C:C | T:T | G:G | A:A | A:A | G:A | G:A | 2.1 |
| rec-7 | G:G | G:G | A:A | G:G | A:A | G:G | C:C | T:T | G:G | A:A | A:A | G:A | G:A | 3.6 |
| rec-8 | G:G | G:G | A:A | G:G | A:A | G:G | C:C | T:T | G:G | A:A | A:A | G:A | G:A | 3.6 |
| rec-9 | G:A | G:A | A:A | G:G | A:A | G:G | C:C | T:T | G:G | A:A | A:A | G:A | G:A | 2.6 |
| rec-10 | G:G | G:G | A:A | G:G | A:A | G:G | C:C | T:T | G:G | A:A | A:A | G:A | G:A | 3.7 |
| rec-11 | G:G | G:G | A:A | G:G | A:A | G:G | C:C | T:T | G:G | A:A | A:A | A:A | G:A | 3.2 |
| rec-12 | G:G | G:G | A:A | G:G | A:A | G:G | C:C | T:T | G:G | A:A | A:A | A:A | G:A | 2.5 |
| rec-13 | G:G | G:G | A:A | G:G | A:A | G:G | C:C | T:T | G:G | A:A | A:A | A:A | G:A | 3.3 |
| rec-14 | G:G | G:G | A:A | G:G | A:A | G:G | C:C | T:T | G:G | A:A | A:A | A:A | G:A | 2.3 |
| rec-15 | G:G | G:G | A:A | G:G | A:A | G:G | C:C | T:T | G:G | A:A | A:A | A:A | G:A | 3.5 |
| rec-16 | G:G | G:G | A:A | G:G | A:A | G:G | C:C | T:T | G:G | A:A | A:A | A:A | G:A | 2.2 |
| rec-17 | G:G | G:G | A:A | G:G | A:A | G:G | C:C | T:T | G:G | A:A | A:A | A:A | G:A | 2.8 |
| rec-18 | G:G | G:G | A:A | G:G | A:A | G:G | C:C | T:T | G:G | A:A | A:A | A:A | G:A | 3.6 |
| rec-19 | G:G | G:G | A:A | G:G | A:A | G:G | C:C | T:T | G:G | A:A | A:A | A:A | G:A | 2.8 |
| rec-20 | G:A | G:A | A:A | G:G | A:A | G:G | C:C | T:T | G:G | A:A | A:A | A:A | G:A | 3.2 |
| rec-21 | G:A | G:A | G:A | G:A | G:A | G:A | C:A | G:T | G:G | A:A | A:A | A:A | A:A | 3.6 |
| rec-22 | G:A | G:A | G:A | G:A | G:A | G:A | C:A | T:T | G:G | A:A | A:A | A:A | A:A | 1.5 |
| rec-23 | G:A | G:A | G:A | G:A | G:A | G:A | C:A | T:T | G:G | A:A | A:A | A:A | A:A | 2.9 |
| rec-24 | G:A | G:A | G:A | G:A | G:A | G:A | C:C | T:T | G:G | A:A | A:A | A:A | A:A | 3.5 |
| rec-25 | G:A | G:A | G:A | G:A | G:A | G:A | C:C | T:T | G:G | A:A | A:A | A:A | A:A | 3.5 |
| rec-26 | G:A | G:A | G:A | G:A | G:A | G:A | C:C | T:T | G:G | A:A | A:A | A:A | A:A | 2.7 |
| rec-27 | G:A | G:A | G:A | G:A | G:A | G:A | C:C | T:T | G:G | A:A | A:A | A:A | A:A | 2.6 |
| rec-28 | G:A | G:A | G:A | G:A | G:A | G:A | C:C | T:T | G:G | A:A | A:A | A:A | A:A | 2.8 |
| rec-29 | G:A | G:A | G:A | G:A | G:A | G:A | C:C | T:T | G:G | A:A | A:A | A:A | A:A | 3.1 |
| rec-30 | G:A | G:A | G:A | G:A | G:A | G:A | C:C | T:T | G:G | A:A | A:A | A:A | A:A | 3.3 |
| rec-31 | G:A | G:A | G:A | G:A | G:A | G:A | C:C | T:T | G:G | A:A | A:A | A:A | A:A | 2.6 |
| rec-32 | G:A | G:A | G:A | G:A | G:A | G:A | C:C | T:T | G:G | A:A | A:A | A:A | A:A | 2.9 |
| rec-33 | G:A | G:A | G:A | G:A | G:A | G:A | C:C | T:T | G:G | A:A | A:A | A:A | A:A | 3 |
| rec-34 | G:A | G:A | G:A | G:A | G:A | G:A | C:C | T:T | G:G | A:A | A:A | A:A | A:A | 2.5 |
| rec-35 | G:A | G:A | G:A | G:A | G:A | G:G | C:C | T:T | G:G | A:A | A:A | A:A | A:A | 2.2 |
| rec-36 | G:A | G:A | G:A | G:A | G:A | G:G | C:C | T:T | G:G | A:A | A:A | A:A | A:A | 3.5 |
| rec-37 | G:A | G:A | G:A | G:A | G:A | G:G | C:C | T:T | G:G | A:A | A:A | A:A | A:A | 2.3 |
| rec-38 | G:A | G:A | G:A | G:A | G:A | G:G | C:C | T:T | G:G | A:A | A:A | A:A | A:A | 2.4 |
| rec-39 | G:A | G:A | G:A | G:A | A:A | G:G | C:C | T:T | G:G | A:A | A:A | A:A | A:A | 3.2 |
| rec-40 | G:A | G:A | G:A | G:A | A:A | G:G | C:C | T:T | G:G | A:A | A:A | A:A | A:A | 2.9 |
| rec-41 | G:A | G:A | G:A | G:A | A:A | G:G | C:C | T:T | G:G | A:A | A:A | A:A | A:A | 3.2 |
| rec-42 | G:A | G:A | G:A | G:A | A:A | G:G | C:C | T:T | G:G | A:A | A:A | A:A | A:A | 2.3 |
| rec-43 | G:A | G:A | G:A | G:A | A:A | G:G | C:C | T:T | G:G | A:A | A:A | A:A | A:A | 3.5 |
| rec-44 | G:A | G:A | G:A | G:A | A:A | G:G | C:C | T:T | G:G | A:A | A:A | A:A | A:A | 3.3 |
| rec-45 | G:A | G:A | G:A | G:A | A:A | G:G | C:C | T:T | G:G | A:A | A:A | A:A | A:A | 3.6 |
| rec-46 | G:A | G:A | G:A | G:A | A:A | G:G | C:C | T:T | G:G | A:A | A:A | A:A | A:A | 2.5 |
| rec-47 | G:A | G:A | G:A | G:A | A:A | G:G | C:C | T:T | G:G | A:A | A:A | A:A | A:A | 3.2 |
| rec-48 | G:A | G:A | G:A | G:A | A:A | G:G | C:C | T:T | G:G | A:A | A:A | A:A | A:A | 2.7 |
| rec-49 | G:A | G:A | G:A | G:A | A:A | G:G | C:C | T:T | G:G | A:A | A:A | A:A | A:A | 2.7 |
| rec-50 | G:A | G:A | G:A | G:A | A:A | G:G | C:C | T:T | G:G | A:A | A:A | A:A | A:A | 1.8 |
| rec-51 | G:A | G:A | G:A | G:A | A:A | G:G | C:C | T:T | G:G | A:A | A:A | A:A | A:A | 3.5 |

| | | | | | | | | | | | | | | |
|---|---|---|---|---|---|---|---|---|---|---|---|---|---|---|
| rec-52 | G:A | G:A | G:A | G:A | A:A | G:G | C:C | T:T | G:G | A:A | A:A | A:A | A:A | 3.4 |
| rec-53 | G:A | G:A | G:A | G:A | A:A | G:G | C:C | T:T | G:G | A:A | A:A | A:A | A:A | 2.3 |
| rec-54 | G:A | G:A | G:A | G:A | A:A | G:G | C:C | T:T | G:G | A:A | A:A | A:A | A:A | 2.4 |
| rec-55 | G:A | G:A | G:A | G:A | A:A | G:G | C:C | T:T | G:G | A:A | A:A | A:A | A:A | 3 |
| rec-56 | G:A | G:A | G:A | G:A | A:A | G:G | C:C | T:T | G:G | A:A | A:A | A:A | A:A | 2.8 |
| rec-57 | G:A | G:A | G:A | G:A | A:A | G:G | C:C | T:T | G:G | A:A | A:A | A:A | A:A | 1.9 |
| rec-58 | G:A | G:A | G:A | G:A | A:A | G:G | C:C | T:T | G:G | A:A | A:A | A:A | A:A | 3.5 |
| rec-59 | G:A | G:A | G:A | G:A | A:A | G:G | C:C | T:T | G:G | A:A | A:A | A:A | A:A | 3.1 |
| rec-60 | G:A | G:A | G:A | G:G | A:A | G:G | C:C | T:T | G:G | A:A | A:A | A:A | A:A | 2.5 |
| rec-61 | G:A | G:A | G:A | G:G | A:A | G:G | C:C | T:T | G:G | A:A | A:A | A:A | A:A | 3.2 |
| rec-62 | G:A | G:A | G:A | G:G | A:A | G:G | C:C | T:T | G:G | A:A | A:A | A:A | A:A | 3.1 |
| rec-63 | G:A | G:A | G:A | G:G | A:A | G:G | C:C | T:T | G:G | A:A | A:A | A:A | A:A | 3.4 |
| rec-64 | G:A | G:A | G:A | G:G | A:A | G:G | C:C | T:T | G:G | A:A | A:A | A:A | A:A | 2.4 |
| rec-65 | G:A | G:A | G:A | G:G | A:A | G:G | C:C | T:T | G:G | A:A | A:A | A:A | A:A | 2.2 |
| rec-66 | G:A | G:A | G:A | G:G | A:A | G:G | C:C | T:T | G:G | A:A | A:A | A:A | A:A | 3.2 |
| rec-67 | G:A | G:A | G:A | G:G | A:A | G:G | C:C | T:T | G:G | A:A | A:A | A:A | A:A | 2.6 |
| rec-68 | G:A | G:A | G:A | G:G | A:A | G:G | C:C | T:T | G:G | A:A | A:A | A:A | A:A | 3 |
| rec-69 | G:A | G:A | A:A | G:G | A:A | G:G | C:C | T:T | G:G | A:A | A:A | A:A | A:A | 3.1 |
| rec-70 | G:A | G:A | A:A | G:G | A:A | G:G | C:C | T:T | G:G | A:A | A:A | A:A | A:A | 3 |
| rec-71 | G:A | G:A | A:A | G:G | A:A | G:G | C:C | T:T | G:G | A:A | A:A | A:A | A:A | 3 |
| rec-72 | G:A | G:A | A:A | G:G | A:A | G:G | C:C | T:T | G:G | A:A | A:A | A:A | A:A | 3.5 |
| rec-73 | G:A | G:A | A:A | G:G | A:A | G:G | C:C | T:T | G:G | A:A | A:A | A:A | A:A | 1.5 |
| rec-74 | G:A | G:A | A:A | G:G | A:A | G:G | C:C | T:T | G:G | A:A | A:A | A:A | A:A | 3.5 |
| rec-75 | G:A | G:A | A:A | G:G | A:A | G:G | C:C | T:T | G:G | A:A | A:A | A:A | A:A | 3.6 |
| rec-76 | G:A | G:A | A:A | G:G | A:A | G:G | C:C | T:T | G:G | A:A | A:A | A:A | A:A | 3.6 |
| rec-77 | G:A | G:A | A:A | G:G | A:A | G:G | C:C | T:T | G:G | A:A | A:A | A:A | A:A | 3.2 |
| rec-78 | G:A | G:A | A:A | G:G | A:A | G:G | C:C | T:T | G:G | A:A | A:A | A:A | A:A | 1.9 |
| rec-79 | G:A | G:A | A:A | G:G | A:A | G:G | C:C | T:T | G:G | A:A | A:A | A:A | A:A | 2.8 |
| rec-80 | G:A | G:A | A:A | G:G | A:A | G:G | C:C | T:T | G:G | A:A | A:A | A:A | A:A | 2.5 |
| rec-81 | G:A | G:A | A:A | G:G | A:A | G:G | C:C | T:T | G:G | A:A | A:A | A:A | A:A | 2.5 |
| rec-82 | G:A | G:A | A:A | G:G | A:A | G:G | C:C | T:T | G:G | A:A | A:A | A:A | A:A | 3.1 |
| rec-83 | G:A | G:A | A:A | G:G | A:A | G:G | C:C | T:T | G:G | A:A | A:A | A:A | A:A | 3.3 |
| rec-84 | G:A | G:A | A:A | G:G | A:A | G:G | C:C | T:T | G:G | A:A | A:A | A:A | A:A | 3.5 |
| rec-85 | G:A | G:A | A:A | G:G | A:A | G:G | C:C | T:T | G:G | A:A | A:A | A:A | A:A | 3.5 |
| rec-86 | G:A | G:A | A:A | G:G | A:A | G:G | C:C | T:T | G:G | A:A | A:A | A:A | A:A | 2.5 |
| rec-87 | G:A | G:A | A:A | G:G | A:A | G:G | C:C | T:T | G:G | A:A | A:A | A:A | A:A | 2.2 |
| rec-88 | G:A | G:A | A:A | G:G | A:A | G:G | C:C | T:T | G:G | A:A | A:A | A:A | A:A | 2.9 |
| rec-89 | G:A | G:A | A:A | G:G | A:A | G:G | C:C | T:T | G:G | A:A | A:A | A:A | A:A | 3.3 |
| rec-90 | G:A | G:A | A:A | G:G | A:A | G:G | C:C | T:T | G:G | A:A | A:A | A:A | A:A | 3.7 |
| rec-91 | G:A | G:A | A:A | G:G | A:A | G:G | C:C | T:T | G:G | A:A | A:A | A:A | A:A | 2.5 |
| rec-92 | G:A | G:A | A:A | G:G | A:A | G:G | C:C | T:T | G:G | A:A | A:A | A:A | A:A | 3 |
| rec-93 | G:A | G:A | A:A | G:G | A:A | G:G | C:C | T:T | G:G | A:A | A:A | A:A | A:A | 3.7 |
| rec-94 | G:A | G:A | A:A | G:G | A:A | G:G | C:C | T:T | G:G | A:A | A:A | A:A | A:A | 2.8 |
| rec-95 | G:A | G:A | A:A | G:G | A:A | G:G | C:C | T:T | G:G | A:A | A:A | A:A | A:A | 2.5 |
| rec-96 | G:A | G:A | A:A | G:G | A:A | G:G | C:C | T:T | G:G | A:A | A:A | A:A | A:A | 2 |
| rec-97 | G:A | G:A | A:A | G:G | A:A | G:G | C:C | T:T | G:G | A:A | A:A | A:A | A:A | 3.3 |
| rec-98 | G:A | G:A | A:A | G:G | A:A | G:G | C:C | T:T | G:G | A:A | A:A | A:A | A:A | 2 |
| rec-99 | G:A | G:A | A:A | G:G | A:A | G:G | C:C | T:T | G:G | A:A | A:A | A:A | A:A | 2.8 |
| rec-100 | G:A | G:A | A:A | G:G | A:A | G:G | C:C | T:T | G:G | A:A | A:A | A:A | A:A | 2.3 |
| rec-101 | G:A | G:A | A:A | G:G | A:A | G:G | C:C | T:T | G:G | A:A | A:A | A:A | A:A | 3.2 |
| rec-102 | G:A | G:A | A:A | G:G | A:A | G:G | C:C | T:T | G:G | A:A | A:A | A:A | A:A | 3.3 |
| rec-103 | G:A | G:A | A:A | G:G | A:A | G:G | C:C | T:T | G:G | A:A | A:A | A:A | A:A | 3.4 |
| rec-104 | G:A | G:A | A:A | G:G | A:A | G:G | C:C | T:T | G:G | A:A | A:A | A:A | A:A | 2.7 |
| rec-105 | G:A | G:A | A:A | G:G | A:A | G:G | C:C | T:T | G:G | A:A | A:A | A:A | A:A | 3.5 |
| rec-106 | G:A | G:A | A:A | G:G | A:A | G:G | C:C | T:T | G:G | A:A | A:A | A:A | A:A | 3.2 |
| rec-107 | G:A | G:A | A:A | G:G | A:A | G:G | C:C | T:T | G:G | A:A | A:A | A:A | A:A | 3.5 |
| rec-108 | G:A | G:A | A:A | G:G | A:A | G:G | C:C | T:T | G:G | A:A | A:A | A:A | A:A | 2 |

| | | | | | | | | | | | | | | |
|---|---|---|---|---|---|---|---|---|---|---|---|---|---|---|
| rec-109 | G:A | G:A | A:A | G:G | A:A | G:G | C:C | T:T | G:G | A:A | A:A | A:A | A:A | 2.7 |
| rec-110 | G:A | G:A | A:A | G:G | A:A | G:G | C:C | T:T | G:G | A:A | A:A | A:A | A:A | 2.8 |
| rec-111 | G:A | G:G | A:A | G:G | A:A | G:G | C:C | T:T | G:G | A:A | A:A | A:A | A:A | 3.2 |
| rec-112 | G:A | G:G | A:A | G:G | A:A | G:G | C:C | T:T | G:G | A:A | A:A | A:A | A:A | 3.5 |
| rec-113 | G:A | G:G | A:A | G:G | A:A | G:G | C:C | T:T | G:G | A:A | A:A | A:A | A:A | 3.3 |
| rec-114 | G:A | G:G | A:A | G:G | A:A | G:G | C:C | T:T | G:G | A:A | A:A | A:A | A:A | 3.5 |
| rec-115 | G:A | G:G | A:A | G:G | A:A | G:G | C:C | T:T | G:G | A:A | A:A | A:A | A:A | 3 |
| rec-116 | G:A | G:G | A:A | G:G | A:A | G:G | C:C | T:T | G:G | A:A | A:A | A:A | A:A | 2.4 |
| rec-117 | G:A | G:G | A:A | G:G | A:A | G:G | C:C | T:T | G:G | A:A | A:A | A:A | A:A | 3.5 |
| rec-118 | G:A | G:G | A:A | G:G | A:A | G:G | C:C | T:T | G:G | A:A | A:A | A:A | A:A | 2.9 |
| rec-119 | G:A | G:G | A:A | G:G | A:A | G:G | C:C | T:T | G:G | A:A | A:A | A:A | A:A | 2.8 |
| rec-120 | G:A | G:G | A:A | G:G | A:A | G:G | C:C | T:T | G:G | A:A | A:A | A:A | A:A | 2.3 |
| rec-121 | G:A | G:G | A:A | G:G | A:A | G:G | C:C | T:T | G:G | A:A | A:A | A:A | A:A | 3.6 |
| rec-122 | G:A | G:G | A:A | G:G | A:A | G:G | C:C | T:T | G:G | A:A | A:A | A:A | A:A | 2.8 |
| rec-123 | G:A | G:G | A:A | G:G | A:A | G:G | C:C | T:T | G:G | A:A | A:A | A:A | A:A | 3.2 |
| rec-124 | G:A | G:G | A:A | G:G | A:A | G:G | C:C | T:T | G:G | A:A | A:A | A:A | A:A | 3.3 |
| rec-125 | G:A | G:G | A:A | G:G | A:A | G:G | C:C | T:T | G:G | A:A | A:A | A:A | A:A | 2.4 |
| rec-126 | G:A | G:G | A:A | G:G | A:A | G:G | C:C | T:T | G:G | A:A | A:A | A:A | A:A | 3.5 |
| rec-127 | G:A | G:G | A:A | G:G | A:A | G:G | C:C | T:T | G:G | A:A | A:A | A:A | A:A | 3.5 |
| rec-128 | G:A | G:G | A:A | G:G | A:A | G:G | C:C | T:T | G:G | A:A | A:A | A:A | A:A | 3 |
| rec-129 | G:A | G:G | A:A | G:G | A:A | G:G | C:C | T:T | G:G | A:A | A:A | A:A | A:A | 3.1 |

**Table S6. X1132x-derived lines and Huangzaosi-improved lines used to screen the 4-bp InDel alleles.**

| Group | Line name | Genotype | Bred and provided by |
|---|---|---|---|
| X1132x-derived lines | 91227 | TCTT:TCTT | Northwest Agriculture and Forestry University |
| | D9H | / : / | Maize Research Center, Beijing Academy of Agriculture and Forestry Sciences |
| | D9B | / : / | Maize Research Center, Beijing Academy of Agriculture and Forestry Sciences |
| | DH382 | / : / | Shandong Denghai Seeds Co., Ltd. |
| | Jing4055 (京4055) | / : / | Maize Research Center, Beijing Academy of Agriculture and Forestry Sciences |
| | Jing464 (京464) | TCTT:TCTT | Maize Research Center, Beijing Academy of Agriculture and Forestry Sciences |
| | Jing724 (京724) | TCTT:TCTT | Maize Research Center, Beijing Academy of Agriculture and Forestry Sciences |
| | Jing72464 (京72464) | TCTT:TCTT | Maize Research Center, Beijing Academy of Agriculture and Forestry Sciences |
| | Jing725 (京725) | TCTT:TCTT | Maize Research Center, Beijing Academy of Agriculture and Forestry Sciences |
| | Jing88 (京88) | / : / | Maize Research Center, Beijing Academy of Agriculture and Forestry Sciences |
| | B547 | / : / | Maize Research Center, Beijing Academy of Agriculture and Forestry Sciences |
| | MC01 | / : / | Maize Research Center, Beijing Academy of Agriculture and Forestry Sciences |
| | JingX220 (京X220) | TCTT:TCTT | Maize Research Center, Beijing Academy of Agriculture and Forestry Sciences |
| | YuA9241 (豫A9241) | TCTT:TCTT | Henan Agricultural University |
| Huangzaosi-improved lines | Jing2416 (京2416) | TCTT:TCTT | Maize Research Center, Beijing Academy of Agriculture and Forestry Sciences |
| | Jing92 (京92) | TCTT:TCTT | Maize Research Center, Beijing Academy of Agriculture and Forestry Sciences |
| | Chang7-2 (昌7-2) | TCTT:TCTT | Anyang Academy of Agricultural Sciences |
| | Lx9801 | TCTT:TCTT | Maize Institute,Shandong Academy of Agricultural Sciences |

**Table S7. The current hybrid varieties certificated and cultivated in China used in this study.**

| Variety Name | Maternal parent | Paternal parent | Genotype | Approval Number | Bred and provided by |
|---|---|---|---|---|---|
| Xianyu335 (先玉335) | PH6WC | PH4CV | /:TCTT | Guoshenyu2004017 | Tieling Pioneer Seed Research Co., Ltd |
| Denghai605 (登海605) | DH351 | DH382 | /:TCTT | Guoshenyu2010009 | Shandong Denghai Seeds Co., Ltd. |
| Dafeng30 (大丰30) | A311 | PH4CV | /:TCTT | Jinshenyu2012007 | Shanxi Dafeng Seed Co., Ltd. |
| Jingnongke728 (京农科728) | MC01 | Jing2416 | /:TCTT | Guoshenyu20170007 | Maize Research Center, Beijing Academy of Agriculture and Forestry Sciences |
| Jingke528 (京科528) | 90110-2 (D9H) | J2437 | /:TCTT | Jingshenyu2008008 | Maize Research Center, Beijing Academy of Agriculture and Forestry Sciences |
| Denghai685 (登海685) | DH382 | DH357-14 | /:TCTT | Guoshenyu2015011 | Shandong Denghai Seeds Co.,Ltd. |
| Denghai662 (登海662) | DH371 | DH382 | /:TCTT | Guoshenyu2009010 | Shandong Denghai Seeds Co.,Ltd. |
| Jingdan38 (京单38) | D9B | Jing2416 | /:TCTT | Jingshenyu2009005 | Maize Research Center, Beijing Academy of Agriculture and Forestry Sciences |
| Zhengdan958 (郑单958) | Zheng58 | Chang7-2 | TCTT:TCTT | Guoshenyu20000009 | Cereal Crops Institute, Henan Academy of Agricultural Sciences |
| Xundan20 (浚单20) | 9058 | Xun92-8 | TCTT:TCTT | Guoshenyu2003054 | Agricultural Scientific Research Institute of Xunxian in Henan |
| Jingke968 (京科968) | Jing724 | Jing92 | TCTT:TCTT | Guoshenyu2011007 | Maize Research Center, Beijing Academy of Agriculture and Forestry Sciences |
| Demeiya 1 (德美亚1号) | KWS10×KWS73 | KWS49 | TCTT:TCTT | Jishenyu2012048 | KWS SAAT SE & Co. KGaA |
| Nongda108 (农大108) | 178 | Huang C | TCTT:TCTT | Guoshenyu2001002 | China Agricultural University |
| Zhongke11 (中科11号) | CT03 | CT201 | TCTT:TCTT | Guoshenyu2006034 | Beijing Zhongkehuatai Technology Co., Ltd/Henan Ketai Seeds Co.,Ltd. |
| Liyu16 (蠡玉16) | 953 | 91158 | TCTT:TCTT | Sushenyu2003001 | Shijiazhuang Liyu Sci ＆Tech Development Co., Ltd |
| Longping206 (隆平206) | L239 | L7221 | TCTT:TCTT | Wanpinshen07050572 | Anhui Longping Gaoke Seed Industry Co., Ltd. |
| Weike702 (伟科702) | WK858 | WK798-2 | TCTT:TCTT | Guoshenyu2012010 | Zhengzhou Weike Crop Breeding Technology Co., Ltd.<br>Henan Jinyuan SeedIndustry Co., Ltd. |
| Liyu35 (蠡玉35) | L5895 | 912 | TCTT:TCTT | Liaoshenyu2008377 | Shijiazhuang Liyu Sci＆Tech Development Co., Ltd |
| Zhongdan909 (中单909) | Zheng58 | HD568 | TCTT:TCTT | Guoshenyu2011011 | Institute of Crop Sciences, Chinese Academy of Agricultural Sciences |
| Xiangyu998 (翔玉998) | Y822 | X923-1 | TCTT:TCTT | Jishenyu2014038 | Jilin Hongxiang Agriculture Group Hongxiang Seed Industry Co., Ltd |
| Jidan27 (吉单27) | Si-287 | Si-144 | TCTT:TCTT | Jishenyu2002009 | Jinong Gaoxin North Crop Variety Development Center |
| Nonghua101 (农华101) | NH60 | S121 | TCTT:TCTT | Guoshenyu2010008 | Beijing Jinse Nonghua Seed S&T Co., Ltd. |
| Liangyu99 (良玉99) | M03 | M5972 | TCTT:TCTT | Guoshenyu2012008 | Dandong Denghai Liangyu Seed Industry Co., Ltd. |
| Yuyu22 (豫玉22号) | Zong3 | Yu78-1 | TCTT:TCTT | Guoshenyu20000012 | Henan Agricultural University |
| Danyu405 (丹玉405) | Dan299 | DanM9-2 | TCTT:TCTT | Liaoshenyu2008399 | Dandong Academy of Agriculture Sciences |
| Damin3307 (大民3307) | R37 | P2 | TCTT:TCTT | Heishenyu2011009 | Inner Mongolia Damin Seed Co. Ltd. |
| Denghai618 | 521 | DH392 | TCTT:TCTT | Guoshenyu20176113 | Shandong Denghai Seeds Co.,Ltd. |
| Xianyu696 | PH6WC | PHB1M | TCTT:TCTT | Guoshenyu2006025 | Tieling Pioneer Seed Research Company Ltd |
| Zhongdan808 (中单808) | CL11 | NG5 | TCTT:TCTT | Guoshenyu2006037 | Institute of Crop Sciences, Chinese Academy of Agricultural Sciences |
| Nongda372 (农大372) | X24621 | BA702 | TCTT:TCTT | Guoshenyu2015014 | Beijing Nongkeyu Breeding Development Co.,Ltd |
| DK517 (迪卡517) | D1798Z | HCL645 | TCTT:TCTT | Guoshenyu20170005 | Monsanto Far East Limited Beijing Representative Office/China Seed Group Co.,Ltd |
| Jingke665 (京科665) | Jing725 | Jing92 | TCTT:TCTT | Guoshenyu2013003 | Maize Research Center, Beijing Academy of Agriculture and Forestry Sciences |
| NK718 | Jing464 | Jing2416 | TCTT:TCTT | Mengshenyu2011003 | Maize Research Center, Beijing Academy of Agriculture and Forestry Sciences<br>Beijing Nongkeyuan Seed Industry Technology Co.,Ltd |
| MC4592 | Jing4055 | Jing92 | /:TCTT | Jingshenyu2014001 | Maize Research Center, Beijing Academy of Agriculture and Forestry Sciences |

| | | | | | |
|---|---|---|---|---|---|
| MC812 | B547 | Jing2416 | /:TCTT | Jingshenyu2015003 | Maize Research Center, Beijing Academy of Agriculture and Forestry Sciences<br>Shunxin Seed Industry Company |
| Jingdan28 (京单28) | Zheng58 | Jing024 | TCTT:TCTT | Guoshenyu2007001 | Maize Research Center, Beijing Academy of Agriculture and Forestry Sciences |
| Shaandan609 (陕单609) | 91227 | Chang7-2 | TCTT:TCTT | Guoshenyu20016001 | Northwest A&F University |
| Yudan606 (豫单606) | YuA9241 | YuA3 | TCTT:TCTT | Guoshenyu2015018 | Hennan Agricultural University |
| NK971 | Jing388 | Jing372 | TCTT:TCTT | Guoshenyu2014016 | Maize Research Center, Beijing Academy of Agriculture and Forestry Sciences |
| MC220 | X220 | C632 | TCTT:TCTT | Guoshenyu2013018 | Maize Research Center, Beijing Academy of Agriculture and Forestry Sciences |
| Jiandan68 (京单68) | CH8 | Jing2416 | TCTT:TCTT | Guoshenyu2010003 | Maize Research Center, Beijing Academy of Agriculture and Forestry Sciences |
| Jingdan58 (京单58) | CH3 | Jing2416 | TCTT:TCTT | Guoshenyu2010004 | Maize Research Center, Beijing Academy of Agriculture and Forestry Sciences |
| Jingke389 (京科389) | MC03 | Jing2416 | TCTT:TCTT | Guoshenyu2009001 | Maize Research Center, Beijing Academy of Agriculture and Forestry Sciences |
| Jingke308 (京科308) | JN15 | J24-2 | TCTT:TCTT | Guoshenyu2006001 | Maize Research Center, Beijing Academy of Agriculture and Forestry Sciences |
| MC4592 | Jing4055 | Jing92 | /:TCTT | Jingshenyu2014001 | Maize Research Center, Beijing Academy of Agriculture and Forestry Sciences |

**Table S8. Maize landraces, natural population and teosintes used for genotyping *ZmSOS1* gene in this study.**

| Group | Name[a] | Genetype | Provided kindly by |
|---|---|---|---|
| teosinte (*n* = 52) | teosinte-1 | TCTT:TCTT | Prof. Jianbing Yan of Huazhong Agricultural University |
| | teosinte-2 | TCTT:TCTT | |
| | teosinte-3 | TCTT:TCTT | |
| | teosinte-4 | TCTT:TCTT | |
| | teosinte-5 | TCTT:TCTT | |
| | teosinte-6 | TCTT:TCTT | |
| | teosinte-7 | TCTT:TCTT | |
| | teosinte-8 | TCTT:TCTT | |
| | teosinte-9 | TCTT:TCTT | |
| | teosinte-10 | TCTT:TCTT | |
| | teosinte-11 | TCTT:TCTT | |
| | teosinte-12 | TCTT:TCTT | |
| | teosinte-13 | TCTT:TCTT | |
| | teosinte-14 | TCTT:TCTT | |
| | teosinte-15 | TCTT:TCTT | |
| | teosinte-16 | TCTT:TCTT | |
| | teosinte-17 | TCTT:TCTT | |
| | teosinte-18 | TCTT:TCTT | |
| | teosinte-19 | TCTT:TCTT | |
| | teosinte-20 | TCTT:TCTT | |
| | teosinte-21 | TCTT:TCTT | |
| | teosinte-22 | TCTT:TCTT | |
| | teosinte-23 | TCTT:TCTT | |
| | teosinte-24 | TCTT:TCTT | Prof. Xiaohong Yang of China Agricultural University |
| | teosinte-25 | TCTT:TCTT | |
| | teosinte-26 | TCTT:TCTT | |
| | teosinte-27 | TCTT:TCTT | |
| | teosinte-28 | TCTT:TCTT | |
| | teosinte-29 | TCTT:TCTT | |
| | teosinte-30 | TCTT:TCTT | |
| | teosinte-31 | TCTT:TCTT | |
| | teosinte-32 | TCTT:TCTT | |
| | teosinte-33 | TCTT:TCTT | |
| | teosinte-34 | TCTT:TCTT | |
| | teosinte-35 | TCTT:TCTT | |
| | teosinte-36 | TCTT:TCTT | |
| | teosinte-37 | TCTT:TCTT | |
| | teosinte-38 | TCTT:TCTT | |
| | teosinte-39 | TCTT:TCTT | |
| | teosinte-40 | TCTT:TCTT | |
| | teosinte-41 | TCTT:TCTT | |
| | teosinte-42 | TCTT:TCTT | |
| | teosinte-43 | TCTT:TCTT | |
| | teosinte-44 | TCTT:TCTT | |
| | teosinte-45 | TCTT:TCTT | |
| | teosinte-46 | TCTT:TCTT | |
| | teosinte-47 | TCTT:TCTT | |
| | teosinte-48 | TCTT:TCTT | |
| | teosinte-49 | TCTT:TCTT | |
| | teosinte-50 | TCTT:TCTT | |
| | teosinte-51 | TCTT:TCTT | |
| | teosinte-52 | TCTT:TCTT | |

| | | | |
|---|---|---|---|
| landrance ($n$ = 133) | BGC1 | TCTT:TCTT | Prof. Tianyu Wang of Institute of Crop Sciences, Chinese Academy of Agricultural Sciences |
| | Na1110H20c4 | TCTT:TCTT | |
| | Aiqipi (矮七匹) | TCTT:TCTT | |
| | Aiyumi (矮玉米) | TCTT:TCTT | |
| | Bahangbai (八行白) | TCTT:TCTT | |
| | Baibaomi (白包米) | TCTT:TCTT | |
| | Baihenian (白鹤粘) | TCTT:TCTT | |
| | Baihuoyumi (白火玉米) | TCTT:TCTT | |
| | Bailuhuang (白露黄) | TCTT:TCTT | |
| | Baimaya (白马牙) | TCTT:TCTT | |
| | Bainuoyumi (白糯玉米) | TCTT:TCTT | |
| | Bairangziyumi (白穰子玉米) | TCTT:TCTT | |
| | Baitoushuang (白头霜) | TCTT:TCTT | |
| | Baiyugu (白玉谷) | TCTT:TCTT | |
| | Baiyumi (白玉米) | TCTT:TCTT | |
| | Baiyumizibai (白玉米籽白) | TCTT:TCTT | |
| | Bairizao (百日早) | TCTT:TCTT | |
| | Bendibaibaogu (本地白包谷) | TCTT:TCTT | |
| | Bendihuangyumi (本地黄玉米) | TCTT:TCTT | |
| | Bendiyumi (本地玉米) | TCTT:TCTT | |
| | Chuanshazinuo (川沙紫糯) | TCTT:TCTT | |
| | Cibangzi (刺棒子) | TCTT:TCTT | |
| | Dahongguzi (大红骨子) | TCTT:TCTT | |
| | Dahongpao (大红袍) | TCTT:TCTT | |
| | Dajinding (大金顶) | TCTT:TCTT | |
| | Dalihuang (大粒黄) | TCTT:TCTT | |
| | Dapigukuai (大屁股快) | TCTT:TCTT | |
| | Daxinghuang (大兴黄) | TCTT:TCTT | |
| | Dayangbaiyumi (大洋白玉米) | TCTT:TCTT | |
| | Dayumi (大玉米) | TCTT:TCTT | |
| | Denglonghong (灯笼红) | TCTT:TCTT | |
| | Duishengye (对生叶) | TCTT:TCTT | |
| | Erfucao (二伏糙) | TCTT:TCTT | |
| | Erhuangyumi (二黄玉米) | TCTT:TCTT | |
| | Erminziyumi (二民子玉米) | TCTT:TCTT | |
| | Gaoyouyumi (高油玉米) | TCTT:TCTT | |
| | Gaoyouzonghezhong (高油综合种) | TCTT:TCTT | |
| | Guogongzao (郭公早) | TCTT:TCTT | |
| | Hailihuang1 (海里黄) | TCTT:TCTT | |
| | Hailihuang2 (海粒黄) | TCTT:TCTT | |
| | Heihenongjiazhong (黑河农家种) | TCTT:TCTT | |
| | Heisenianyumi (黑色粘玉米) | TCTT:TCTT | |
| | Hongbangzi (红棒子) | TCTT:TCTT | |
| | Hongbaogu (红苞谷) | TCTT:TCTT | |
| | Hongduosuiyumi (红多穗玉米) | TCTT:TCTT | |
| | Honggouzi (红沟子) | TCTT:TCTT | |
| | Hongxinyumi (红心玉米) | TCTT:TCTT | |
| | Huabaogu (花包谷) | TCTT:TCTT | |
| | Huanuoyumi (花糯玉米) | TCTT:TCTT | |
| | Huangbatang (黄八趟) | TCTT:TCTT | |
| | Huangbaogu (黄包谷) | TCTT:TCTT | |
| | Huangbaosu (黄包粟) | TCTT:TCTT | |
| | Huangbaomi (黄苞米) | TCTT:TCTT | |
| | Huangdamayazi (黄大马牙子) | TCTT:TCTT | |
| | Huangfansu (黄番粟) | TCTT:TCTT | |
| | Huanglizi (黄粒子) | TCTT:TCTT | |
| | Huangmaya (黄马牙) | TCTT:TCTT | |

| | | |
|---|---|---|
| Huangnuo (黄糯) | TCTT:TCTT | |
| Huangnuoyumi (黄糯玉米) | TCTT:TCTT | |
| Huangwuyeer (黄五叶儿) | TCTT:TCTT | |
| Huangyingzi (黄硬子) | TCTT:TCTT | |
| Huangyuding (黄玉顶) | TCTT:TCTT | |
| Huangnianyumi (黄粘玉米) | TCTT:TCTT | |
| Jixianbaimaya (蓟县白马牙(混）) | TCTT:TCTT | |
| Jianmingbai (建明白) | TCTT:TCTT | |
| Jinhuanghou (金皇后) | TCTT:TCTT | |
| Jiufengyumi (九峰玉米) | TCTT:TCTT | |
| Junliangchengbaimaya (军粮城白马牙) | TCTT:TCTT | |
| Kuaihongting (快红挺) | TCTT:TCTT | |
| Laolaibie (老来瘪) | TCTT:TCTT | |
| Laorenya (老人牙) | TCTT:TCTT | |
| Liuyuexian (六月鲜) | TCTT:TCTT | |
| Maizibaogu (麦子包谷) | TCTT:TCTT | |
| Maobaogu (毛苞谷) | TCTT:TCTT | |
| Milahuang (蜜腊黄) | TCTT:TCTT | |
| Niuchiyumi (牛齿玉米) | TCTT:TCTT | |
| Pudacunbaogu (普达村包谷) | TCTT:TCTT | |
| Qingpilan (青皮烂) | TCTT:TCTT | |
| Qiubaogu (秋苞谷) | TCTT:TCTT | |
| Ruanyumi (软玉米) | TCTT:TCTT | |
| Taipingandongmaya11 (太平安东马牙11号) | TCTT:TCTT | |
| Wushe (无舌) | TCTT:TCTT | |
| Xiangyabaibaogu (象牙白苞谷) | TCTT:TCTT | |
| Xiaobatang (小八趟) | TCTT:TCTT | |
| Xiaobaici (小白磁) | TCTT:TCTT | |
| Xiaobaigai (小白盖) | TCTT:TCTT | |
| Xiaobaiyumi (小白玉米) | TCTT:TCTT | |
| Xiaobaiyuzi (小白玉籽) | TCTT:TCTT | |
| Xiaohuangyumi (小黄玉米) | TCTT:TCTT | |
| Xiaolihuang (小粒黄) | TCTT:TCTT | |
| Xiaoriqiyumi (小日期玉米) | TCTT:TCTT | |
| Yangyumi (洋玉米) | TCTT:TCTT | |
| Yidaliheiyumi (意大利黑玉米) | TCTT:TCTT | |
| Yinglazi (英粒子) | TCTT:TCTT | |
| Yumi (玉米) | TCTT:TCTT | |
| Zahonggu (杂红骨) | TCTT:TCTT | |
| Zaoshouhuang (早熟黄) | TCTT:TCTT | |
| Zhangshierbaizi (张市二白子) | TCTT:TCTT | |
| Changsuibai (长穗白) | TCTT:TCTT | |
| Zhouluzao (周鹿早) | TCTT:TCTT | |
| landrace-1 | TCTT:TCTT | Prof. Xiaohong Yang of China Agricultural University |
| landrace-2 | TCTT:TCTT | |
| landrace-3 | TCTT:TCTT | |
| landrace-4 | TCTT:TCTT | |
| landrace-5 | TCTT:TCTT | |
| landrace-6 | TCTT:TCTT | |
| landrace-7 | TCTT:TCTT | |
| landrace-8 | TCTT:TCTT | |
| landrace-9 | TCTT:TCTT | |
| landrace-10 | TCTT:TCTT | |
| landrace-11 | TCTT:TCTT | |
| landrace-12 | TCTT:TCTT | |
| landrace-13 | TCTT:TCTT | |
| landrace-14 | TCTT:TCTT | |

| | | | |
|---|---|---|---|
| | landrace-15 | TCTT:TCTT | |
| | landrace-16 | TCTT:TCTT | |
| | landrace-17 | TCTT:TCTT | |
| | landrace-18 | TCTT:TCTT | |
| | landrace-19 | TCTT:TCTT | |
| | landrace-20 | TCTT:TCTT | |
| | landrace-21 | TCTT:TCTT | |
| | landrace-22 | TCTT:TCTT | |
| | landrace-23 | TCTT:TCTT | |
| | landrace-24 | TCTT:TCTT | |
| | landrace-25 | TCTT:TCTT | |
| | landrace-26 | TCTT:TCTT | |
| | landrace-27 | TCTT:TCTT | |
| | landrace-28 | TCTT:TCTT | |
| | landrace-29 | TCTT:TCTT | |
| | landrace-30 | TCTT:TCTT | |
| | landrace-31 | TCTT:TCTT | |
| | landrace-32 | TCTT:TCTT | |
| | landrace-33 | TCTT:TCTT | |
| GWAS panel ($n$ = 489) | 150 | TCTT:TCTT | Prof. Jianbing Yan of Huazhong Agricultural University |
| | 177 | TCTT:TCTT | |
| | 238 | TCTT:TCTT | |
| | 268 | TCTT:TCTT | |
| | 501 | TCTT:TCTT | |
| | 647 | TCTT:TCTT | |
| | 812 | TCTT:TCTT | |
| | 832 | TCTT:TCTT | |
| | 1323 | TCTT:TCTT | |
| | 1462 | TCTT:TCTT | |
| | 3411 | TCTT:TCTT | |
| | 4019 | TCTT:TCTT | |
| | 5213 | TCTT:TCTT | |
| | 5237 | TCTT:TCTT | |
| | 5311 | TCTT:TCTT | |
| | 7327 | TCTT:TCTT | |
| | 7381 | TCTT:TCTT | |
| | 8902 | TCTT:TCTT | |
| | 9642 | TCTT:TCTT | |
| | 9782 | TCTT:TCTT | |
| | 81162 | TCTT:TCTT | |
| | 526018 | TCTT:TCTT | |
| | 04K5672 | TCTT:TCTT | |
| | 04K5686 | TCTT:TCTT | |
| | 04K5702 | TCTT:TCTT | |
| | 05W002 | TCTT:TCTT | |
| | 05WN230 | TCTT:TCTT | |
| | 07KS4 | TCTT:TCTT | |
| | 18-599 | TCTT:TCTT | |
| | 303WX | TCTT:TCTT | |
| | 384-2 | TCTT:TCTT | |
| | 3H-2 | TCTT:TCTT | |
| | 4F1 | TCTT:TCTT | |
| | 835A | TCTT:TCTT | |
| | 835B | TCTT:TCTT | |
| | 975-12 | TCTT:TCTT | |
| | A619 | TCTT:TCTT | |

| | |
|---|---|
| B11 | TCTT:TCTT |
| B110 | TCTT:TCTT |
| B111 | TCTT:TCTT |
| B113 | TCTT:TCTT |
| B114 | TCTT:TCTT |
| B151 | TCTT:TCTT |
| B73 | TCTT:TCTT |
| B77 | TCTT:TCTT |
| BEM | TCTT:TCTT |
| BGY | TCTT:TCTT |
| BS16 | TCTT:TCTT |
| BT1 | TCTT:TCTT |
| BY4839 | TCTT:TCTT |
| BY4944 | TCTT:TCTT |
| BY4960 | TCTT:TCTT |
| BY804 | TCTT:TCTT |
| BY807 | TCTT:TCTT |
| BY809 | TCTT:TCTT |
| BY813 | TCTT:TCTT |
| BY815 | TCTT:TCTT |
| BY843 | TCTT:TCTT |
| BY855 | TCTT:TCTT |
| C8605 | TCTT:TCTT |
| CF3 | TCTT:TCTT |
| CHANG3 | TCTT:TCTT |
| CHANG7-2 | TCTT:TCTT |
| CHENG698 | TCTT:TCTT |
| CHUAN48-2 | TCTT:TCTT |
| CI7 | TCTT:TCTT |
| CIMBL1 | TCTT:TCTT |
| CIMBL10 | TCTT:TCTT |
| CIMBL101 | TCTT:TCTT |
| CIMBL102 | TCTT:TCTT |
| CIMBL104 | TCTT:TCTT |
| CIMBL105 | TCTT:TCTT |
| CIMBL106 | TCTT:TCTT |
| CIMBL107 | TCTT:TCTT |
| CIMBL108 | TCTT:TCTT |
| CIMBL109 | TCTT:TCTT |
| CIMBL11 | TCTT:TCTT |
| CIMBL110 | TCTT:TCTT |
| CIMBL111 | TCTT:TCTT |
| CIMBL112 | TCTT:TCTT |
| CIMBL113 | TCTT:TCTT |
| CIMBL114 | TCTT:TCTT |
| CIMBL115 | TCTT:TCTT |
| CIMBL116 | TCTT:TCTT |
| CIMBL117 | TCTT:TCTT |
| CIMBL118 | TCTT:TCTT |
| CIMBL119 | TCTT:TCTT |
| CIMBL12 | TCTT:TCTT |
| CIMBL120 | TCTT:TCTT |
| CIMBL121 | TCTT:TCTT |
| CIMBL124 | TCTT:TCTT |
| CIMBL125 | TCTT:TCTT |
| CIMBL126 | TCTT:TCTT |
| CIMBL127 | TCTT:TCTT |

| | |
|---|---|
| CIMBL128 | TCTT:TCTT |
| CIMBL129 | TCTT:TCTT |
| CIMBL13 | TCTT:TCTT |
| CIMBL130 | TCTT:TCTT |
| CIMBL131 | TCTT:TCTT |
| CIMBL132 | TCTT:TCTT |
| CIMBL133 | TCTT:TCTT |
| CIMBL134 | TCTT:TCTT |
| CIMBL135 | TCTT:TCTT |
| CIMBL136 | TCTT:TCTT |
| CIMBL138 | TCTT:TCTT |
| CIMBL139 | TCTT:TCTT |
| CIMBL14 | TCTT:TCTT |
| CIMBL140 | TCTT:TCTT |
| CIMBL141 | TCTT:TCTT |
| CIMBL142 | TCTT:TCTT |
| CIMBL143 | TCTT:TCTT |
| CIMBL144 | TCTT:TCTT |
| CIMBL145 | TCTT:TCTT |
| CIMBL146 | TCTT:TCTT |
| CIMBL147 | TCTT:TCTT |
| CIMBL148 | TCTT:TCTT |
| CIMBL149 | TCTT:TCTT |
| CIMBL15 | TCTT:TCTT |
| CIMBL150 | TCTT:TCTT |
| CIMBL151 | TCTT:TCTT |
| CIMBL152 | TCTT:TCTT |
| CIMBL153 | TCTT:TCTT |
| CIMBL154 | TCTT:TCTT |
| CIMBL155 | TCTT:TCTT |
| CIMBL156 | TCTT:TCTT |
| CIMBL157 | TCTT:TCTT |
| CIMBL16 | TCTT:TCTT |
| CIMBL17 | TCTT:TCTT |
| CIMBL18 | TCTT:TCTT |
| CIMBL19 | TCTT:TCTT |
| CIMBL2 | TCTT:TCTT |
| CIMBL20 | TCTT:TCTT |
| CIMBL21 | TCTT:TCTT |
| CIMBL22 | TCTT:TCTT |
| CIMBL23 | TCTT:TCTT |
| CIMBL24 | TCTT:TCTT |
| CIMBL25 | TCTT:TCTT |
| CIMBL26 | TCTT:TCTT |
| CIMBL27 | TCTT:TCTT |
| CIMBL28 | TCTT:TCTT |
| CIMBL3 | TCTT:TCTT |
| CIMBL30 | TCTT:TCTT |
| CIMBL31 | TCTT:TCTT |
| CIMBL32 | TCTT:TCTT |
| CIMBL33 | TCTT:TCTT |
| CIMBL34 | TCTT:TCTT |
| CIMBL35 | TCTT:TCTT |
| CIMBL36 | TCTT:TCTT |
| CIMBL37 | TCTT:TCTT |
| CIMBL38 | TCTT:TCTT |
| CIMBL39 | TCTT:TCTT |

| | |
|---|---|
| CIMBL4 | TCTT:TCTT |
| CIMBL40 | TCTT:TCTT |
| CIMBL41 | TCTT:TCTT |
| CIMBL42 | TCTT:TCTT |
| CIMBL44 | TCTT:TCTT |
| CIMBL45 | TCTT:TCTT |
| CIMBL46 | TCTT:TCTT |
| CIMBL47 | TCTT:TCTT |
| CIMBL48 | TCTT:TCTT |
| CIMBL49 | TCTT:TCTT |
| CIMBL5 | TCTT:TCTT |
| CIMBL50 | /:/ |
| CIMBL51 | TCTT:TCTT |
| CIMBL52 | TCTT:TCTT |
| CIMBL53 | TCTT:TCTT |
| CIMBL54 | TCTT:TCTT |
| CIMBL55 | TCTT:TCTT |
| CIMBL56 | TCTT:TCTT |
| CIMBL57 | TCTT:TCTT |
| CIMBL58 | TCTT:TCTT |
| CIMBL59 | TCTT:TCTT |
| CIMBL6 | TCTT:TCTT |
| CIMBL60 | TCTT:TCTT |
| CIMBL61 | TCTT:TCTT |
| CIMBL62 | TCTT:TCTT |
| CIMBL63 | TCTT:TCTT |
| CIMBL65 | TCTT:TCTT |
| CIMBL67 | TCTT:TCTT |
| CIMBL68 | TCTT:TCTT |
| CIMBL69 | TCTT:TCTT |
| CIMBL7 | TCTT:TCTT |
| CIMBL70 | TCTT:TCTT |
| CIMBL71 | TCTT:TCTT |
| CIMBL72 | TCTT:TCTT |
| CIMBL73 | TCTT:TCTT |
| CIMBL74 | TCTT:TCTT |
| CIMBL75 | TCTT:TCTT |
| CIMBL77 | TCTT:TCTT |
| CIMBL78 | TCTT:TCTT |
| CIMBL79 | TCTT:TCTT |
| CIMBL8 | TCTT:TCTT |
| CIMBL80 | TCTT:TCTT |
| CIMBL81 | TCTT:TCTT |
| CIMBL82 | TCTT:TCTT |
| CIMBL83 | TCTT:TCTT |
| CIMBL84 | /:/ |
| CIMBL85 | TCTT:TCTT |
| CIMBL86 | TCTT:TCTT |
| CIMBL88 | TCTT:TCTT |
| CIMBL89 | TCTT:TCTT |
| CIMBL90 | TCTT:TCTT |
| CIMBL91 | TCTT:TCTT |
| CIMBL92 | TCTT:TCTT |
| CIMBL93 | TCTT:TCTT |
| CIMBL94 | TCTT:TCTT |
| CIMBL95 | TCTT:TCTT |
| CIMBL96 | TCTT:TCTT |

| | |
|---|---|
| CIMBL97 | TCTT:TCTT |
| CIMBL98 | TCTT:TCTT |
| CIMBL99 | TCTT:TCTT |
| CML113 | TCTT:TCTT |
| CML114 | TCTT:TCTT |
| CML115 | TCTT:TCTT |
| CML116 | TCTT:TCTT |
| CML118 | TCTT:TCTT |
| CML121 | TCTT:TCTT |
| CML122 | TCTT:TCTT |
| CML130 | TCTT:TCTT |
| CML134 | TCTT:TCTT |
| CML139 | TCTT:TCTT |
| CML162 | TCTT:TCTT |
| CML163 | TCTT:TCTT |
| CML165 | TCTT:TCTT |
| CML166 | TCTT:TCTT |
| CML168 | TCTT:TCTT |
| CML169 | TCTT:TCTT |
| CML170 | TCTT:TCTT |
| CML171 | TCTT:TCTT |
| CML172 | TCTT:TCTT |
| CML189 | TCTT:TCTT |
| CML191 | TCTT:TCTT |
| CML192 | TCTT:TCTT |
| CML20 | TCTT:TCTT |
| CML223 | TCTT:TCTT |
| CML225 | TCTT:TCTT |
| CML226 | TCTT:TCTT |
| CML228 | TCTT:TCTT |
| CML26 | TCTT:TCTT |
| CML27 | TCTT:TCTT |
| CML28 | TCTT:TCTT |
| CML282 | TCTT:TCTT |
| CML285 | TCTT:TCTT |
| CML287 | TCTT:TCTT |
| CML289 | TCTT:TCTT |
| CML29 | TCTT:TCTT |
| CML290 | TCTT:TCTT |
| CML298 | TCTT:TCTT |
| CML300 | TCTT:TCTT |
| CML304 | TCTT:TCTT |
| CML305 | TCTT:TCTT |
| CML307 | TCTT:TCTT |
| CML31 | TCTT:TCTT |
| CML32 | TCTT:TCTT |
| CML323 | TCTT:TCTT |
| CML324 | TCTT:TCTT |
| CML325 | TCTT:TCTT |
| CML326 | TCTT:TCTT |
| CML327 | TCTT:TCTT |
| CML360 | TCTT:TCTT |
| CML361 | TCTT:TCTT |
| CML364 | TCTT:TCTT |
| CML40 | TCTT:TCTT |
| CML408 | TCTT:TCTT |
| CML411 | TCTT:TCTT |

| | |
|---|---|
| CML412 | TCTT:TCTT |
| CML415 | TCTT:TCTT |
| CML422 | TCTT:TCTT |
| CML423 | TCTT:TCTT |
| CML426 | TCTT:TCTT |
| CML430 | TCTT:TCTT |
| CML432 | TCTT:TCTT |
| CML433 | TCTT:TCTT |
| CML451 | TCTT:TCTT |
| CML454 | TCTT:TCTT |
| CML465 | TCTT:TCTT |
| CML468 | TCTT:TCTT |
| CML470 | TCTT:TCTT |
| CML473 | TCTT:TCTT |
| CML474 | TCTT:TCTT |
| CML479 | TCTT:TCTT |
| CML480 | TCTT:TCTT |
| CML486 | TCTT:TCTT |
| CML493 | TCTT:TCTT |
| CML496 | TCTT:TCTT |
| CML497 | TCTT:TCTT |
| CML50 | TCTT:TCTT |
| CML51 | TCTT:TCTT |
| CML69 | TCTT:TCTT |
| CY72 | TCTT:TCTT |
| D047 | TCTT:TCTT |
| D863F | TCTT:TCTT |
| DAN3130 | TCTT:TCTT |
| DAN340 | TCTT:TCTT |
| DAN360 | TCTT:TCTT |
| DAN4245 | TCTT:TCTT |
| DAN598 | TCTT:TCTT |
| DAN599 | TCTT:TCTT |
| DAN9046 | TCTT:TCTT |
| DE.EX | TCTT:TCTT |
| DH29 | TCTT:TCTT |
| DH3732 | TCTT:TCTT |
| DONG237 | TCTT:TCTT |
| DONG46 | TCTT:TCTT |
| DSB | TCTT:TCTT |
| EN25 | TCTT:TCTT |
| ES40 | TCTT:TCTT |
| FCD0602 | TCTT:TCTT |
| GEMS10 | TCTT:TCTT |
| GEMS11 | TCTT:TCTT |
| GEMS12 | TCTT:TCTT |
| GEMS13 | TCTT:TCTT |
| GEMS14 | TCTT:TCTT |
| GEMS15 | TCTT:TCTT |
| GEMS17 | TCTT:TCTT |
| GEMS18 | TCTT:TCTT |
| GEMS19 | TCTT:TCTT |
| GEMS20 | TCTT:TCTT |
| GEMS21 | TCTT:TCTT |
| GEMS23 | TCTT:TCTT |
| GEMS24 | TCTT:TCTT |
| GEMS25 | TCTT:TCTT |

| | |
|---|---|
| GEMS27 | TCTT:TCTT |
| GEMS28 | TCTT:TCTT |
| GEMS29 | TCTT:TCTT |
| GEMS3 | TCTT:TCTT |
| GEMS30 | TCTT:TCTT |
| GEMS31 | TCTT:TCTT |
| GEMS32 | TCTT:TCTT |
| GEMS33 | TCTT:TCTT |
| GEMS35 | TCTT:TCTT |
| GEMS36 | TCTT:TCTT |
| GEMS37 | TCTT:TCTT |
| GEMS39 | TCTT:TCTT |
| GEMS4 | TCTT:TCTT |
| GEMS40 | TCTT:TCTT |
| GEMS41 | TCTT:TCTT |
| GEMS42 | TCTT:TCTT |
| GEMS43 | TCTT:TCTT |
| GEMS44 | TCTT:TCTT |
| GEMS45 | TCTT:TCTT |
| GEMS47 | TCTT:TCTT |
| GEMS48 | TCTT:TCTT |
| GEMS49 | TCTT:TCTT |
| GEMS5 | TCTT:TCTT |
| GEMS50 | TCTT:TCTT |
| GEMS51 | TCTT:TCTT |
| GEMS52 | TCTT:TCTT |
| GEMS53 | TCTT:TCTT |
| GEMS54 | TCTT:TCTT |
| GEMS55 | TCTT:TCTT |
| GEMS56 | TCTT:TCTT |
| GEMS57 | TCTT:TCTT |
| GEMS58 | TCTT:TCTT |
| GEMS59 | TCTT:TCTT |
| GEMS6 | TCTT:TCTT |
| GEMS60 | TCTT:TCTT |
| GEMS61 | TCTT:TCTT |
| GEMS63 | TCTT:TCTT |
| GEMS64 | TCTT:TCTT |
| GEMS9 | TCTT:TCTT |
| GY1032 | TCTT:TCTT |
| GY220 | TCTT:TCTT |
| GY237 | TCTT:TCTT |
| GY246 | TCTT:TCTT |
| GY386 | TCTT:TCTT |
| GY462 | TCTT:TCTT |
| GY798 | TCTT:TCTT |
| GY923 | TCTT:TCTT |
| H21 | TCTT:TCTT |
| HAI1134 | TCTT:TCTT |
| HSBN | TCTT:TCTT |
| HTH-17 | TCTT:TCTT |
| HU803 | TCTT:TCTT |
| HUA83-2 | TCTT:TCTT |
| HUANGC | TCTT:TCTT |
| HYS | TCTT:TCTT |
| HZS | TCTT:TCTT |
| IRF291 | TCTT:TCTT |

| | |
|---|---|
| IRF314 | TCTT:TCTT |
| J4112 | TCTT:TCTT |
| JH59 | TCTT:TCTT |
| JH96C | TCTT:TCTT |
| JI53 | TCTT:TCTT |
| JI63 | TCTT:TCTT |
| JI842 | TCTT:TCTT |
| JI846 | TCTT:TCTT |
| JI853 | TCTT:TCTT |
| JIAO51 | TCTT:TCTT |
| JY01 | TCTT:TCTT |
| K10 | TCTT:TCTT |
| K12 | TCTT:TCTT |
| K14 | TCTT:TCTT |
| K22 | TCTT:TCTT |
| L3180 | TCTT:TCTT |
| LG001 | TCTT:TCTT |
| LIAO138 | TCTT:TCTT |
| LIAO159 | TCTT:TCTT |
| LIAO5114 | TCTT:TCTT |
| LIAO5262 | TCTT:TCTT |
| LIAO5263 | TCTT:TCTT |
| LK11 | TCTT:TCTT |
| LV28 | TCTT:TCTT |
| LX9801 | TCTT:TCTT |
| LXN | TCTT:TCTT |
| LY | TCTT:TCTT |
| LY042 | TCTT:TCTT |
| M153 | TCTT:TCTT |
| M165 | TCTT:TCTT |
| M97 | TCTT:TCTT |
| MN | TCTT:TCTT |
| MO113 | TCTT:TCTT |
| MO17 | TCTT:TCTT |
| NAN21-3 | TCTT:TCTT |
| NMJT | TCTT:TCTT |
| P138 | TCTT:TCTT |
| P178 | TCTT:TCTT |
| Q1261 | TCTT:TCTT |
| QI205 | TCTT:TCTT |
| QI319 | TCTT:TCTT |
| R08 | TCTT:TCTT |
| R15 | TCTT:TCTT |
| SI273 | TCTT:TCTT |
| SI434 | TCTT:TCTT |
| SI444 | TCTT:TCTT |
| SI446 | TCTT:TCTT |
| SK | TCTT:TCTT |
| SW1611 | TCTT:TCTT |
| SW92E114 | TCTT:TCTT |
| SY1032 | TCTT:TCTT |
| SY1035 | TCTT:TCTT |
| SY1039 | TCTT:TCTT |
| SY1052 | TCTT:TCTT |
| SY1077 | TCTT:TCTT |
| SY1128 | TCTT:TCTT |
| SY3073 | TCTT:TCTT |

| | |
|---|---|
| SY998 | TCTT:TCTT |
| SY999 | TCTT:TCTT |
| TIAN77 | TCTT:TCTT |
| TIE7922 | TCTT:TCTT |
| TT16 | TCTT:TCTT |
| TX5 | TCTT:TCTT |
| TY1 | TCTT:TCTT |
| TY10 | TCTT:TCTT |
| TY11 | TCTT:TCTT |
| TY2 | TCTT:TCTT |
| TY3 | TCTT:TCTT |
| TY4 | TCTT:TCTT |
| TY5 | TCTT:TCTT |
| TY6 | TCTT:TCTT |
| TY7 | TCTT:TCTT |
| TY8 | TCTT:TCTT |
| TY9 | TCTT:TCTT |
| W138 | TCTT:TCTT |
| WH413 | TCTT:TCTT |
| WMR | TCTT:TCTT |
| WU109 | TCTT:TCTT |
| X1141P | TCTT:TCTT |
| XI502 | TCTT:TCTT |
| XUN971 | TCTT:TCTT |
| XZ698 | TCTT:TCTT |
| YAN414 | TCTT:TCTT |
| YE107 | TCTT:TCTT |
| YE478 | TCTT:TCTT |
| YE488 | TCTT:TCTT |
| YE515 | TCTT:TCTT |
| YE52106 | TCTT:TCTT |
| YU374 | TCTT:TCTT |
| YU87-1 | TCTT:TCTT |
| YUN46 | TCTT:TCTT |
| YZ15 | TCTT:TCTT |
| Z2018F | TCTT:TCTT |
| ZAC546 | TCTT:TCTT |
| ZB648 | TCTT:TCTT |
| ZHENG22 | TCTT:TCTT |
| ZHENG28 | TCTT:TCTT |
| ZHENG29 | TCTT:TCTT |
| ZHENG30 | TCTT:TCTT |
| ZHENG32 | TCTT:TCTT |
| ZHENG35 | TCTT:TCTT |
| ZHENG58 | TCTT:TCTT |
| ZHENG653 | TCTT:TCTT |
| ZHI41 | TCTT:TCTT |
| ZHONG69 | TCTT:TCTT |
| ZI330 | TCTT:TCTT |
| ZONG3 | TCTT:TCTT |
| ZONG31 | TCTT:TCTT |
| ZZ01 | TCTT:TCTT |
| ZZ03 | TCTT:TCTT |

[a] The names and pedigrees of all teosintes and 33 landraces provided by Prof. Xiaohong Yang were not available. The names of landraces in Chinese were indicated in parentheses.

**Table S9. List of the 55 SNPs used for association analysis of *ZmSOS1* gene.**

| SNPs[a] | Alleles | Codon change | Amino acid change | Position (B73 RefGen_v2) | Position ( in *ZmSOS1* gene)[b] |
|---|---|---|---|---|---|
| chr1.S_180953555 | C/G | 5'-UTR | | Chr1: 180953555 | - 44 |
| chr1.S_180953506 | C/G | GGC/GGG | G/G | Chr1: 180953506 | + 6 |
| chr1.S_180953453 | A/C | CTG/CGG | L/R | Chr1: 180953453 | + 59 |
| chr1.S_180953402 | A/C | GTC/GGC | V/R | Chr1: 180953402 | + 110 |
| chr1.S_180950975 | A/G | GGC/GGT | G/G | Chr1: 180950975 | + 2537 |
| chr1.S_180950973 | A/G | GTT/GCT | V/A | Chr1: 180950973 | + 2539 |
| chr1.S_180950861 | A/T | ACA/ACT | T/T | Chr1: 180950861 | + 2651 |
| chr1.S_180950807 | T/C | GCA/GCG | A/A | Chr1: 180950807 | + 2705 |
| chr1.S_180950792 | A/G | CTC/CTT | L/L | Chr1: 180950792 | + 2720 |
| PZE-101140295 | A/G | intron 5 | | Chr1: 180950030 | + 3482 |
| PZE-101140288 | A/G | intron 9 | | Chr1: 180948029 | + 5483 |
| chr1.S_180947145 | T/G | GGC/GGA | G/G | Chr1: 180947145 | + 6367 |
| PZE-101140284 | A/G | intron 11 | | Chr1: 180946735 | + 6777 |
| chr1.S_180943545 | T/C | GCC/ACC | A/T | Chr1: 180943545 | + 9967 |
| chr1.S_180943510 | A/T | GCT/GCA | A/A | Chr1: 180943510 | + 10002 |
| chr1.S_180943508 | A/C | GTT/GGT | V/G | Chr1: 180943508 | + 10004 |
| chr1.S_180941772 | A/G | TAC/TAT | Y/Y | Chr1: 180941772 | + 11740 |
| chr1.S_180941748 | A/C | GAG/GAT | E/D | Chr1: 180941748 | + 11764 |
| chr1.S_180941652 | T/C | GAG/GAA | E/E | Chr1: 180941652 | + 11860 |
| chr1.S_180939766 | A/G | GCC/GCT | A/A | Chr1: 180939766 | + 13746 |
| chr1.S_180939763 | A/C | GCG/GCT | A/A | Chr1: 180939763 | + 13749 |
| SYN2836 | A/C | ATA/ATC | I/I | Chr1: 180939733 | + 13779 |
| PZE-101140277 | A/T | CGT/CGA | R/R | Chr1: 180939514 | + 13998 |
| chr1.S_180937488 | T/C | AAG/AAA | K/K | Chr1: 180937488 | + 16024 |
| chr1.S_180937459 | A/G | GTC/GCC | V/A | Chr1: 180937459 | + 16053 |
| chr1.S_180937280 | A/G | GTT/GTC | V/V | Chr1: 180937280 | + 16232 |
| chr1.S_180937271 | A/C | CTT/CTC | L/L | Chr1: 180937271 | + 16241 |
| SYN2838 | A/G | GTG/GTC | V/V | Chr1: 180937259 | + 16253 |
| chr1.S_180937183 | C/G | CCA/GCA | P/A | Chr1: 180937183 | + 16329 |
| chr1.S_180937012 | T/C | AGA/AGG | R/R | Chr1: 180937012 | + 16500 |
| chr1.S_180936988 | T/C | TTA/TTG | L/L | Chr1: 180936988 | + 16524 |
| chr1.S_180936979 | T/C | GGG/GGA | G/G | Chr1: 180936979 | + 16533 |
| chr1.S_180936871 | T/C | GAG/GAA | E/E | Chr1: 180936871 | + 16641 |
| chr1.S_180936866 | A/C | CTC/CGC | L/R | Chr1: 180936866 | + 16646 |
| chr1.S_180936677 | C/G | CTG/CTC | L/L | Chr1: 180936677 | + 16835 |
| chr1.S_180936579 | A/G | ATC/ACC | I/T | Chr1: 180936579 | + 16933 |
| chr1.S_180936537 | T/G | CCA/CAA | P/Q | Chr1: 180936537 | + 16975 |
| chr1.S_180936343 | T/C | ATC/GTC | I/V | Chr1: 180936343 | + 17169 |
| chr1.S_180936296 | A/G | AGT/AGC | S/S | Chr1: 180936296 | + 17216 |
| chr1.S_180936201 | T/C | AGC/AAC | S/N | Chr1: 180936201 | + 17311 |
| chr1.S_180936169 | T/C | GGC/AGC | G/S | Chr1: 180936169 | + 17343 |
| chr1.S_180936133 | T/C | GGG/AGG | G/R | Chr1: 180936133 | + 17379 |
| chr1.S_180936092 | A/G | CAC/CAT | H/H | Chr1: 180936092 | + 17420 |
| chr1.S_180936041 | A/G | AAC/AAT | N/N | Chr1: 180936041 | + 17471 |
| chr1.S_180936038 | T/C | CAG/CAA | Q/Q | Chr1: 180936038 | + 17474 |
| chr1.S_180935932 | A/G | CCT/TCT | P/S | Chr1: 180935932 | + 17580 |
| chr1.S_180935922 | T/G | GCT/GAT | A/D | Chr1: 180935922 | + 17590 |
| chr1.S_180935756 | A/G | CCC/CCT | P/P | Chr1: 180935756 | + 17756 |
| chr1.S_180935736 | A/G | 3'-UTR | | Chr1: 180935736 | + 17776 |
| PZE-101140256 | A/G | 3'-UTR | | Chr1: 180935716 | + 17796 |
| chr1.S_180935656 | A/T | 3'-UTR | | Chr1: 180935656 | + 17856 |
| chr1.S_180935650 | T/C | 3'-UTR | | Chr1: 180935650 | + 17862 |
| chr1.S_180935648 | T/C | 3'-UTR | | Chr1: 180935648 | + 17864 |
| chr1.S_180935647 | A/C | 3'-UTR | | Chr1: 180935647 | + 17865 |
| chr1.S_180935613 | A/G | 3'-UTR | | Chr1: 180935613 | + 17899 |

[a] The SNPs were retrieved from the 1.1-million SNPs for 368 GWAS lines (http://www.maizego.org/Resources.html) (Fu *et al*., 2013).

[b] The positions of the SNPs were based on the start codon position of *ZmSOS1* gene (NATG, N = -1 and A = +1).

**Table S10. Phenotypes of shoot $Na^+$ content and root length of GWAS lines under salt stress used for association analysis of *ZmSOS1* gene.**

| Line names | Shoot $Na^+$ content (mg $g^{-1}$ DW)[a] | Root length (cm)[b] | chr1.S _180950792 | chr1.S _180935613 | chr1.S _180936537 | chr1.S _180941748 |
|---|---|---|---|---|---|---|
| 150 | NaN | 3.8 | GG | GG | GG | CC |
| 177 | 9.885 | 3.5 | GG | GG | GG | CC |
| 238 | 1.244 | 4.5 | GG | GG | GG | CC |
| 268 | 12.148 | 4.18 | GG | GG | GG | CC |
| 647 | 1.154 | 4.74 | GG | GG | GG | CC |
| 1462 | 3.882 | 5.4 | GG | GG | GG | CC |
| 4019 | 6.381 | 5.28 | GG | GG | GG | CC |
| 5213 | 8.672 | 3.13 | GG | GG | GG | CC |
| 5237 | 7.353 | 3.02 | GG | GG | GG | CC |
| 7327 | 3.409 | 4.45 | GG | GG | GG | CC |
| 7381 | 10.646 | 4.41 | GG | GG | GG | CC |
| 8902 | 2.180 | NaN | GG | GG | GG | CC |
| 9642 | 3.737 | 4.5 | GG | GG | GG | CC |
| 526018 | 12.328 | 4.47 | GG | GG | GG | CC |
| 05W002 | 2.428 | 5.63 | GG | GG | GG | CC |
| 05WN230 | 4.336 | 6.19 | GG | GG | GG | CC |
| 07KS4 | 3.726 | 4.44 | GG | GG | GG | CC |
| 18-599 | NaN | 5.35 | GG | GG | GG | CC |
| 303WX | 5.926 | 2.78 | GG | GG | GG | CC |
| 7884-4HT | 8.379 | NaN | GG | GG | GG | CC |
| 4F1 | NaN | 5.42 | GG | GG | GG | CC |
| 835A | 8.526 | 4.71 | GG | GG | GG | CC |
| 835B | 11.429 | 4.31 | GG | GG | GG | CC |
| 975-12 | 19.969 | 8.38 | GG | GG | GG | CC |
| B11 | 4.219 | 5.19 | GG | GG | GG | CC |
| B110 | NaN | 4.77 | GG | GG | GG | CC |
| B111 | 16.948 | 3.6 | GG | GG | GG | CC |
| B113 | 7.269 | 3 | GG | GG | GG | CC |
| B114 | 2.082 | 6.29 | GG | GG | GG | CC |
| B151 | 4.970 | 3.68 | GG | GG | GG | CC |
| B73 | 9.536 | 4.84 | GG | GG | GG | CC |
| B77 | 2.608 | 5.84 | GG | GG | GG | CC |
| BS16 | 1.646 | 4.18 | GG | GG | GG | CC |
| BY4839 | 8.648 | 2.45 | GG | GG | GG | CC |
| BY4944 | NaN | 4.17 | GG | GG | GG | CC |
| BY4960 | 7.686 | 5.52 | GG | GG | GG | CC |
| BY804 | 5.270 | 3.08 | GG | GG | GG | CC |
| BY807 | 6.078 | 2.97 | GG | GG | GG | CC |
| BY809 | 7.392 | 3.75 | GG | GG | GG | CC |
| BY813 | 5.707 | 3.8 | GG | GG | GG | CC |
| BY815 | NaN | 2.71 | GG | GG | GG | CC |
| BY855 | 10.705 | 3.76 | GG | GG | GG | CC |
| CHANG3 | 7.242 | 8.43 | GG | GG | GG | CC |
| CHENG698 | 13.269 | 8.43 | GG | GG | GG | CC |
| CHUAN48-2 | 8.369 | 7.86 | GG | GG | GG | CC |
| CI7 | 4.796 | 10.24 | GG | GG | GG | CC |
| CIMBL1 | 3.135 | 7.94 | GG | GG | GG | CC |
| CIMBL10 | 5.743 | NaN | GG | GG | GG | CC |
| CIMBL100 | NaN | 7.83 | GG | GG | GG | CC |
| CIMBL101 | 4.395 | 9.35 | GG | GG | GG | CC |
| CIMBL102 | 4.397 | 10.67 | GG | GG | GG | CC |

| | | | | | | |
|---|---|---|---|---|---|---|
| CIMBL105 | 20.653 | 7.72 | GG | AA | TT | AA |
| CIMBL106 | 6.166 | 6.23 | GG | GG | GG | CC |
| CIMBL108 | NaN | 6.52 | GG | GG | GG | CC |
| CIMBL109 | NaN | 7.15 | GG | GG | GG | CC |
| CIMBL11 | 8.984 | 12.13 | GG | GG | GG | CC |
| CIMBL111 | 6.630 | 9.63 | GG | GG | GG | CC |
| CIMBL113 | NaN | 5.89 | GG | GG | GG | CC |
| CIMBL114 | 9.555 | 6.62 | GG | GG | GG | CC |
| CIMBL115 | 5.201 | 10.93 | GG | GG | GG | CC |
| CIMBL116 | 4.244 | 7.99 | GG | GG | GG | CC |
| CIMBL119 | 5.549 | 6.78 | GG | GG | GG | CC |
| CIMBL12 | 7.616 | 4.24 | GG | GG | GG | CC |
| CIMBL120 | 15.253 | 5.37 | GG | GG | GG | CC |
| CIMBL121 | 5.954 | 5.5 | GG | GG | GG | CC |
| CIMBL122 | 7.271 | 6.11 | GG | GG | GG | CC |
| CIMBL123 | 3.168 | 4.96 | GG | GG | GG | CC |
| CIMBL124 | 9.081 | 4.17 | GG | GG | GG | CC |
| CIMBL125 | 17.960 | 5.35 | GG | GG | GG | CC |
| CIMBL127 | 2.063 | 2.37 | GG | GG | GG | CC |
| CIMBL129 | 0.816 | 3.11 | GG | GG | GG | CC |
| CIMBL13 | 10.469 | 7.46 | GG | GG | GG | CC |
| CIMBL133 | NaN | 4.53 | GG | GG | GG | CC |
| CIMBL139 | 10.523 | 6.43 | GG | GG | GG | CC |
| CIMBL140 | 13.394 | 5.02 | GG | GG | GG | CC |
| CIMBL141 | 6.044 | 6.17 | GG | GG | GG | CC |
| CIMBL142 | 10.307 | 4.59 | GG | GG | GG | CC |
| CIMBL143 | 7.329 | 4.72 | GG | GG | GG | CC |
| CIMBL144 | 4.220 | 4.8 | GG | GG | GG | CC |
| CIMBL145 | 7.795 | 2.92 | GG | GG | GG | CC |
| CIMBL147 | 1.950 | 4.39 | GG | GG | GG | CC |
| CIMBL149 | 11.429 | 4.61 | GG | GG | GG | CC |
| CIMBL15 | 3.447 | 5.45 | GG | GG | GG | CC |
| CIMBL150 | NaN | 5.09 | GG | GG | GG | CC |
| CIMBL151 | 3.685 | 4.84 | GG | GG | GG | CC |
| CIMBL152 | 7.240 | 7.21 | GG | GG | GG | CC |
| CIMBL153 | 6.957 | 5.48 | GG | GG | GG | CC |
| CIMBL156 | NaN | 5.82 | GG | GG | GG | CC |
| CIMBL157 | 12.266 | 2.75 | AA | GG | GG | CC |
| CIMBL16 | 22.203 | 5.17 | GG | GG | GG | CC |
| CIMBL17 | 6.364 | 6.83 | GG | GG | GG | CC |
| CIMBL18 | 9.148 | 4.3 | GG | GG | GG | CC |
| CIMBL19 | 0.588 | 6.18 | GG | GG | GG | CC |
| CIMBL2 | 3.297 | 5.04 | GG | GG | GG | CC |
| CIMBL21 | 5.885 | 3.01 | GG | GG | GG | CC |
| CIMBL22 | 4.739 | 4.79 | GG | GG | GG | CC |
| CIMBL23 | 8.554 | 3.9 | GG | GG | GG | CC |
| CIMBL25 | 12.559 | 5.71 | GG | GG | GG | CC |
| CIMBL27 | 3.874 | 4.02 | GG | GG | GG | CC |
| CIMBL28 | 5.076 | 3.6 | GG | GG | GG | CC |
| CIMBL29 | 2.467 | 4.25 | GG | GG | GG | CC |
| CIMBL3 | 5.685 | NaN | GG | GG | GG | CC |
| CIMBL32 | 4.357 | 5.06 | GG | GG | GG | CC |
| CIMBL38 | 9.443 | 3.65 | GG | GG | GG | CC |
| CIMBL4 | NaN | 7.07 | GG | GG | GG | CC |
| CIMBL40 | 6.436 | 5.13 | GG | GG | GG | CC |
| CIMBL42 | 3.911 | 6.06 | GG | GG | GG | CC |

| | | | | | | |
|---|---|---|---|---|---|---|
| CIMBL43 | 2.547 | 8.5 | GG | GG | GG | CC |
| CIMBL46 | 9.415 | 6.91 | GG | GG | GG | CC |
| CIMBL47 | 5.254 | 8.6 | GG | GG | GG | CC |
| CIMBL48 | 13.982 | 5.97 | GG | GG | GG | CC |
| CIMBL49 | 5.208 | 8.79 | GG | GG | GG | CC |
| CIMBL5 | 10.836 | 8.05 | GG | GG | GG | CC |
| CIMBL50 | NaN | 4.81 | GG | GG | GG | CC |
| CIMBL51 | 7.428 | 8.61 | GG | GG | GG | CC |
| CIMBL52 | 7.122 | 5.11 | GG | GG | GG | CC |
| CIMBL53 | 8.641 | 7.05 | GG | GG | GG | CC |
| CIMBL54 | 3.159 | 4.68 | GG | GG | GG | CC |
| CIMBL55 | 12.712 | 7.67 | GG | GG | GG | CC |
| CIMBL56 | 6.357 | 5.79 | GG | GG | GG | CC |
| CIMBL58 | 6.539 | 6.61 | GG | GG | GG | CC |
| CIMBL59 | NaN | 8.14 | GG | GG | GG | CC |
| CIMBL6 | 5.988 | 5.15 | GG | AA | GG | CC |
| CIMBL60 | 3.350 | 5.66 | GG | GG | GG | CC |
| CIMBL62 | 6.771 | 5.33 | GG | GG | GG | CC |
| CIMBL63 | 10.469 | 5.67 | GG | GG | GG | CC |
| CIMBL66 | NaN | 5.21 | GG | GG | GG | CC |
| CIMBL68 | 8.250 | 3.21 | GG | GG | GG | CC |
| CIMBL69 | 4.347 | 5.81 | GG | GG | GG | CC |
| CIMBL7 | 4.165 | 6.12 | GG | GG | GG | CC |
| CIMBL70 | 0.939 | 4.4 | GG | GG | GG | CC |
| CIMBL71 | 10.035 | 6.24 | GG | GG | GG | CC |
| CIMBL74 | NaN | 5.98 | GG | GG | GG | CC |
| CIMBL75 | NaN | 6.98 | GG | GG | GG | CC |
| CIMBL77 | 4.446 | 4.67 | GG | GG | GG | CC |
| CIMBL79 | 3.491 | 4.07 | GG | GG | GG | CC |
| CIMBL81 | 10.363 | 6.79 | GG | GG | GG | CC |
| CIMBL82 | 7.312 | 4.93 | GG | GG | GG | CC |
| CIMBL83 | 5.730 | 4.02 | GG | GG | GG | CC |
| CIMBL84 | 12.380 | 4.01 | GG | GG | GG | CC |
| CIMBL86 | 3.382 | 7.57 | GG | GG | GG | CC |
| CIMBL87 | 6.657 | NaN | GG | GG | GG | CC |
| CIMBL88 | 2.946 | 9.32 | GG | GG | GG | CC |
| CIMBL89 | 5.438 | 6.44 | GG | GG | GG | CC |
| CIMBL9 | 9.727 | 4.55 | GG | GG | GG | CC |
| CIMBL90 | 12.626 | 3.87 | GG | GG | GG | CC |
| CIMBL91 | NaN | 5.01 | GG | GG | GG | CC |
| CIMBL92 | NaN | 8.2 | GG | GG | GG | CC |
| CIMBL93 | 4.655 | 4.65 | GG | GG | GG | CC |
| CIMBL94 | 6.044 | 5.42 | GG | GG | GG | CC |
| CIMBL95 | 1.544 | 5.55 | GG | GG | GG | CC |
| CIMBL96 | 7.457 | 5.05 | GG | GG | GG | CC |
| CIMBL98 | 5.687 | 5.51 | GG | GG | GG | CC |
| CIMBL99 | 8.196 | 8.3 | GG | GG | GG | CC |
| CML114 | 5.275 | 8.37 | GG | GG | GG | CC |
| CML115 | 4.851 | 6.66 | GG | GG | GG | CC |
| CML116 | NaN | 3.23 | GG | GG | GG | CC |
| CML118 | 3.922 | 4.97 | GG | GG | GG | CC |
| CML121 | 1.639 | 5.03 | GG | GG | GG | CC |
| CML122 | 7.332 | 4.38 | GG | GG | GG | CC |
| CML130 | 4.862 | 2.97 | GG | GG | GG | CC |
| CML134 | 7.012 | 6.08 | GG | GG | GG | CC |
| CML139 | NaN | 5.24 | GG | GG | GG | CC |

| | | | | | | |
|---|---|---|---|---|---|---|
| CML162 | 11.416 | 3.33 | GG | GG | GG | CC |
| CML163 | 7.812 | 5.77 | GG | GG | GG | CC |
| CML165 | NaN | 5.16 | GG | GG | GG | CC |
| CML169 | 3.113 | 3.78 | GG | GG | GG | CC |
| CML170 | 10.104 | 4.69 | GG | GG | GG | CC |
| CML171 | 5.416 | 6.34 | GG | GG | GG | CC |
| CML172 | 3.685 | 5.36 | GG | GG | GG | CC |
| CML189 | 2.986 | NaN | GG | GG | GG | CC |
| CML191 | 4.353 | 4.49 | GG | GG | GG | CC |
| CML192 | 4.156 | 2.95 | GG | GG | GG | CC |
| CML20 | 8.383 | 3.89 | GG | GG | GG | CC |
| CML290 | 4.505 | 3.15 | GG | GG | GG | CC |
| CML298 | 4.248 | 4.86 | GG | GG | GG | CC |
| CML304 | 12.893 | 2.48 | GG | GG | GG | CC |
| CML31 | 9.566 | 4.56 | GG | GG | GG | CC |
| CML32 | 3.364 | 4.46 | GG | GG | GG | CC |
| CML323 | NaN | 2.98 | GG | GG | GG | CC |
| CML324 | NaN | 4.54 | GG | GG | GG | CC |
| CML325 | NaN | 4.97 | GG | GG | GG | CC |
| CML327 | 12.152 | 4.77 | GG | GG | GG | CC |
| CML360 | 28.433 | 4.74 | GG | AA | GG | CC |
| CML361 | NaN | 3.84 | GG | GG | GG | CC |
| CML411 | 1.989 | 5.07 | GG | GG | GG | CC |
| CML415 | 13.293 | 3.86 | GG | GG | GG | CC |
| CML422 | NaN | 3.81 | GG | GG | GG | CC |
| CML423 | NaN | 5.4 | GG | GG | GG | CC |
| CML426 | NaN | 5.35 | GG | GG | GG | CC |
| CML431 | 2.541 | NaN | GG | GG | GG | CC |
| CML432 | 7.459 | 5.23 | GG | GG | GG | CC |
| CML433 | 1.287 | 5.7 | GG | GG | GG | CC |
| CML454 | 4.213 | 5.28 | GG | GG | GG | CC |
| CML470 | 10.665 | 4.02 | GG | GG | GG | CC |
| CML479 | 8.268 | 6.45 | GG | GG | GG | CC |
| CML480 | 2.714 | 4.41 | GG | GG | GG | CC |
| CML486 | 12.780 | 5.87 | GG | GG | GG | CC |
| CML493 | 15.395 | NaN | GG | GG | GG | CC |
| CML496 | 3.414 | 2.71 | GG | GG | GG | CC |
| CML50 | 4.891 | 6.44 | GG | GG | GG | CC |
| CML69 | NaN | 8 | GG | GG | GG | CC |
| D863F | 8.140 | 3.43 | GG | GG | GG | CC |
| DAN3130 | 7.360 | 9.45 | GG | GG | GG | CC |
| DAN340 | 1.397 | 6.54 | GG | GG | GG | CC |
| DAN360 | 1.793 | 11.04 | GG | GG | GG | CC |
| DAN4245 | 2.903 | 6.83 | GG | GG | GG | CC |
| DAN599 | 10.788 | 7.03 | GG | GG | GG | CC |
| DH3732 | 13.689 | 9.65 | GG | GG | GG | CC |
| DONG237 | NaN | 5.73 | GG | GG | GG | CC |
| DONG46 | NaN | 7.06 | GG | GG | GG | CC |
| EN25 | 0.846 | 5.43 | GG | GG | GG | CC |
| ES40 | 2.491 | 4.37 | GG | GG | GG | CC |
| FCD0602 | 1.543 | 6.27 | GG | GG | GG | CC |
| GEMS1 | 7.851 | 5.83 | GG | GG | GG | CC |
| GEMS10 | 7.176 | 8 | GG | GG | GG | CC |
| GEMS11 | 2.305 | 6.94 | GG | GG | GG | CC |
| GEMS13 | NaN | 5.01 | GG | GG | GG | CC |
| GEMS14 | 7.343 | 4.93 | GG | GG | GG | CC |

| | | | | | | |
|---|---|---|---|---|---|---|
| GEMS15 | NaN | 6.23 | GG | GG | GG | CC |
| GEMS16 | 4.200 | 5.62 | GG | GG | GG | CC |
| GEMS17 | 5.779 | 6.16 | GG | GG | GG | CC |
| GEMS18 | 6.686 | 3.7 | GG | GG | GG | CC |
| GEMS19 | 4.977 | 4.85 | GG | GG | GG | CC |
| GEMS2 | 4.696 | 5.25 | GG | GG | GG | CC |
| GEMS20 | NaN | 4.2 | GG | GG | GG | CC |
| GEMS21 | 8.160 | 3.89 | GG | GG | GG | CC |
| GEMS23 | 7.981 | 7.52 | GG | GG | GG | CC |
| GEMS25 | 13.957 | 4.83 | GG | GG | GG | CC |
| GEMS28 | 4.467 | 5.94 | GG | GG | GG | CC |
| GEMS29 | 7.983 | 5.23 | GG | GG | GG | CC |
| GEMS3 | 14.255 | 6.6 | GG | GG | GG | CC |
| GEMS30 | 4.387 | 4.66 | GG | GG | GG | CC |
| GEMS31 | 5.217 | 5.54 | GG | GG | GG | CC |
| GEMS32 | 6.445 | 4.94 | GG | GG | GG | CC |
| GEMS33 | 1.568 | 6.54 | GG | GG | GG | CC |
| GEMS35 | 11.039 | 6.02 | GG | AA | TT | AA |
| GEMS36 | 5.088 | 9.88 | GG | GG | GG | CC |
| GEMS37 | 2.197 | 10.22 | GG | AA | TT | AA |
| GEMS39 | 4.411 | 6.61 | GG | GG | GG | CC |
| GEMS4 | 8.876 | 5.76 | GG | GG | GG | CC |
| GEMS40 | 2.311 | 5.24 | GG | GG | GG | CC |
| GEMS41 | 1.176 | 2.78 | AA | GG | GG | CC |
| GEMS42 | 3.330 | 2.9 | AA | GG | GG | CC |
| GEMS44 | 4.240 | 8.06 | GG | GG | GG | CC |
| GEMS46 | 3.228 | NaN | GG | GG | GG | CC |
| GEMS48 | 0.844 | 8.61 | GG | GG | GG | CC |
| GEMS49 | 3.853 | 12.93 | GG | GG | GG | CC |
| GEMS5 | 1.823 | 8.99 | GG | GG | GG | CC |
| GEMS50 | NaN | 8.8 | GG | GG | GG | CC |
| GEMS51 | 5.417 | 12.56 | GG | GG | GG | CC |
| GEMS54 | 7.216 | 9.19 | GG | GG | GG | CC |
| GEMS55 | 1.302 | 10.95 | GG | GG | GG | CC |
| GEMS56 | NaN | 8.45 | GG | GG | GG | CC |
| GEMS58 | 3.680 | 6.88 | GG | GG | GG | CC |
| GEMS59 | 1.672 | 5.14 | GG | GG | GG | CC |
| GEMS6 | NaN | 4.02 | GG | GG | GG | CC |
| GEMS60 | 2.302 | 4.28 | GG | GG | GG | CC |
| GEMS61 | 10.057 | 5.82 | GG | GG | GG | CC |
| GEMS62 | 6.550 | 3.14 | GG | GG | GG | CC |
| GEMS63 | NaN | 6.07 | GG | GG | GG | CC |
| GEMS64 | 14.059 | 3.95 | GG | GG | GG | CC |
| GEMS65 | NaN | 6.45 | GG | GG | GG | CC |
| GEMS66 | 7.557 | 6.6 | GG | GG | GG | CC |
| GEMS9 | 4.035 | 5.26 | GG | GG | GG | CC |
| GY1007 | 4.149 | NaN | GG | GG | GG | CC |
| GY1032 | 5.126 | 6.73 | GG | GG | GG | CC |
| GY386 | NaN | 5.04 | GG | GG | GG | CC |
| GY462 | NaN | 4.22 | GG | GG | GG | CC |
| GY798 | 9.138 | 8.7 | GG | GG | GG | CC |
| GY923 | 6.688 | 7.25 | GG | GG | GG | CC |
| HTH-17 | NaN | 10.94 | GG | GG | GG | CC |
| HUA83-2 | 4.700 | 10.72 | GG | GG | GG | CC |
| HYS | 5.671 | 8.75 | GG | GG | GG | CC |
| HZS | 3.417 | 4.72 | GG | GG | GG | CC |

| | | | | | | |
|---|---|---|---|---|---|---|
| IRF314 | 2.654 | 9.8 | GG | GG | GG | CC |
| J4112 | 0.580 | 9.48 | GG | GG | GG | CC |
| JH59 | 0.494 | 6.87 | GG | GG | GG | CC |
| JH96C | 1.623 | 10.15 | GG | GG | GG | CC |
| JI63 | NaN | 3.77 | GG | GG | GG | CC |
| JI842 | NaN | 5.12 | GG | GG | GG | CC |
| JI846 | 6.315 | 8.65 | GG | GG | GG | CC |
| JI853 | 5.532 | 5.48 | GG | GG | GG | CC |
| JIAO51 | 16.520 | 5.4 | GG | GG | GG | CC |
| JY01 | 9.898 | 5.62 | GG | GG | GG | CC |
| K10 | 6.200 | 5.48 | GG | GG | GG | CC |
| K12 | 3.910 | 7.6 | GG | GG | GG | CC |
| K14 | NaN | 5.88 | GG | GG | GG | CC |
| K22 | 3.373 | 5.18 | GG | GG | GG | CC |
| L3180 | 2.539 | 6.22 | GG | GG | GG | CC |
| LG001 | 11.775 | 4.46 | GG | GG | GG | CC |
| LIAO138 | 7.561 | 7.02 | GG | GG | GG | CC |
| LIAO159 | 1.746 | 7.06 | GG | GG | GG | CC |
| LIAO5114 | 2.912 | 4.27 | GG | GG | GG | CC |
| LIAO5262 | 1.044 | 5.95 | GG | GG | GG | CC |
| LIAO5263 | 4.991 | 5.99 | GG | GG | GG | CC |
| LK11 | 4.032 | 5.85 | GG | GG | GG | CC |
| LV28 | 1.109 | 5.3 | GG | GG | GG | CC |
| LX9801 | NaN | 4.49 | GG | GG | GG | CC |
| LXN | 18.802 | 10.6 | GG | GG | GG | CC |
| LY042 | NaN | 8.19 | GG | GG | GG | CC |
| M153 | 3.286 | 10.46 | GG | GG | GG | CC |
| M97 | 2.968 | 7.99 | GG | GG | GG | CC |
| MO113 | 4.681 | 5.22 | GG | GG | GG | CC |
| MO17 | 3.186 | 7.89 | GG | GG | GG | CC |
| NAN21-3 | 7.225 | 11.6 | GG | GG | GG | CC |
| P178 | 5.720 | 7.05 | GG | GG | GG | CC |
| Q1261 | 3.615 | 8.93 | GG | GG | GG | CC |
| QI205 | 1.261 | 8.89 | GG | GG | GG | CC |
| R15 | 2.132 | 6.33 | GG | GG | GG | CC |
| R15X1141 | 1.706 | 5.09 | GG | GG | GG | CC |
| RY713 | NaN | 5.5 | GG | GG | GG | CC |
| RY729 | 3.954 | 4.75 | GG | GG | GG | CC |
| S22 | 1.172 | 5.31 | GG | GG | GG | CC |
| SC55 | NaN | 6.63 | GG | GG | GG | CC |
| SHEN5003 | 2.598 | NaN | GG | GG | GG | CC |
| SI273 | 4.311 | 3.27 | GG | GG | GG | CC |
| SI434 | NaN | 3.4 | GG | GG | GG | CC |
| SI446 | 4.179 | 2.13 | GG | GG | GG | CC |
| SW92E114 | 2.920 | 5.07 | GG | GG | GG | CC |
| SY1032 | NaN | 6.4 | GG | GG | GG | CC |
| SY1035 | 4.327 | 5.09 | GG | GG | GG | CC |
| SY1039 | 1.370 | 4.02 | GG | GG | GG | CC |
| SY1052 | NaN | 5.65 | GG | GG | GG | CC |
| SY1128 | 4.139 | 5.71 | GG | GG | GG | CC |
| SY3073 | 2.259 | 4.82 | GG | GG | GG | CC |
| TIAN77 | 6.653 | 4.99 | GG | GG | GG | CC |
| TIE7922 | 0.571 | 5.59 | GG | GG | GG | CC |
| TY1 | 2.462 | 7.73 | GG | GG | GG | CC |
| TY11 | 4.339 | 9.22 | GG | GG | GG | CC |
| TY2 | NaN | 5.16 | GG | GG | GG | CC |

| | | | | | | |
|---|---|---|---|---|---|---|
| TY3 | 6.109 | 6.27 | GG | GG | GG | CC |
| TY4 | 2.954 | 6.29 | GG | GG | GG | CC |
| TY5 | 4.133 | 4.52 | GG | GG | GG | CC |
| TY6 | 7.367 | 5.77 | GG | GG | GG | CC |
| U8112 | 0.807 | 8.45 | GG | GG | GG | CC |
| W138 | 0.334 | 9.15 | GG | GG | GG | CC |
| WH413 | NaN | 9.7 | GG | GG | GG | CC |
| WU109 | NaN | 9.4 | GG | GG | GG | CC |
| XI502 | 2.657 | 6.21 | GG | GG | GG | CC |
| XUN971 | 0.335 | 9.13 | GG | GG | GG | CC |
| XZ698 | 4.798 | NaN | GG | GG | GG | CC |
| YE478 | 0.572 | NaN | GG | GG | GG | CC |
| YE515 | 5.643 | 4.51 | GG | GG | GG | CC |
| YE52106 | 3.698 | 6.23 | GG | GG | GG | CC |
| YE8001 | 2.791 | 6.7 | GG | GG | GG | CC |
| YU374 | 8.023 | 5.31 | GG | GG | GG | CC |
| Z2018F | 12.972 | 6.98 | GG | GG | GG | CC |
| ZAC546 | 2.758 | 3.33 | GG | GG | GG | CC |
| ZB648 | 13.773 | 3.6 | GG | GG | GG | CC |
| ZH68 | 1.748 | NaN | GG | GG | GG | CC |
| ZHENG28 | 3.363 | 6.24 | GG | GG | GG | CC |
| ZHENG29 | 0.450 | 7.78 | GG | GG | GG | CC |
| ZHENG30 | 14.362 | 2.8 | GG | GG | GG | CC |
| ZHENG32 | 1.969 | 4.93 | GG | GG | GG | CC |
| ZHENG35 | 1.350 | 4.69 | GG | GG | GG | CC |
| ZHENG653 | 1.287 | 4.7 | GG | GG | GG | CC |
| ZHI41 | 1.609 | 4.19 | GG | GG | GG | CC |
| ZHONG69 | 4.434 | 3.18 | GG | GG | GG | CC |
| ZONG31 | 3.894 | 5.12 | GG | GG | GG | CC |
| ZZ01 | 1.240 | 4.38 | GG | GG | GG | CC |
| ZZ03 | 10.348 | NaN | GG | GG | GG | CC |

[a] Shoot $Na^+$ content data of 304 inbred lines under salt stress were obtained from Cao *et al*. (2020) DW, Dry weight.

[b] Root length data of 347 inbred lines under salt stress were obtained from Luo *et al.* (2021)

**Table S11. The PCR primers used in this study.**

| Primer name | Sequence (5'->3') | Usage |
|---|---|---|
| 1F | TTGTGGACATGCGACGTGAG | RT-PCR analysis of *AC186524.3_FG001* |
| 1R | CACCACTGACATCACGACCA | |
| 2F | TGTTCGAGTTACATTCCCGCA | RT-PCR analysis of *GRMZM2G098494* |
| 2R | GGTTGGCCTGGATCCTTCTC | |
| 3F | GATACGGGTAAGAAATCTCGCC | RT-PCR analysis of *GRMZM2G399359* |
| 3R | TGTTCATGTGCACCCTCTGT | |
| 4F | ACCACTCAAAGAACACCCCA | RT-PCR analysis of *GRMZM2G399367* |
| 4R | TTCCATCCACCTTGCCATCC | |
| 5F | GGTTTATTTGGTTGGGTAGGGC | RT-PCR analysis of *GRMZM2G541900* |
| 5R | CGAATAGGAGAGGTGAACTGCA | |
| 6F | TCATGGTGCCAAGGTCTTTATG | RT-PCR analysis of *AC186524.3_FG005* |
| 6R | TCCGACTTAGCATGCCACTT | |
| 7F | TCTCCTTGGCCTTGATGACG | RT-PCR analysis of *AC186524.3_FG006* |
| 7R | CCCTTCCATACTCCTCCCCA | |
| Actin1F | CCTGACACTGAAGTACCCGA | RT-PCR analysis of *Actin1* , *GRMZM2G126010* , as an internal contronl gene |
| Actin1R | CAGTCTCCAGCTCCTGTTCA | |
| P41F | CATCGCCTGTTACTTGCTGC | PCR and sequencing analysis of *ZmSOS1* genomic DNA, promoter |
| P41R | ACGGACAACACTTCTCACCC | |
| P34F | GATGGCGCTAGGAGATTGCT | |
| P34R | CCATCTCGCTGGTCCTTGTA | |
| P42F | GCAACCTGTTACCCGCTAGA | PCR and sequencing analysis of *ZmSOS1* genomic DNA, gene body |
| P42R | TCCGTGGCATCTACTTCAACC | |
| P18F | GTGCTCTTCGTTGGGGTGTC | |
| P18R | TGGGGTTTAGATGGCTGGCT | |
| P19F | TGCTTGCCATTACCAGCCAG | |
| P19R | TTTGCAGCAGTCCCTAAGCG | |
| P20F | TGCCTCCAGAACATCCCAGA | |
| P20R | CCCTGGCTGACTCAACCTCT | |
| P21F | ACAGGACCGCTATTGTTGCC | |
| P21R | TCCCATGTGTACAGAAGAGCT | |
| P22F | AGAACTCCTGCCGGTCCTTT | |
| P22R | AGCAACCATTTCCCTGAGTACA | |
| P23F | CGACCTATTTGCACCTGGGG | |
| P23R | ACGACATAGAGGGTGCGACA | |
| P24F | TGTCGCACCCTCTATGTCGT | |
| P24R | TGAAACTCCGAGGACACCGA | |
| P25F | TCGGTGTCCTCGGAGTTTCA | |
| P25R | CTGCTGTCACGTGCAAAGGA | |
| P26F | TCCTTTGCACGTGACAGCAG | |
| P26R | TCTGGGGTTTGGGGTGAAGT | |
| P33F | ACTTCACCCCAAACCCCAGA | |
| P33R | GTTGGTCTGCTCTGCCCATC | |
| P43F | GGAACAGATGGGGAGCCAGA | |
| P43R | CCCCGTACATGCTCAGTTGC | |
| M13F | GTAAAACGACGGCCAGT | Cloning and sequencing analysis of *ZmSOS1* cDNA using the pEASY Blunt Simple vector (TransGen Biotech., Beijing, China) |
| M13R | CAGGAAACAGCTATGAC | |
| 11F | CCAAGGGGTCGTCACATACA | |
| 17905R | ACACCACACCACACCACTG | |
| 9899F | GGGGCTTCCTTCTTCTGCTC | |
| 9985R | CAAAGTGACGCAACATCGGG | |
| 2F | TGTTCGAGTTACATTCCCGCA | |
| 2R | GGTTGGCCTGGATCCTTCTC | |
| 5-ROP | GCTGATGGCGATGAATGAACACTG | 5'-RACE primer, outer primer for nest PCR |
| 5-369ROP | TCCAAGCTTGCCTAGACCATGT | 5'-RACE primer, outer primer fornest PCR |
| 5-RIP | GAACACTGCGTTTGCTGGCTTTGATG | 5'-RACE primer, inner primer for nest PCR |
| 5-129RIP | AGAGCAAGCGACGAAGAACAGAGT | 5'-RACE primer, inner primer for nest PCR |
| 3-3239F | CCACGACGATGACAAACCAG | 3'-RACE primer, outer primer for nest PCR |
| 3-ROP | GCGAGCACAGAATTAATACGACT | 3'-RACE primer, outer primer for nest PCR |
| 3-3364F | CCTGAGGTTGCTGCCACCGC | 3'-RACE primer, inner primer for nest PCR |
| 3-RIP | CGCGGATCCGAATTAATACGACTCACTATAGG | 3'-RACE primer, inner primer for nest PCR |

| | | |
|---|---|---|
| SOS1qF1<br>SOS1qR1 (= P43R) | GCCGCCCATCTTCATGTTCA<br>CCCCGTACATGCTCAGTTGC | For qRT-PCR analysis of *ZmSOS1* gene in response to salt stress |
| ACTqF3<br>ACTqR3 | CCCCAAGGCCAACAGAGAGA<br>GCTCACACCATCACCGGAAT | For qRT-PCR analysis, the internal contronl gene, *Actin1* , *GRMZM2G126010* |
| X-SOS1GFP-F1<br>X-SOS1GFP-R1 | AGCAGATCTATCGATTCTAGAAAGGGGTCGTCACATACAAGG<br>TCCTTTGCCCATGGCTCTAGAGGACTCACGGAAAGAGAGCAT | For pM999-ZmSOS1[Jing724]-eGFP construction using *Xba* I site. The adapter sequence for homologous recombination was highlighted in |
| SOS1-F1<br>SOS1-T7-R1 | GCAACTGAGCATGTACGGG<br>CATTAATACGACTCACTATAGGGAGAGAAGACCACATCCAGGC | Antisense probes synthesis *in vitro* for in situ hybridization detection of *ZmSOS1* gene (T7 promoter sequence was underlined) |
| SOS1-T7-F1<br>SOS1-R1 | CATTAATACGACTCACTATAGGGGCAACTGAGCATGTACGGG<br>AGAGAAGACCACATCCAGGC | Sense probes synthesis *in vitro* for in situ hybridization detection of *ZmSOS1* gene |
| 2F<br>2R | TGTTCGAGTTACATTCCCGCA<br>GGTTGGCCTGGATCCTTCTC | For PCR analysis of EMS mutant *zmsos1-1* allele. |
| 296F<br>296R | AGCTCTTCTGTACACATGGGA<br>GGAGTCAACCACCCAGTCGG | For PCR analysis of EMS mutant *zmsos1-2* allele. |
| 296F2<br>296R2 | ACTGCTAAAAGAACTCGGAGCA<br>CCCATACCCAAAGTGACGCA | For identification of alternatively spliced transcripts in *zmsos1-1* mutant. |
| S-11F1<br>S-17905R1 | AAACGCACTAGTATCCCGGGCCAAGGGGTCGTCACATACA<br>TTATGGCGCGCCTTCCCGGGACACCACACCACACCACTG | Constrution of the overexpression vector pZZ-ZmSOS1 using *Sma* I site. (Ubiquitin promoter-ZmSOS1[Jing724]-NOS terminitor) |
| B-11F<br>B-17905R | CAGGTCGACTCTAGAGGATCCCAAGGGGTCGTCACATACA<br>GAGCTCGGTACCCGGGGATCCACACCACACCACACCACTG | Constrution of the overexpression vector pMRC35-ZmSOS1 using *Bam* HI site. (CaMV 35S promoter-ZmSOS1[Jing724]-PolyA terminitor) |
| atsos1-1F<br>atsos1-1R | GCATTGGTCTGGCGTTTGG<br>AGCACAGGAATTCAAGGTCTACA | For PCR detection of *atsos1-1* mutation allele. |
| 2F<br>2R | TGTTCGAGTTACATTCCCGCA<br>GGTTGGCCTGGATCCTTCTC | For PCR detection of *ZmSOS1* [Jing724] CDS in *Arabidopsis* transformant |
| 35S-F<br>35S-R | GCCTCTTCGCTATTACGCCA<br>GGGCTCATGGTAGACTCGA | For PCR detection of CaMV 35S promoter in *Arabidopsis* transformai |

**Table S12. Construction of yeast expression vectors for *ZmSOS1*, *AtSOS2* and *AtSOS3* genes.**

| Vectors of interest | PCR templates | Primer Name | Primer sequence (5'-to-3') | Inserted into (Enzyme site) | Expression cassettes |
|---|---|---|---|---|---|
| pDR196-ZmSOS1 | Jing724 seedling cDNA | S-11F2 | TATACCCCAGCCTCGACTAGTCCAAGGGGTCGTCACATACA | pDR196 (*Spe* I/*XhoI*) | PMA promoter-ZmSOS1-ADH1 terminator |
| | | X-17905R2 | GGTACCGGGCCCCCCCTCGAGACACCACACCACACCACTG | | |
| pDR196-AtSOS2 | Col-0 seedling cDNA | S-180F | TATACCCCAGCCTCGACTAGTACGCCTCTTTCATCAACCCT | | PMA promoter-AtSOS2-ADH1 terminator |
| | | X-1657R | GGTACCGGGCCCCCCCTCGAGAGCAAATCTTCCAATCTCCCTGA | | |
| pDR196-AtSOS3 | | S-419F | TATACCCCAGCCTCGACTAGTGGTGTGTTTGTATGGGCTGC | | PMA promoter-AtSOS3-ADH1 terminator |
| | | X-1114R | GGTACCGGGCCCCCCCTCGAGCCTTGCATGGCTTATATTAGGAAGA | | |
| p404MET25-AtSOS2 | Col-0 seedling cDNA | S-180F2 | CATCCATACTCTAGAACTAGTACGCCTCTTTCATCAACCCT | p404MET25 (Addgene, Plasmid #17420) (*Spe* I/*Xho* I) | MEF17 promoter-AtSOS2-CYC1 terminator |
| | | X-1657R2 | TAACTAATTACATGACTCGAGCAAATCTTCCAATCTCCCTGA | | |
| p404MET25-AtSOS3 | | M419F | CATCCATACTCTAGAACTAGTGGTGTGTTTGTATGGGCTGC | | MEF17 promoter-AtSOS3-CYC1 terminator |
| | | M1114R | TAACTAATTACATGACTCGAGCCTTGCATGGCTTATATTAGGAAGA | | |
| pDR196-ZmSOS1-AtSOS2 | p404MET25-AtSOS2 plasmid DNA | KM13/pUC_R | GCCTGAATGGCGAATGGCGCCAGCGGATAACAATTTCACACAGG | pDR196-ZmSOS1 (*Kas* I) | PMA promoter-ZmSOS1-ADH1 terminator + MEF17 promoter-AtSOS2-CYC1 terminator |
| | | KM13Fplus | GAAAATACCGCATCAGGCGCCTGTAAAACGACGGCCAGTGA | | |
| pDR196-ZmSOS1-AtSOS3 | p404MET25-AtSOS3 plasmid DNA | KM13/pUC_R | GCCTGAATGGCGAATGGCGCCAGCGGATAACAATTTCACACAGG | | PMA promoter-ZmSOS1-ADH1 terminator + MEF17 promoter-AtSOS3-CYC1 terminator |
| | | KM13Fplus | GAAAATACCGCATCAGGCGCCTGTAAAACGACGGCCAGTGA | | |
| pDR196-AtSOS2-AtSOS3 | | KM13/pUC_R | GCCTGAATGGCGAATGGCGCCAGCGGATAACAATTTCACACAGG | pDR196-AtSOS2 (*Kas* I) | PMA promoter-AtSOS2-ADH1 terminator + MEF17 promoter-AtSOS3-CYC1 terminator |
| | | KM13Fplus | GAAAATACCGCATCAGGCGCCTGTAAAACGACGGCCAGTGA | | |
